# IntS6 and the Integrator phosphatase module tune the efficiency of select premature transcription termination events

**DOI:** 10.1101/2023.03.05.531184

**Authors:** Rina Fujiwara, Si-Nan Zhai, Dongming Liang, Matthew Tracey, Xu-Kai Ma, Aayushi P. Shah, Christopher J. Fields, María Saraí Mendoza-Figueroa, Michele C. Meline, Deirdre C. Tatomer, Li Yang, Jeremy E. Wilusz

## Abstract

The metazoan-specific Integrator complex catalyzes 3’ end processing of small nuclear RNAs (snRNAs) as well as premature transcription termination events that attenuate expression of many protein-coding genes. Integrator has RNA endonuclease and protein phosphatase activities, but it remains unclear if both are always required for complex function. Here, we show that IntS6 (Integrator subunit 6) over-expression is sufficient to block Integrator function at a subset of *Drosophila* protein-coding genes, while having no effect on snRNA processing or attenuation of other loci. Over-expressed IntS6 titrates protein phosphatase 2A (PP2A) subunits from the rest of the Integrator complex and thus the only loci affected are those where the phosphatase module is necessary for Integrator function. IntS6 functions analogous to a PP2A regulatory B subunit as over-expression of canonical B subunits, which do not associate with Integrator, are also sufficient to inhibit Integrator function at select loci. Altogether, these results show that the phosphatase module is critical and limiting at only a subset of Integrator regulated genes and point to recruitment of PP2A via IntS6 as a tunable step that can be used to modulate transcription termination efficiency.

## INTRODUCTION

Transcription of all eukaryotic protein-coding genes as well as most noncoding RNAs is catalyzed by RNA polymerase II (Pol II), which itself is regulated by many transcription factors and co-factors that act during the initiation, elongation, and termination steps (for review, see ^1,2,3^). These multiple macromolecular assemblies collectively help dictate the levels, processing, and functions of the transcripts produced. In recent years, the multi-subunit Integrator complex has emerged as a key transcriptional regulator across metazoans that controls the fates of many nascent RNAs (for review, see ^4–8^). Integrator is > 1.5 MDa and interacts with the C-terminal domain (CTD) of Pol II, with most of the more than 14 subunits in the Integrator complex lacking identifiable paralogs.^9^ The notable exceptions are Integrator subunits 11 (IntS11) and 9 (IntS9), which are homologous to the RNA endonuclease CPSF73 (cleavage and polyadenylation specificity factor 73, also known as CPSF3) and CPSF100 (also known as CPSF2) that cleave mRNA 3’ ends prior to poly(A) tail addition.^10^ The catalytic and scaffolding subunits of protein phosphatase 2A (PP2A) are also stably associated with Integrator, and thus the Integrator complex can have dual RNA cleavage and phosphatase catalytic activities.^11–17^

It has long been recognized that Integrator catalyzes RNA cleavage at the 3’ ends of nascent small nuclear RNA (snRNA) gene loci, enabling termination of transcription and release of the snRNA transcript that goes on to function in pre-mRNA splicing.^9,18^ Integrator has further been implicated in the processing of a variety of other Pol II transcripts, including enhancer RNAs,^19,20^ telomerase RNA,^21^ viral microRNAs,^22^ piwi-interacting RNAs,^23,24^ replication-dependent histone mRNAs,^25^ and many canonical protein-coding genes.^26–31^ It is thus perhaps not surprising that mis-regulation of Integrator is associated with developmental and disease phenotypes, including in humans.^4,32–35^

Analogous to how Integrator functions at snRNAs, the complex can endonucleolytically cleave nascent mRNAs to enable transcription termination at protein-coding genes coupled to nascent RNA release.^26–29^ These Integrator cleavage events predominantly do not occur at the 3’ ends of genes where CPSF73 acts but instead close to the transcription start site (TSS), often within 100 nucleotides at sites of Pol II pausing. This is at least in part because the Integrator phosphatase module can antagonize kinases that promote release of paused Pol II.^11–14^ Instead of allowing Pol II to productively elongate, Integrator causes transcription to prematurely terminate, and the released short mRNAs are rapidly degraded by the RNA exosome.^26–28^ In some cases, Integrator cleavage has been proposed to activate protein-coding transcription by enabling removal of stalled, inactive Pol II,^31^ but termination driven by Integrator can also attenuate gene expression by blocking full length mRNA production. Attenuation can be potent and readily observed in *Drosophila* cells, as exemplified by our prior genome-scale RNAi screen that revealed depletion of Integrator subunits resulted in the largest increase (among all genes tested) in the output of the Metallothionein A (MtnA) promoter.^27^ RNA-seq further revealed more than 400 mRNAs that were up-regulated in *Drosophila* cells upon depletion of IntS9, some by more than 100-fold, compared to only 49 mRNAs that were down-regulated. Integrator is thus a major attenuator of protein-coding gene outputs in *Drosophila*, yet how Integrator activity can be toggled on/off at a given gene locus depending on cellular transcriptional needs remains poorly understood.

There is emerging evidence that Integrator recruitment and/or activity is differentially regulated across gene loci (for review, see ^4^). For example, depletion of many non-catalytic Integrator subunits has only a minimal effect on snRNA 3’ end processing, yet these subunits are critical (especially IntS1, 2, 5, 6, 7, 8, and 12) for the ability of Integrator to attenuate the outputs of protein-coding genes.^27,36,37^ This suggests these non-catalytic Integrator subunits may act to ensure the proper balance between full-length mRNA production and premature termination at protein-coding genes, but the underlying molecular mechanisms remain unclear. There is also evidence that Integrator subunits can function independently of the rest of the complex, e.g. as part of the DNA damage response^38^ or to activate enhancers.^20^

Here, to identify regulatory subunits that enable the Integrator complex to be distinctly regulated in a locus-specific manner, we tested the effect of individually over-expressing each Integrator subunit in *Drosophila* cells. This revealed that IntS6 over-expression uniquely blocks Integrator function at a subset of protein-coding and long non-coding RNA genes. IntS6 can thus function in a dominant-negative manner, but IntS6 over-expression is not sufficient to fully disable Integrator across the genome: nascent snRNAs continue to be processed and the expression of many protein-coding genes remains attenuated by Integrator. The only loci affected by IntS6 over-expression are those where the PP2A phosphatase module is required for Integrator function, which is only a subset of gene loci bound by Integrator. We found that over-expressed IntS6 is a molecular sponge that titrates PP2A subunits from the rest of the Integrator complex by acting in a manner analogous to canonical PP2A regulatory B subunits. Altogether, these results show that the PP2A phosphatase module is critical at some, but not all genes for Integrator function and point to recruitment of PP2A via IntS6 as a tunable step that can control Integrator activity in a locus-specific manner.

## RESULTS

### Over-expression of IntS6 uniquely inhibits Integrator function at a subset of protein-coding promoters

Cleavage of nascent snRNA transcripts by Integrator is critical for production of functional snRNAs,^9^ while cleavage of nascent mRNAs by Integrator triggers premature transcription termination coupled to nascent transcript degradation and attenuation of protein-coding gene expression.^26,27^ Given these opposite effects on the ultimate gene outputs, we reasoned that the Integrator complex must be regulated in a distinct manner at protein-coding genes compared to snRNAs. To reveal the critical regulatory differences, we took advantage of a set of eGFP reporters that are driven by *Drosophila* protein-coding promoters **(Figure 1A, left)** and compared their regulation patterns to that of snRNA readthrough reporters that produce eGFP when Integrator fails to process the 3’ end of the encoded snRNA **(Figure 1A, right)**. Each of the examined protein-coding gene promoters, which included MtnA, Hemolectin (Hml), CG8620, anachronism (ana), and the small subunit of ribonucleoside reductase (RnrS), are normally attenuated by Integrator in *Drosophila* DL1 cells as eGFP mRNA expression increased upon treatment with double-stranded RNA (dsRNA) targeting IntS4, a scaffolding component of the Integrator endonuclease cleavage module,^39,40^ relative to treatment with a control (β-galactosidase, β-gal) dsRNA **(Figures 1B, left and S1A-C)**. As expected, the U4:39B and U5:34A snRNA readthrough reporters were likewise sensitive to depletion of IntS4 as demonstrated by a potent increase in eGFP levels due to Integrator not cleaving near the 3’ box sequence **(Figure 1B, left)**. In contrast, a reporter driven by the ubiquitin-63E (Ubi-p63e) promoter was not affected by IntS4 depletion **(Figure 1B, left)**, consistent with prior RNA-seq results that showed endogenous Ubi-p63e mRNA levels are not regulated by Integrator.^27^

**Figure 1.**
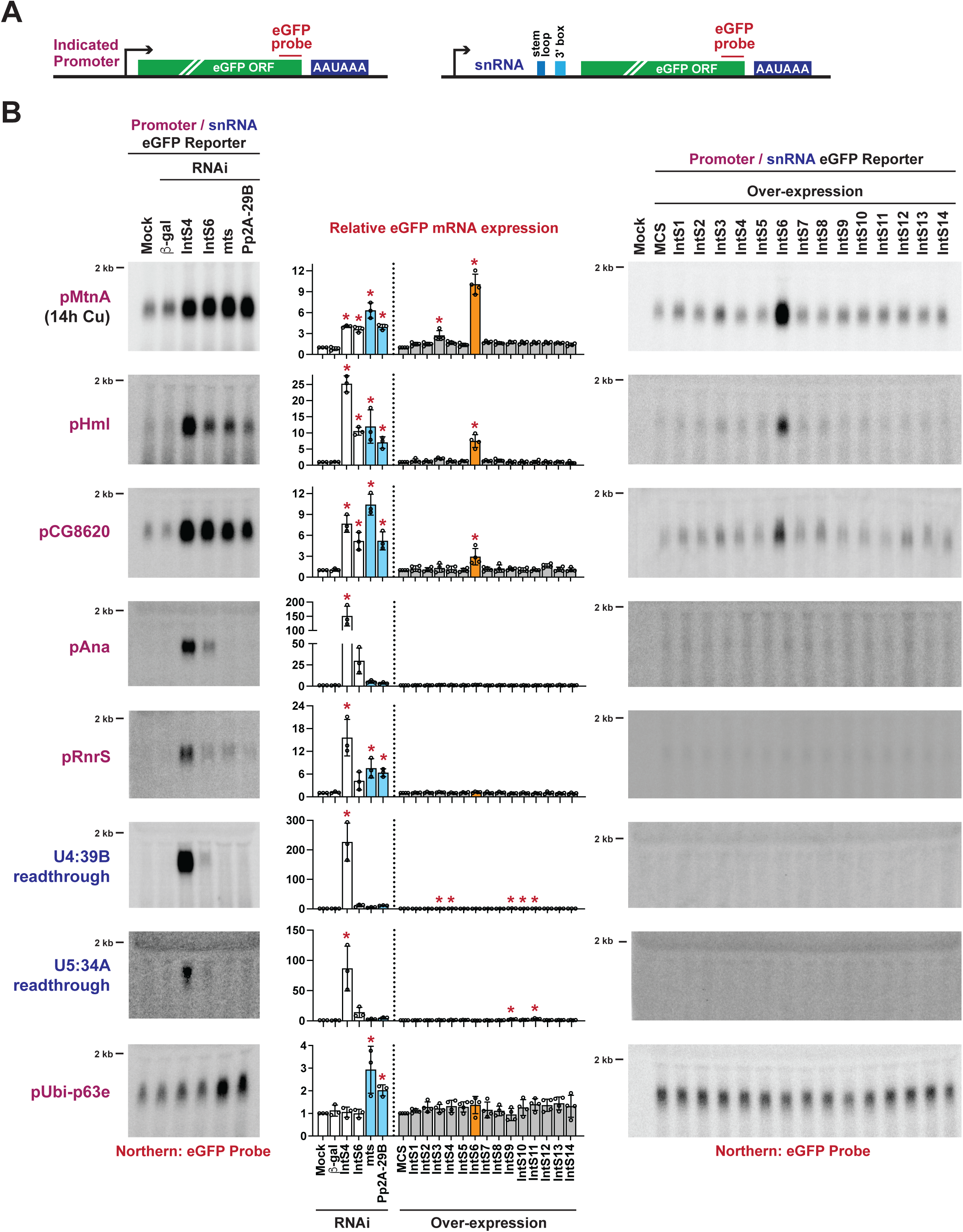
Integrator activity at a subset of reporter genes is lost upon over-expression of IntS6. **(A)** eGFP-based reporters to examine Integrator activity at protein-coding genes (left) and snRNAs (right). The promoter and 5’ UTR of each of the indicated protein-coding genes was cloned upstream of eGFP (left). The snRNA promoter, coding sequence, and downstream region were cloned upstream of eGFP, thereby enabling eGFP production when Integrator fails to process the snRNA 3’ end between the stem loop and 3’ box sequences (right). **(B)** Each individual reporter plasmid was transfected into DL1 cells that had been treated with the indicated dsRNAs (left) or was co-transfected with 100 ng of a plasmid that over-expresses a FLAG-tagged Integrator subunit from the Ubi-p63e promoter (right). A plasmid containing a multi-cloning site (MCS) driven by the Ubi-p63e promoter was used as a control for the over-expression experiments. CuSO_4_ was added for the last 14 h only when measuring eGFP production from the MtnA promoter. Total RNA was isolated and Northern blots (20 μg/lane) were used to measure expression of each eGFP reporter mRNA. Representative blots are shown and loading controls are provided in **Figure S1C**. RT-qPCR was used to quantify eGFP mRNA expression levels (middle). RNAi data were normalized to the mock samples and over-expression data were normalized to the MCS samples. Data are shown as mean ± SD, *N* ≥ 3. (*) *P* < 0.05.

We next took advantage of the eGFP reporters and examined the functional effects of individually over-expressing a FLAG-tagged version of each Integrator subunit (IntS1-IntS14) **(Figure S1D)**. Compared to over-expressing a control transgene that encodes a multicloning site (MCS), individual over-expression of almost all Integrator subunits had no or minimal effect on the amounts of eGFP mRNA produced from the reporters **(Figures 1B, right and S1C)**. The notable exception was IntS6 over-expression which caused significantly increased amounts of eGFP mRNA to be produced from the MtnA, Hml, and CG8620 promoters **(Figure 1B, right)**. There was, however, no change in the outputs of the Ana, RnrS, or snRNA readthrough reporters upon IntS6 over-expression **(Figure 1B, right)**. This suggested IntS6 over-expression may have a dominant-negative effect and be sufficient to disable Integrator function at a subset of protein-coding promoters, while simultaneously having no effect on the ability of the Integrator complex to attenuate other protein-coding promoters or process nascent snRNAs. To confirm the uniqueness of IntS6 over-expression for selectively inhibiting Integrator activity, increasing amounts of plasmids that over-express IntS6, IntS8 (which forms part of the Integrator shoulder module12), or IntS12 (which contains a PHD finger and can interact with the Integrator backbone module^41^) were transfected into DL1 cells. Effects on the eGFP reporters were then examined **(Figures S2A and S2B)**. A progressive increase in eGFP levels from the MtnA promoter was observed with titration of the IntS6 plasmid, while IntS8 and IntS12 over-expression had no or minimal effect even at the highest tested levels.

The 1,284 amino acid *Drosophila* IntS6 protein contains a conserved Von Willebrand factor type A (VWA) domain at its N-terminus, a > 250 amino acid region in the middle of the protein that is absent from IntS6 homologs in other species (e.g., human or mouse), and a conserved C-terminal domain that is observed in several additional proteins **(Figure S2C)**. To define the key regions in *Drosophila* IntS6 responsible for the dominant-negative effect, FLAG-tagged IntS6 over-expression plasmids with deletions from the N- or C-terminus were generated **(Figure S2C)** and transfected into DL1 cells **(Figure S2D)**. This revealed that the N-terminus (amino acids 1-600) and the C-terminus (amino acids 1197-1284) are each sufficient on their own to inhibit Integrator activity at the MtnA promoter **(Figure S2E)**. Both functional regions of IntS6 contain conserved protein domains, and we confirmed that over-expressing IntS6 homologs from human **(Figure S2F)** was likewise sufficient to inhibit Integrator activity at select promoters in *Drosophila* cells **(Figure S2G)**.

Depletion of IntS6 using RNAi interestingly revealed that this subunit was differentially required for Integrator function across the eGFP reporters **(Figures 1B, left and S1B)**. When compared to IntS4 depletion, IntS6 depletion had only a minimal effect on the output of the snRNA readthrough reporters (e.g., 226 vs. 11-fold increase in eGFP expression for the U4:39B reporter) while IntS4 and IntS6 depletion resulted in near equal increases in eGFP mRNA expression from the MtnA promoter (4.0 and 3.6-fold, respectively) **(Figure 1B, left)**. Hence, we concluded that IntS6 varies from dispensable (or near dispensable) to essential for Integrator function in *Drosophila* depending on which reporter is examined.

### A subset of endogenous protein-coding genes normally attenuated by Integrator are up-regulated upon IntS6 over-expression

To extend the results from the eGFP reporters to endogenous genes, we generated DL1 cell lines that stably maintain IntS6 or IntS12 transgenes driven by the copper inducible MtnA promoter (**Figure 2A**). Parental DL1 cells and the inducible cell lines were grown in the presence of Cu^2+^ for 24 h, which enabled robust induction of IntS6 and IntS12 proteins (18 and 17-fold, respectively) in the stable cell lines **(Figures S3A and S3B)**. As a control, we verified that expression of additional Integrator subunits (IntS8 and IntS11) was unchanged upon Cu^2+^ addition in all cell lines **(Figures S3A)**. Total RNA from 3 biological replicates was isolated from each cell line to generate ribosomal RNA depleted RNA-seq libraries (**Figures 2A** **and S3C)** which were then analyzed to identify differentially expressed transcripts **(Figure S3D**).

**Figure 2.**
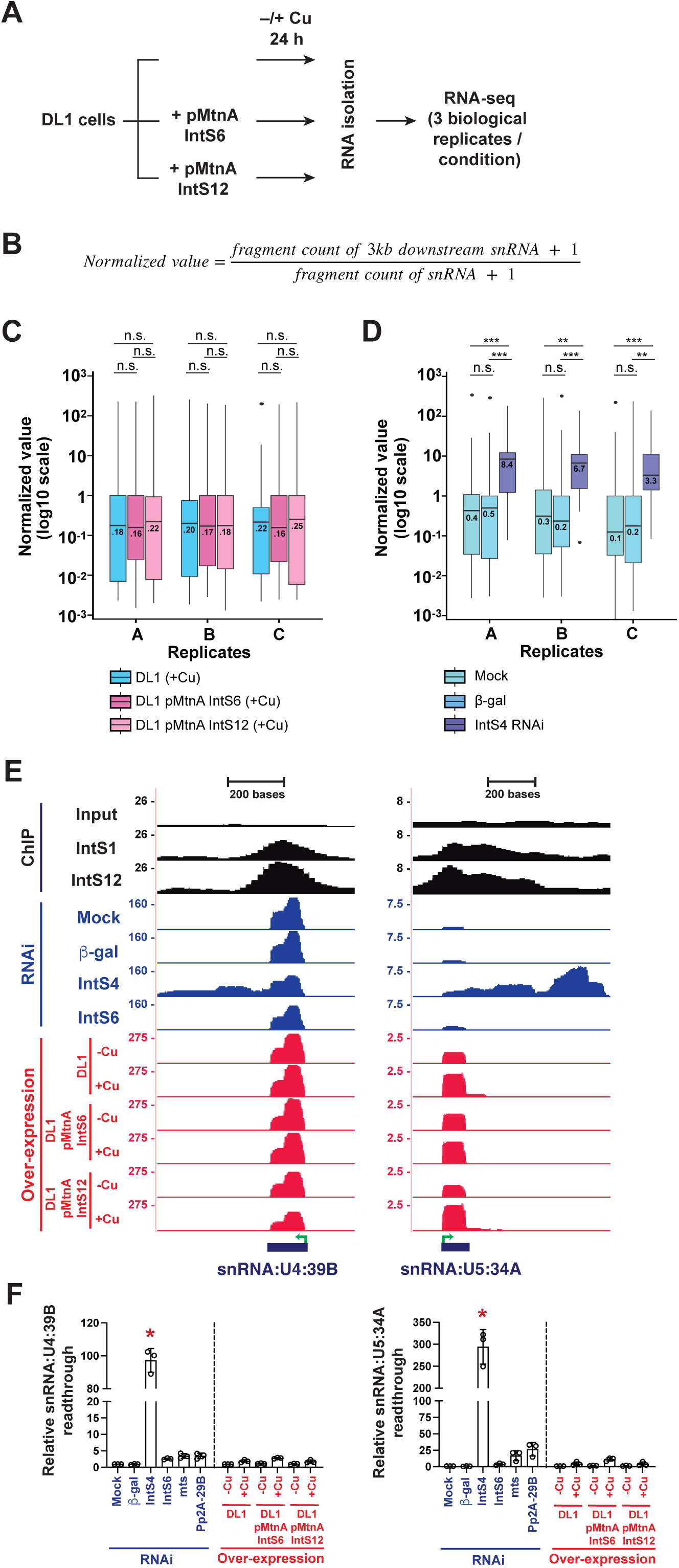
Over-expression of IntS6 does not affect Integrator activity at endogenous *Drosophila* snRNA loci. **(A)** Parental DL1 cells or DL1 cells stably maintaining IntS6 or IntS12 transgenes driven by the copper inducible MtnA promoter were grown for 3 d. 500 μM CuSO_4_ was added for the last 24 h prior to total RNA isolation from three independent biological replicates. rRNA depleted RNA-seq libraries were then generated, sequenced, and analyzed. **(B)** To quantify readthrough transcription downstream of endogenous snRNAs, the levels of RNA-seq fragments that map to the 3 kb downstream of mature snRNA 3’ ends were normalized to the levels of fragments that map to mature snRNA sequences. **(C-D)** Normalized values of endogenous snRNA readthrough among **(C)** CuSO_4_ treated parental DL1 cells and DL1 cells stably maintaining IntS6 or IntS12 transgenes, and **(D)** DL1 cells subjected to mock, control (β-gal) dsRNA, or IntS4 dsRNA treatments. Center lines represent medians, boxes represent interquartile ranges (IQRs), and whiskers represent extreme data points within 1.5× IQRs. Black points were outliners exceeding 1.5× IQRs. *P* values were calculated by *Wilcoxon* signed-rank test. (**) *P* < 0.01; (***) *P* < 0.001; *n.s.*, not significant. **(E)** UCSC genome browser tracks depicting exemplar snRNA loci. IntS1 and IntS12 ChIP-seq profiles in DL1 cells (GSE114467) are shown in black. RNA-seq data generated from DL1 cells treated for 3 d with control (β-gal), IntS4, or IntS6 dsRNAs are shown in blue. RNA-seq data generated from parental DL1 cells or DL1 cells stably maintaining copper-inducible IntS6 or IntS12 transgenes are shown in red. 500 μM CuSO_4_ was added for 24 h as indicated. Green arrow, transcription start site (TSS). **(F)** Readthrough downstream from snRNA transcripts was quantified using RT-qPCR. Data are shown as mean ± SD, *N* = 3. (*) *P* < 0.05.

To first examine effects on endogenous snRNA transcription termination, sequencing reads that mapped within 3 kb downstream of mature snRNA sequences were quantified and normalized to mature snRNA levels (**Figure 2B**). Over-expression of IntS6 or IntS12 had no significant effect on endogenous snRNA readthrough transcript levels (**Figure 2C** **and Table S1)**, mirroring the results obtained with the eGFP reporters **(Figure 1B, right)**. In contrast, depletion of IntS4 using RNAi resulted in the expected increase in endogenous snRNA readthrough transcripts (median increase >30-fold) due to disabled Integrator endonuclease activity (**Figure 2D** **and Table S1)**. These expression effects were readily observed at the endogenous U4:39B and U5:34A loci (**Figure 2E**) and confirmed using RT-qPCR (**Figure 2F**).

We next examined changes in protein-coding gene expression and found that over-expression of IntS6 (**Figures 3A and 3B**) but not IntS12 (**Figure 3C**) resulted in up-regulation of more than 100 genes with almost no genes being down-regulated (fold change > 1.5 and adjusted *P* < 0.001). Depending on whether the IntS6 over-expression data were compared to parental DL1 cells (**Figure 3A**) or to the IntS12 over-expression cell line (**Figure 3B**), slightly different numbers of up-regulated genes were identified but the vast majority (107 genes) overlapped (**Figure 3D** **and Table S2)**. These 107 genes are enriched in axon generation, protein folding, and heat response pathways **(Figure S3E)**, are significantly longer in length than the average *Drosophila* gene **(Figure S3F)**, and there is evidence that almost all are directly attenuated by Integrator in DL1 cells under normal conditions (**Figure 3D**). IntS1 and/or IntS12 chromatin immunoprecipitation sequencing (ChIP-seq) peaks were detected at 84 of the 107 genes (78%) (**Figures 3D** **and S4)**, and 102 of these genes (95%) were up-regulated (fold change > 1.5 and adjusted *P* < 0.001) upon RNAi depletion of IntS4 and/or IntS6 in the parental DL1 cells **(Figures 3D, S5A-C, and Table S3)**. There was, in fact, a strong positive correlation (∼0.8) between the degree of up-regulation observed for these genes upon IntS6 over-expression with that observed upon depletion of IntS4 **(Figure S5D)** or IntS6 **(Figure S5E)**. In addition, attenuation of many of the 107 genes is clearly dependent on the IntS11 endonuclease as their mRNA levels increased upon depletion of IntS11, and this could be rescued by expression of a wild-type IntS11 transgene but not by a catalytically dead (E203Q) IntS11 transgene **(Figures S5F and S5G)**.^9,26^ Using the form3 and CG6847 genes as examples, Integrator subunits are normally bound at their 5’ ends in DL1 cells **(Figure 3E, top)** and the encoded mRNAs were up-regulated when IntS4 or IntS6 was depleted by RNAi **(Figure 3E, middle)**. These mRNAs were likewise up-regulated when IntS6, but not IntS12, was over-expressed **(Figure 3E, bottom)**, and RT-qPCR confirmed the expression changes at both the mRNA (**Figure 3F**) and pre-mRNA levels **(Figure S6A)**. Similar effects on transcriptional outputs were confirmed at additional exemplar loci (CG8547, Kal1, wun2, and Wdp) as shown in **Figures S6A and S7A-D**.

**Figure 3.**
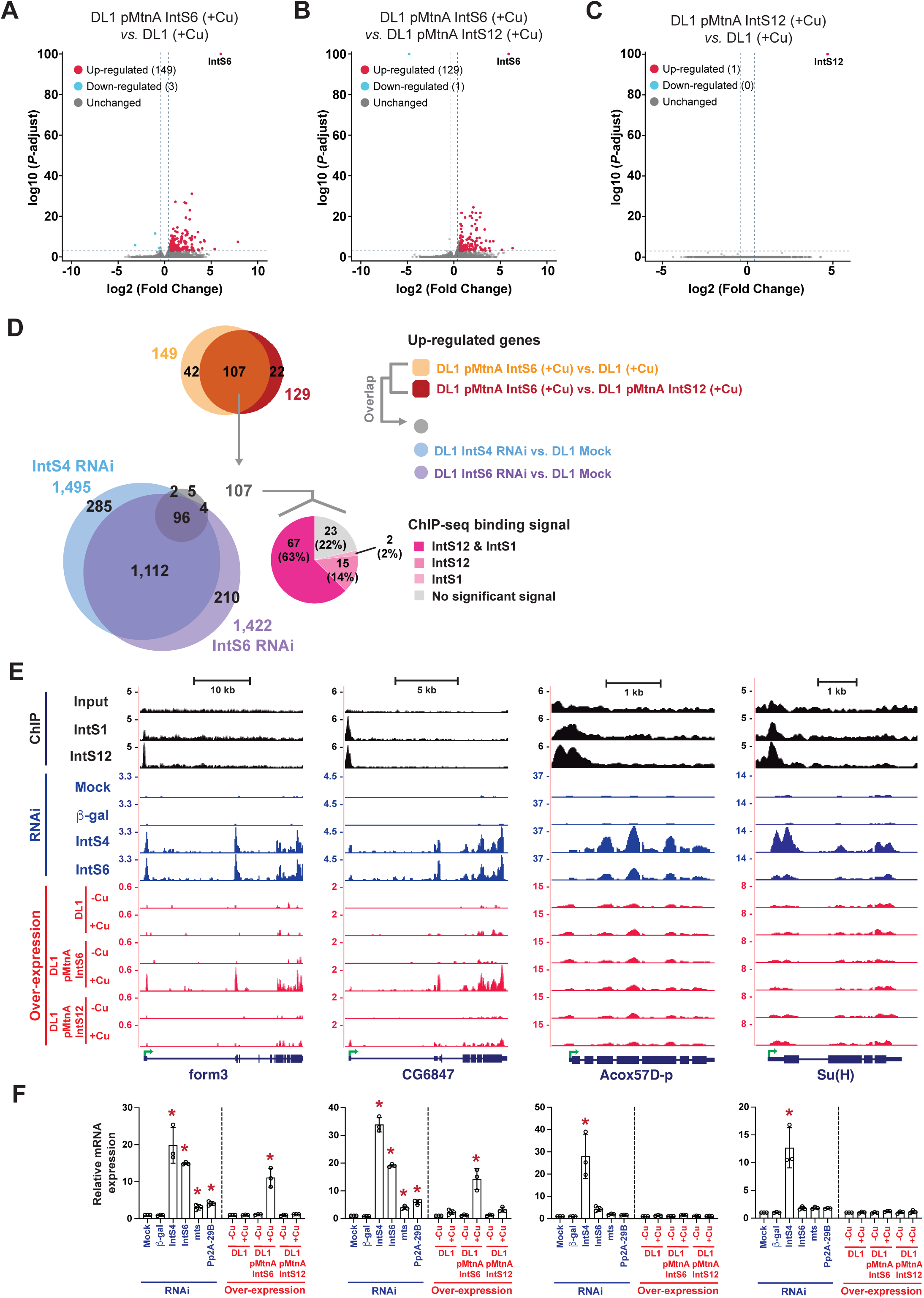
Over-expression of IntS6 blocks Integrator activity at a subset of endogenous *Drosophila* protein-coding genes. **(A-C)** The magnitude of change in mRNA expression compared with statistical significance (adjusted *P-*value) is shown as volcano plots. Endogenous mRNA expression levels upon IntS6 over-expression were compared to that in parental DL1 cells **(A)** or upon IntS12 over-expression **(B)**. mRNA expression levels upon IntS12 over-expression were compared to parental DL1 cells **(C)**. Threshold used to define differentially expressed mRNAs was |log2(fold change)| > 0.585 and adjusted *P* < 0.001. **(D)** The overlapping set of 107 protein-coding genes that were up-regulated upon IntS6 over-expression were compared to the sets of genes up-regulated upon RNAi depletion of IntS4 or IntS6 (left) and the sets of genes bound by IntS12 and/or IntS1 in DL1 cells in previously published (GSE114467) ChIP-seq experiments (right). **(E)** UCSC genome browser tracks depicting example protein-coding loci that are (form3, CG6847) or are not (Acox57D-p, Su(H)) affected by IntS6 over-expression. IntS1 and IntS12 ChIP-seq profiles in DL1 cells (GSE114467) are shown in black. RNA-seq data generated from DL1 cells treated for 3 d with control (β-gal), IntS4, or IntS6 dsRNAs are shown in blue. RNA-seq data generated from parental DL1 cells or DL1 cells stably maintaining copper-inducible IntS6 or IntS12 transgenes are shown in red. 500 μMCuSO4 was added for 24 h as indicated. Green arrow, transcription start site (TSS). **(F)** Expression of the indicated mRNAs (order as in **E**) was quantified using RT-qPCR. Data are shown as mean ± SD, *N* = 3. (*) *P* < 0.05.

Although IntS6 over-expression resulted in up-regulation of a set of protein-coding genes normally attenuated by Integrator, the vast majority of Integrator regulated genes were unaffected by IntS6 over-expression. More than 1,100 genes became up-regulated (fold change > 1.5 and adjusted *P* < 0.001) when IntS4 or IntS6 was depleted using RNAi **(Figures 3D, S5B, S5C, and Table S3)** and many of these genes are normally bound by IntS1 and/or IntS12, suggesting they are direct Integrator targets (discussed further in **Figure 5**). For example, Integrator subunits are bound at the 5’ ends of the Acox57D-p and Su(H) genes **(Figure 3E, top)** and these mRNAs were up-regulated upon depletion of IntS4 **(Figure 3E, middle)**, yet they were unaffected by IntS6 over-expression **(Figures 3E, bottom, 3F, and S6B)**. Similar expression patterns were observed at the endogenous Ana and RnrS loci **(Figures S7E and S7F)**, mirroring the lack of effect that was observed when these promoters were tested in the eGFP reporter assay (**Figure 1B**). We thus conclude that IntS6 over-expression has a dominant-negative effect on Integrator function at only a subset of endogenous target loci.

### IntS6 over-expression blocks Integrator function by titrating PP2A

We next aimed to understand why only a subset of Integrator regulated loci are affected by IntS6 over-expression. Recent cryo-electron microscopy (cryoEM) efforts characterizing the Integrator complex revealed that IntS6 is located far (> 75 angstroms) from the IntS11 endonuclease active site and instead contacts the protein phosphatase 2A (PP2A) subunits (**Figure 4A**).^12,14^ This led us to hypothesize that IntS6 over-expression may somehow affect the phosphatase module of the Integrator complex (in line with recent work^13^). PP2A typically functions as a trimer in which a regulatory B subunit (twins (tws), widerborst (wdb), Connector of kinase to AP-1 (Cka), or well-rounded (wrd) in *Drosophila*) enables the scaffolding (A) and catalytic (C) subunits (Pp2A-29B and microtubule star (mts), respectively, in *Drosophila*) to be targeted to a variety of substrates in cells.^42–44^ Both the A (Pp2A-29B) and C (mts) subunits are present in the Integrator complex, but no canonical B subunits have been detected in association with Integrator (**Figure 4A**).^11–14^ Consistent with IntS6 over-expression altering the expression of genes normally controlled by PP2A subunits, significant overlap was observed when comparing the list of 107 up-regulated genes to those that were up-regulated when the A (Pp2A-29B) or C (mts) subunit was depleted using RNAi in DL1 cells **(Figures 4B, S5A, and Table S3)**. This suggested that IntS6 over-expression may disable the phosphatase activity of the Integrator complex, and we hypothesized this could be due to IntS6 sponging or titrating the PP2A subunits from the rest of the complex. Consistent with such a model, the PP2A C (mts) subunit was co-immunoprecipitated with over-expressed FLAG-tagged IntS6, but not with over-expressed IntS12 (**Figure 4C**). This result nicely mirrors our observation that only IntS6 over-expression (not IntS12) alters Integrator complex function (**Figures 1, 2, and 3)**.

**Figure 4.**
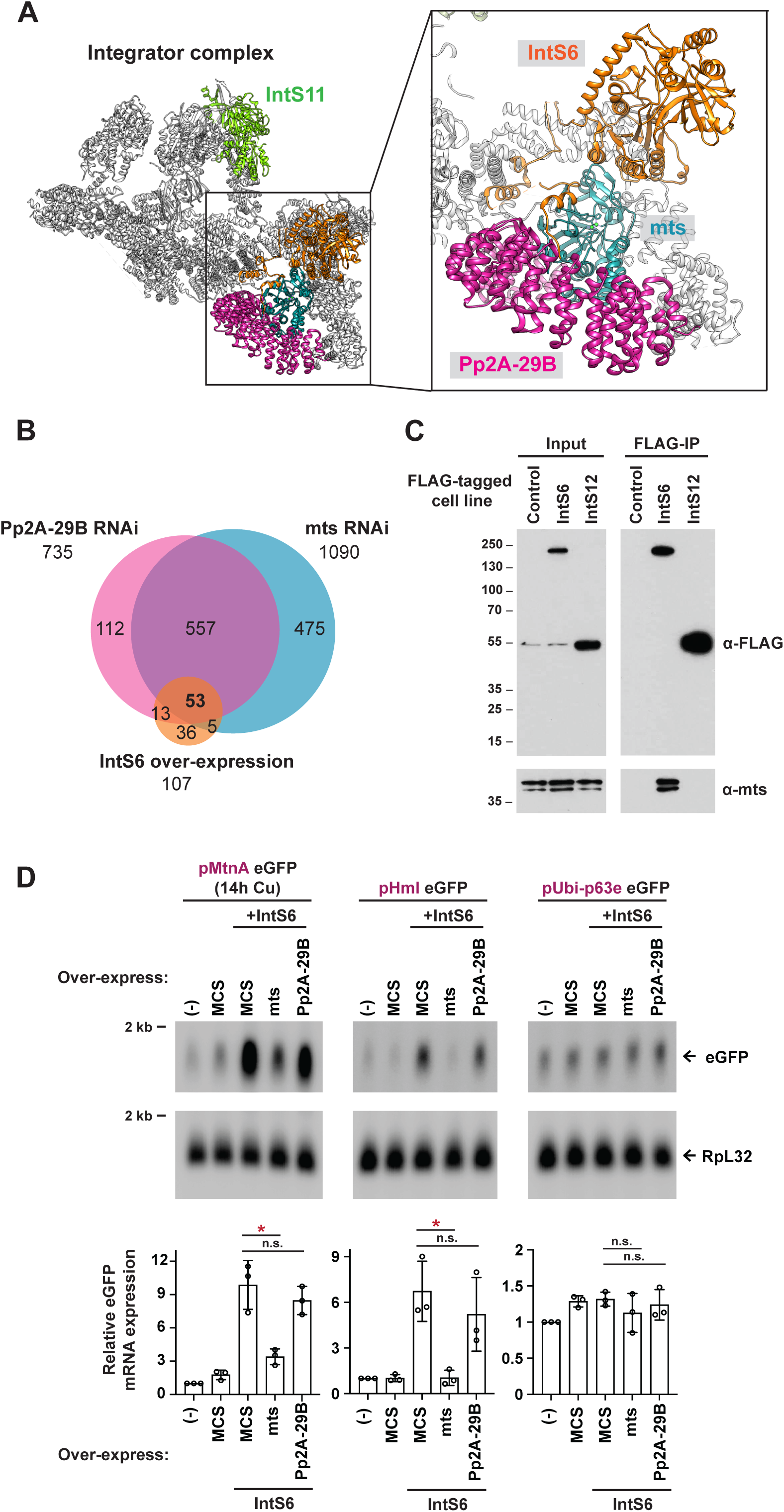
IntS6 over-expression titrates the catalytic subunit of PP2A and causes it to be limiting for Integrator activity. **(A)** Cryo-EM structure (PDB: 7PKS) of the human Integrator complex,^14^ highlighting the positions of the RNA endonuclease IntS11 (green), IntS6 (orange), and PP2A subunits (teal and pink). There are direct contacts between IntS6 and the PP2A subunits. **(B)** DL1 cells were treated for 3 d with control (β-gal), Pp2A-29B, or mts dsRNAs and RNA-seq data generated. The sets of endogenous genes up-regulated upon Pp2A-29B or mts depletion (fold change > 1.5 and adjusted *P* value < 0.001) were compared to the set of 107 genes that were up-regulated upon over-expression of IntS6. **(C)** Parental DL1 cells (Control) and DL1 cells stably maintaining inducible FLAG-tagged IntS6 or IntS12 transgenes were treated with 500 μM CuSO_4_ for 24 h to induce transgene expression. Immunoprecipitation (IP) using anti-FLAG resin was then performed. Western blots of input nuclear extracts (left) and IP (right) are shown. **(D)** DL1 cells were co-transfected with 300 ng of eGFP reporter plasmid and 100 ng of the indicated IntS6/PP2A subunit over-expression plasmids (driven by the Ubi-p63e promoter). Empty vector (pUb 3xFLAG MCS) was added as needed so that 500 ng DNA was transfected in all samples. CuSO_4_ was added for the last 14 h only when measuring eGFP production from the MtnA promoter. Northern blots (20 μg/lane) were used to quantify expression of each eGFP reporter mRNA. Representative blots are shown and RpL32 mRNA was used as a loading control. Data are shown as mean ± SD, *N* = 3. (*) *P* < 0.05; *n.s.*, not significant.

Prior work in *S. cerevisiae* has shown that the PP2A A subunit is expressed in excess over the C subunit,^45^ suggesting a possible role for stoichiometry in control of PP2A function. If PP2A subunits are indeed limiting in DL1 cells, we reasoned that over-expressing the C (mts) and/or A (Pp2A-29B) subunits should be sufficient to reverse the gene expression changes caused by IntS6 over-expression. Increasing the expression of the C (mts) subunit, but not the A (Pp2A-29B) subunit, was sufficient to reduce the amount of eGFP mRNA produced from the MtnA and Hml promoter reporters to near wild-type levels, indicating a restoration of Integrator activity at these promoters (**Figure 4D**). These data further indicate that the PP2A C (mts) subunit can be limiting for Integrator function. As expected, there was no change in the output of the control Ubi-p63e promoter, which is not regulated by Integrator, when PP2A subunits were over-expressed (**Figure 4D**).

### The phosphatase module of Integrator is critical at only a subset of Integrator-regulated loci

Based on the above results, a model emerged in which IntS6 over-expression inhibits Integrator function by binding/titrating PP2A subunits and thereby removing the phosphatase activity from the Integrator complex. Given that IntS6 over-expression only affects the outputs of a subset of Integrator regulated eGFP reporters (**Figure 1**) and endogenous genes (**Figures 2 and 3**), this model suggests that the PP2A phosphatase activity is differentially required across loci for regulation by the Integrator complex. To test this hypothesis, RNAi was used to deplete the PP2A A (Pp2A-29B) or C (mts) subunits **(Figure S1A)** and we then examined the effect on the outputs of the eGFP reporters **(Figure 1B, left)**. eGFP expression from the MtnA, Hml, and CG8620 promoters all increased upon depletion of PP2A subunits, consistent with a critical role for phosphatase activity in enabling Integrator function at these promoters. In stark contrast, there was no or minimal change in the outputs of the Ana, RnrS, and snRNA readthrough reporters upon depletion of PP2A subunits **(Figure 1B, left)**. These results perfectly mirror the effects observed with IntS6 over-expression: only at promoters where PP2A subunits are required (MtnA, Hml, and CG8620) does IntS6 over-expression result in increased eGFP reporter expression **(Figure 1B, right)**.

We next examined individual endogenous snRNA (**Figure 2F**) and protein-coding genes (**Figures 3F** **and S7)** using RT-qPCR and again observed that only genes that require PP2A subunits for Integrator function were significantly affected by IntS6 over-expression. For example, the Integrator phosphatase module is not required at the endogenous Acox57D-p and Su(H) genes as there was only a minimal increase in expression of these mRNAs upon depletion of the PP2A A (Pp2A-29B) or C (mts) subunits, and these loci were unaffected when IntS6 was over-expressed (**Figure 3F**). In contrast, PP2A subunits were required for Integrator function at the form3 and CG6847 loci, and the expression of these genes was increased with IntS6 over-expression (**Figure 3F**).

To determine how broadly the phosphatase activity is required for Integrator function across the genome, we first examined readthrough transcription downstream of all endogenous snRNA transcripts. Depletion of PP2A C (mts) or A (Pp2A-29B) resulted in modest effects (compared to IntS4 depletion) that were not statistically significant (**Figures 5A and 5B**). We then used ChIP-seq data from DL1 cells^26^ to identify 3,932 protein-coding genes with peaks of IntS1 and/or IntS12 binding located ±1 kb of gene bodies **(Figures 5C, middle and S4)**. Most (>70%) of these peaks were close to transcription start sites (TSSs) **(Figures S4C and S4D)** and we reasoned these 3,932 genes represent loci where Integrator may have direct effects. We thus determined if their expression changed upon depletion of IntS4, IntS6, PP2A C (mts), or PP2A A (Pp2A-29B) **(Figure 5C, top, S5A, and Table S3)**. Consistent with Integrator catalyzing premature transcription termination events, most of the protein-coding genes that changed were up-regulated (fold change > 1.5 and adjusted *P* < 0.001) upon depletion of IntS4 or IntS6 **(Figure 5C, top)**. Of the 3,932 genes bound by Integrator, 903 and 915 were up-regulated upon IntS4 or IntS6 depletion, respectively, and ∼40% of these genes were also up-regulated upon depletion of PP2A C (mts) and/or A (Pp2A-29B) **(Figure 5C, bottom)**. This suggests that the phosphatase activity of Integrator may be required at ∼40% of protein-coding genes attenuated by Integrator, while it is dispensable at many other genes that are nonetheless still controlled by the complex. This result helps explain why IntS6 over-expression had no effect on most Integrator regulated genes (**Figure 3D**). The 107 protein-coding genes that were up-regulated by IntS6 over-expression may represent those genes that are most sensitive to tuning of Integrator function by PP2A activity.

**Figure 5.**
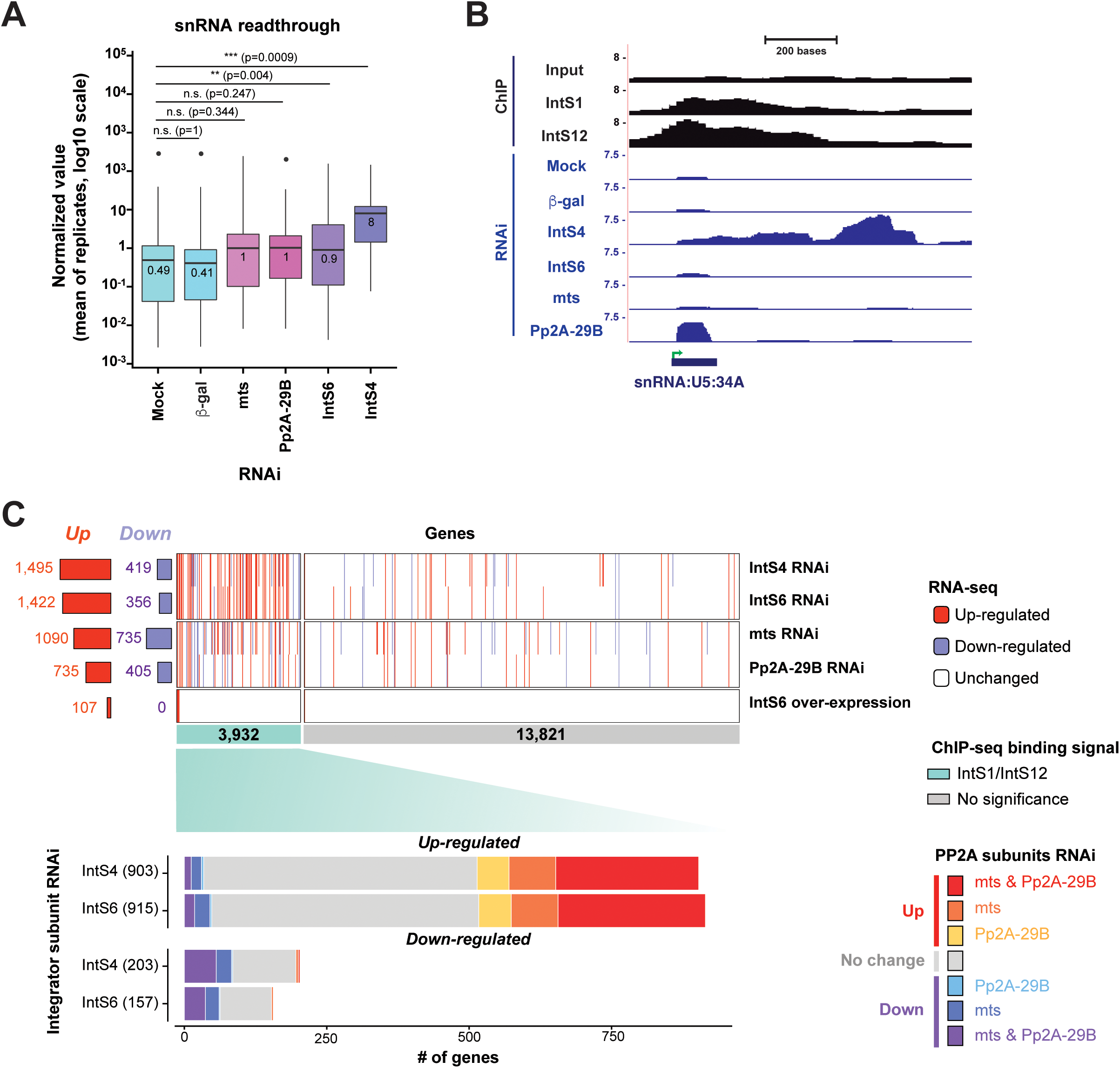
The phosphatase module is differentially required for Integrator activity across the genome. **(A)** Means of normalized values of endogenous snRNA readthrough. Endogenous snRNA readthrough values (see Figure 2B) were calculated from RNA-seq data of DL1 cells subjected to a mock treatment or treatment with β gal, mts, Pp2A-29B, IntS6, or IntS4 dsRNAs (3 independent biological replicates). Center lines represent medians, boxes represent interquartile ranges (IQRs), and whiskers represent extreme data points within 1.5× IQRs. Black points were outliners exceeding 1.5× IQRs. *P* values were calculated by *Wilcoxon* signed-rank test. (**) *P* < 0.01; (***) *P* < 0.001; *n.s.*, not significant. **(B)** ChIP-seq and RNA-seq tracks at the U5:34A snRNA locus. IntS1 and IntS12 ChIP-seq profiles in DL1 cells (GSE114467) are shown in black. RNA-seq data generated from DL1 cells treated for 3 d with control (β-gal), IntS4, IntS6, mts, or Pp2A-29B dsRNAs are shown in blue. Green arrow, transcription start site (TSS). **(C)** RNA-seq was used to define genes that were up- or down-regulated (|log2(fold change)| > 0.585 and adjusted *P* < 0.001) upon IntS4, IntS6, mts, or Pp2A-29B depletion using RNAi or upon IntS6 over-expression (top). These gene lists were then stratified by ChIP-seq data that identified 3,932 protein-coding genes with peaks of IntS1 and/or IntS12 binding located ±1 kb of gene bodies in DL1 cells (green, middle). For genes bound by Integrator subunits and differentially regulated upon IntS4 or IntS6 depletion, the effect of depleting PP2A subunits on their expression is graphed (bottom).

### IntS6 behaves like a canonical PP2A B subunit

The PP2A holoenzyme typically functions as a heterotrimer consisting of an A, B, and C subunit (**Figure 6A**), but no canonical B subunits have been detected in the Integrator complex.^11–14^ This has made it somewhat unclear how the Integrator phosphatase activity is controlled and regulated. Because IntS6 binds the PP2A A and C subunit in Integrator (**Figure 4A**) and IntS6 over-expression is sufficient to titrate PP2A activity (**Figure 4D**), we reasoned that IntS6 may be a previously unappreciated regulatory B subunit. If true, this model predicts that over-expression of canonical B subunits (tws, wdb, Cka, or wrd in *Drosophila*) may likewise be sufficient to titrate the PP2A C subunit and inhibit the attenuation activity of Integrator. We thus individually over-expressed FLAG-tagged versions of each PP2A B subunit (**Figure 6B**) and examined the functional effects on the eGFP reporters (**Figure 6C**). Over-expression of the canonical B subunits tws and wdb did indeed lead to increased eGFP expression from the MtnA and Hml promoters, but not from the control Ubi-p63e promoter that is not regulated by Integrator (**Figure 6C**). It should be noted the effects were not as large as that observed with IntS6 over-expression, despite the FLAG-tagged versions of tws and wdb being expressed at higher levels than FLAG-tagged IntS6 (**Figure 6B**). In total, these results indicate that IntS6 behaves, at least in some ways, similar to canonical B subunits and that alterations in the levels of B subunits not normally associated with Integrator can nonetheless still affect the efficiency of select premature transcription termination events.

**Figure 6.**
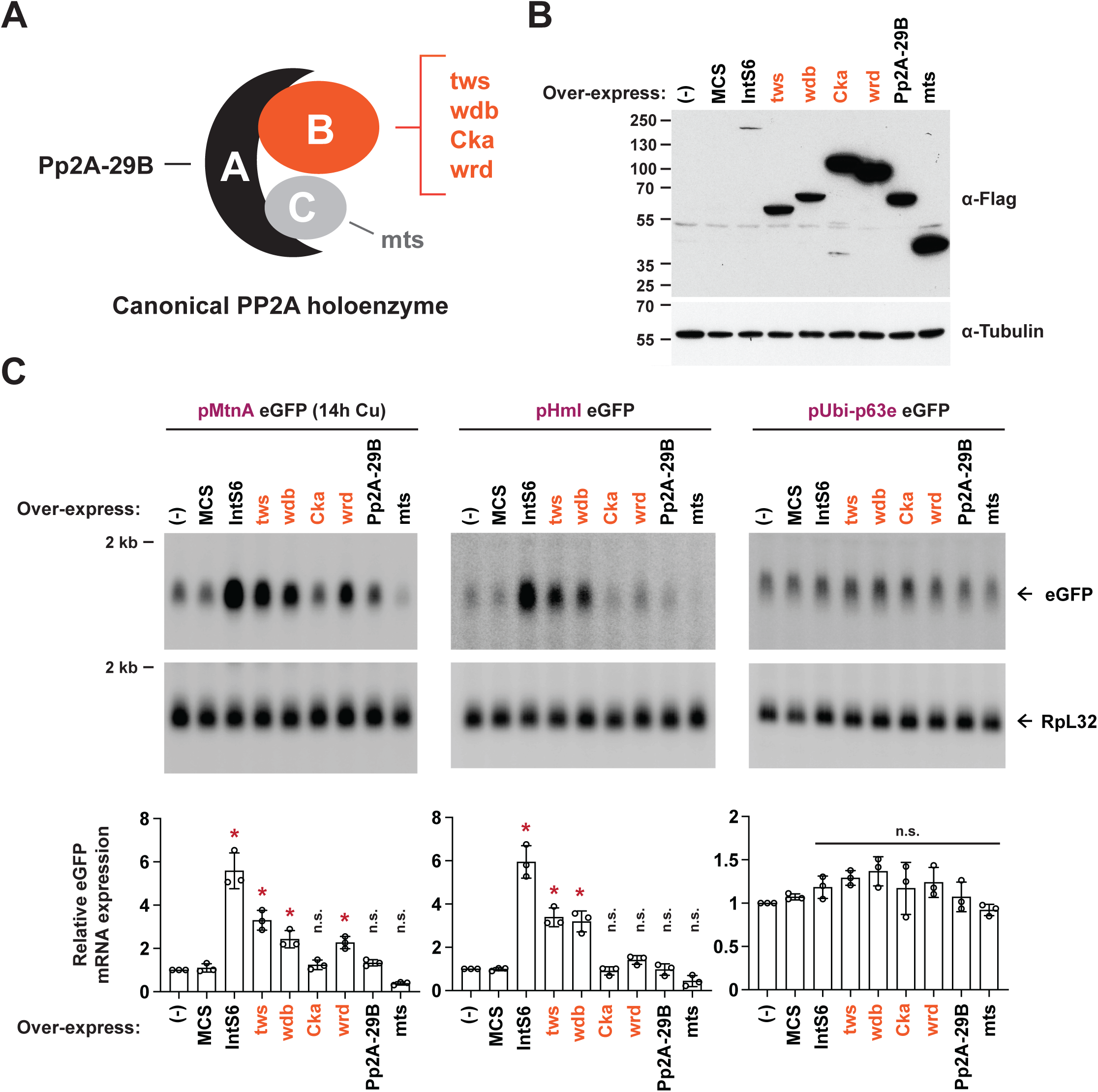
Over-expression of canonical PP2A B subunits can inhibit Integrator activity. **(A)** Schematic of the canonical *Drosophila* PP2A holoenzyme that consists of a scaffolding A subunit (Pp2A-29B), a catalytic subunit (mts), and a variable regulatory B subunit (tws, wdb, Cka, or wrd). **(B)** DL1 cells were transfected with 500 ng of the indicated FLAG-tagged expression plasmid and total protein was harvested after 48 h. A plasmid containing a multi-cloning site (MCS) was used as a control. Western blot analysis using an antibody that recognizes FLAG was used to confirm expression of each subunit. α-tubulin was used as a loading control. **(C)** DL1 cells were co-transfected with 400 ng of eGFP reporter plasmid and 100 ng of the indicated PP2A subunit over-expression plasmid (driven by the Ubi-p63e promoter). CuSO_4_ was added for the last 14 h only when measuring eGFP production from the MtnA promoter. Northern blots (20 μg/lane) were used to quantify expression of each eGFP reporter mRNA. Representative blots are shown and RpL32 mRNA was used as a loading control. Data are shown as mean ± SD, *N* = 3. (*) P < 0.05; n.s., not significant.

## DISCUSSION

The Integrator complex has RNA endonuclease and protein phosphatase activities, but it has remained unclear if the coordinated action of both activities is required for complex function or if they can be independently harnessed in a locus-specific manner. Here, using reporter genes as well as transcriptomics, we revealed that (i) the phosphatase module is functionally required only at a subset of *Drosophila* genes that are regulated by Integrator and (ii) that the ability of the PP2A catalytic subunit to act as part of Integrator can be tuned by the levels of IntS6 as well as canonical PP2A regulatory B subunits. Integrator regulates transcription elongation (**Figure 7A**) and, on one hand, we identified a set of protein-coding genes that became potently de-attenuated when IntS4 (a component of the endonuclease module), IntS6, or PP2A subunit levels were modulated, indicating that the endonuclease and phosphatase modules are both critical for Integrator function at these genes **(Figure 7B, left)**. Meanwhile, cleavage of nascent snRNAs as well as attenuation of many other protein-coding genes required the Integrator endonuclease module but was largely unaffected by modulation of IntS6 or PP2A subunit levels. This suggests the phosphatase module plays little or perhaps even no functional role at these loci **(Figure 7B, right)**. Prior work suggested the phosphatase module is broadly necessary for Integrator function across *Drosophila* protein-coding genes and snRNAs,^11^ which contrasts with the locus specificity we observed. However, most of the prior contrasting conclusions were based on manipulating IntS8 (by mutating a 4 amino acid region that binds the Pp2A-29B subunit) rather than directly testing roles for PP2A subunits themselves, especially on a genome-wide scale. Indeed, when the catalytic subunit of PP2A, IntS6, or IntS8 were individually depleted in human cell lines, no or minimal effect on transcription termination of exemplar snRNA genes was observed,^12,13^ which mirrors the results described here. It thus appears that the previously described IntS8 mutant may cause additional unintended effects beyond modulating recruitment of the Integrator phosphatase module.

**Figure 7.**
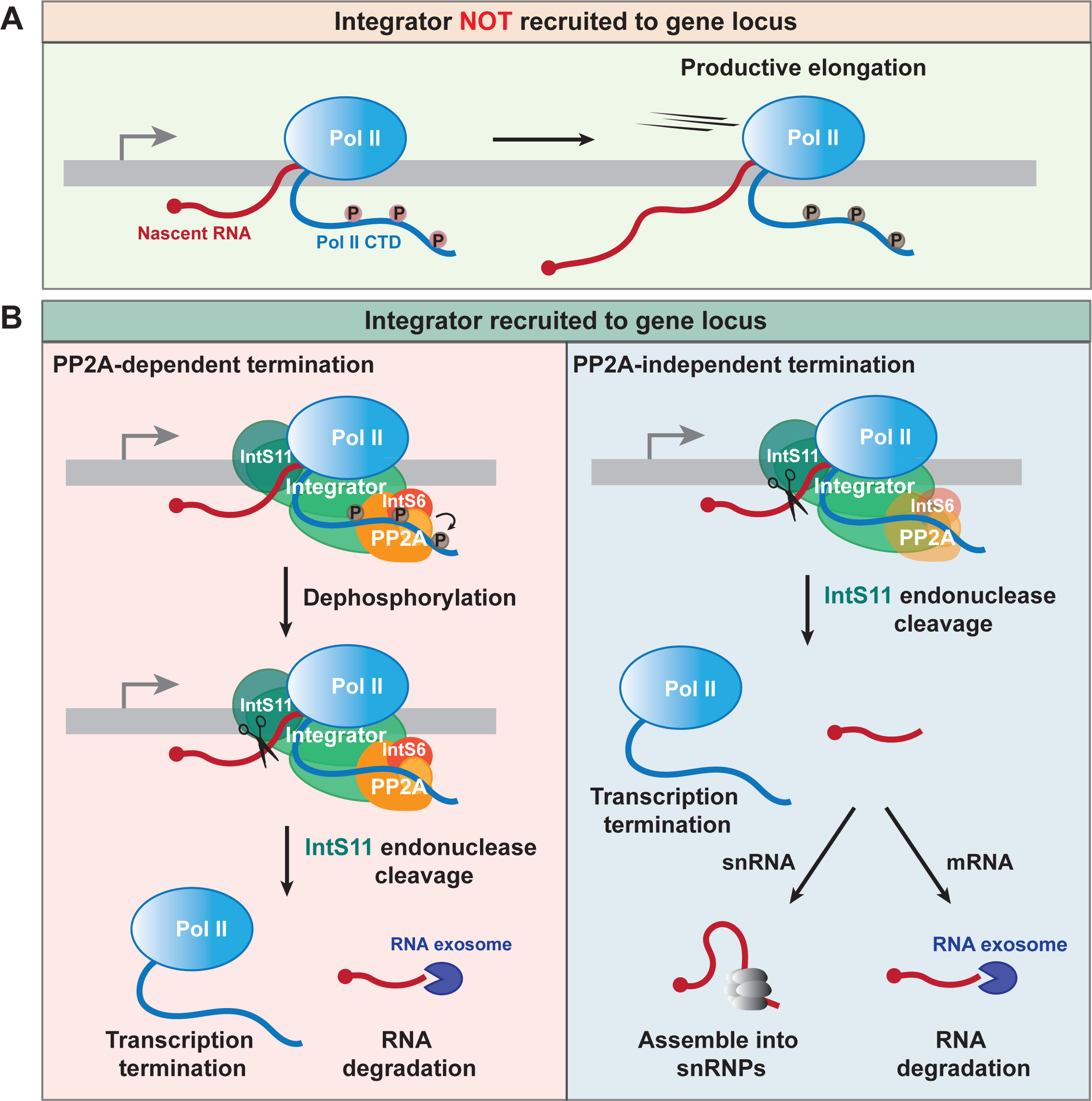
Integrator can catalyze transcription termination via PP2A-dependent and -independent mechanisms. **(A)** In the absence of Integrator, Pol II is able to productively elongate. **(B)** Integrator recruitment facilitates transcription termination. (Left) At some protein-coding genes, the Integrator phosphatase module must act prior to or, at minimum, simultaneously with the IntS11 endonuclease to enable cleavage of the nascent RNA, which is subsequently degraded by the RNA exosome. (Right) In contrast, the phosphatase module is dispensable for Integrator function at snRNA and many other protein-coding gene loci. Cleavage by IntS11 enables stable snRNAs to be produced and prematurely terminated mRNAs to be rapidly degraded.

Structural studies using cryoEM have made it increasingly clear that Integrator contains discrete endonuclease, phosphatase, and auxiliary modules that each bind around a core set of Integrator subunits.^12,14,39,46,47^ Overall complex integrity is disrupted upon depletion of a core subunit (e.g., IntS2), while depletion of module specific subunits, including IntS6 or its binding partner IntS8, does not appreciably affect the binding of other modules.^11,13^ Our functional data support this concept of Integrator complex modularity as we found that altering IntS6 or PP2A subunit levels had no effect on endonuclease activity at many gene loci (e.g., snRNAs continued to be efficiently processed at their 3’ ends). Why then is the Integrator phosphatase module required at a subset of loci? It has been recently shown that PP2A can dephosphorylate components of the Pol II complex, e.g. Spt5 or the C-terminal domain of Pol II itself, to antagonize kinases that stimulate Pol II pause release and productive elongation.^11–13^ Based on our data, it appears that the order by which the Integrator RNA endonuclease and protein phosphatase activities act can be critical. In particular, at a subset of genes, the Integrator phosphatase module must act prior to or, at minimum, simultaneously with the IntS11 endonuclease to slow/pause Pol II and thereby enable cleavage of the nascent RNA **(Figure 7B, left)**. In the absence of phosphatase activity, the temporal window of opportunity for IntS11 to cleave may be too short and/or IntS11 may fail to achieve an active conformation that accommodates RNA.^12,14^ The Pol II juggernaut^2^ is hence unable to be stopped by Integrator and instead elongates to produce full length, functional mRNAs. Given that RNA cleavage by IntS11 is presumably a non-reversible step, modulating PP2A activity and thus the IntS11 temporal window of opportunity represents an important way that the efficiency of premature termination events can be tuned. Here, we found that changes in protein levels can modulate the interaction of Integrator with the phosphatase module, and we propose that post-translational modifications (e.g., on IntS6 or the PP2A subunits) or the action of additional binding partners likely also tunes Integrator phosphatase activity.

At genes regulated by Integrator in a manner independent of the phosphatase module **(Figure 7B, right)**, there are presumably alternative mechanisms that enable a sufficiently long temporal window of opportunity for IntS11 to cleave. A conserved but relatively degenerate 3’ box sequence is located 9-19 nucleotides downstream from the 3’ ends of mature snRNA transcripts that is required for Integrator cleavage.^48^ Details of the underlying molecular mechanism unfortunately remain poorly understood and sequences resembling a 3’ box are not present at most protein-coding genes attenuated by Integrator. Instead, nucleosome occupancy^31^ or redundant phosphatases may act to stall or pause Pol II near protein-coding promoters, e.g. protein phosphatase 1 (PP1) which is known to help slow Pol II near polyadenylation signals.^49–51^ Differences in Pol II speed/dynamics caused by histone modifications or RNA sequence/structure may also be at play.

PP2A is responsible for the vast majority of serine/threonine phosphatase activity in eukaryotes and participates in many signaling cascades (for review, see ^42,43,44^). We nonetheless found that expression of the PP2A catalytic subunit (mts) can be limiting for Integrator function. Increasing the levels of IntS6 or canonical regulatory B subunits was sufficient to titrate PP2A subunits from Integrator, pointing to a critical role for stoichiometry in controlling PP2A function and Integrator activity. Consistent with this idea, PP2A regulatory B subunits are differentially expressed across tissues and developmental time, and IntS6 is known to be lost or down-regulated in several types of human cancer.^52,53^ We thus propose that modulating the expression, localization, or modification status of B subunits (including IntS6) may be used by cells to couple rapid dephosphorylation of protein substrates to longer term transcriptional changes via modulation of Integrator activity. It is also noteworthy that IntS6 underwent a gene duplication in many mammals, including humans (but not *Drosophila*), and it will be interesting in the future to determine whether IntS6 and IntS6-like (IntS6L) modulate PP2A activity in distinct ways and/or at distinct sets of Integrator regulated genes.

In summary, the Integrator complex controls the fates of many nascent RNAs in metazoans and our work has revealed key insights how its functional modules, especially the IntS6/PP2A phosphatase module, can be differentially employed across gene loci. Further work will be required to clarify if all the Integrator modules are normally recruited as a group *in vivo* or if they can be selectively assembled at a gene locus. Regardless, our results make clear that the Integrator RNA endonuclease and protein phosphatase activities do not always have to be coordinated with one another, but instead can act independently. Modulation of phosphatase activity by IntS6 is sufficient to disable Integrator at some loci, and we anticipate there will be additional mechanisms that tune Integrator in a gene-specific manner to meet a cell’s ever-changing transcriptional needs.

## ACKNOWLEDGMENTS

We thank Veerle Janssens and members of the Wilusz and Yang labs for helpful discussions. This paper is dedicated to the memory of our dear colleague Deirdre Tatomer, who passed away while this work was ongoing. Supported by National Institutes of Health grant R35-GM119735 (to J.E.W.), Cancer Prevention & Research Institute of Texas grant RR210031 (to J.E.W.), and National Natural Science Foundation of China grant 31925011 (to L.Y.). J.E.W. is a CPRIT Scholar Cancer Research.

## AUTHOR CONTRIBUTIONS

D.C.T. and J.E.W. conceived the project. R.F., D.L., M.T., A.P.S., C.J.F., M.S.M.-F., M.C.M., and D.C.T. performed experiments and analyzed data. S.-N.Z., X.-K.M., and L.Y. analyzed the RNA-seq and ChIP-seq results. R.F., S.-N. Z., L.Y., and J.E.W. wrote the manuscript with input from the other authors.

## EXPERIMENTAL PROCEDURES

### Cell culture

*Drosophila* DL1 cells were cultured at 25°C in Schneider’s *Drosophila* medium (Thermo Fisher Scientific 21720024), supplemented with 10% (v/v) fetal bovine serum (HyClone SH30396.03), 1% (v/v) penicillin-streptomycin (Thermo Fisher Scientific 15140122), and 1% (v/v) L-glutamine (Thermo Fisher Scientific 35050061).

### Expression plasmids

*Drosophila* reporter plasmids expressing eGFP under the control of the inducible Metallothionein A promoter (Hy_pMT eGFP SV40; Addgene #69911), Hml promoter (Hy_pHml eGFP SV40; Addgene #132645), CG8620 promoter (Hy_pCG8620 eGFP SV40; Addgene #132646), or Ubi-p63e promoter (Hy_pUbi-p63e eGFP SV40; Addgene #132650) were described previously.^27,54^ Reporter plasmids expressing eGFP under the control of the RnrS promoter (Hy_pRnrS eGFP SV40, Addgene #195062) or Ana promoter (Hy_pAna eGFP SV40, Addgene #195063) were generated from Hy_pPepck1 eGFP SV40 (Addgene #132644) as detailed in the **Supplemental Material**. Plasmids expressing individual Integrator or PP2A subunits under the control of the Ubi-p63e promoter were generated using the previously described pUb 3xFLAG MCS plasmid^55^ as detailed in the **Supplemental Material**. Full-length IntS1-14 expression plasmids were obtained from Eric J. Wagner, University of Rochester. Plasmids expressing IntS6 (pMtnA FLAG-IntS6 puro, Addgene #195076) or IntS12 (pMtnA FLAG-IntS12 puro, Addgene #195077) under the control of the Metallothionein A promoter were generated using the previously described pMT FLAG MCS puro plasmid.^26^ The reporter plasmid expressing eGFP downstream of the U4:39B snRNA gene was previously described.^41^ An analogous reporter expressing eGFP downstream of U5:34A snRNA (Hy_U5:34A eGFP SV40, Addgene #195064) was generated from Hy_pPepck1 eGFP SV40. Details of all cloning, including full plasmid sequences, are described in the **Supplemental Material**.

### Double-stranded RNAs

Double-stranded RNAs (dsRNAs) from the DRSC (*Drosophila* RNAi Screening Center) were generated by *in vitro* transcription (MEGAscript kit, Thermo Fisher Scientific AMB13345) of PCR templates containing the T7 promoter sequence on both ends as previously described in detail.^56^ Primer sequences are provided in **Table S4**.

### Generation of stable cell lines

To generate DL1 cells stably maintaining the inducible IntS6 or IntS12 transgenes, 2 x 10^6^ cells were first plated in complete media in 6-well dishes. After 24 h, 2 μg of pMtnA FLAG-IntS6 puro (Addgene #195076) or pMtnA FLAG-IntS12 puro (Addgene #195077) plasmid was transfected using Effectene (Qiagen 301427; 16 μL Enhancer and 30 μL Effectene Reagent). On the following day, 5 μg/mL puromycin was added to the media to select and maintain the cell population.

### Transient transfections, RNAi, and RNA isolation

To determine the effect of Integrator subunit over-expression on the output of the eGFP reporters, DL1 cells were seeded in 12-well plates (5 x 10^5^ cells per well) in complete Schneider’s *Drosophila* medium and cultured overnight. On the following day, 500 ng of plasmid DNA was transfected into each well using Effectene (4 μL of enhancer and 5 μL of Effectene reagent; Qiagen 301427). Unless otherwise noted, 400 ng of eGFP plasmid was transfected along with 100 ng of the Integrator/PP2A subunit expression plasmid. Total RNA was isolated ∼48 h later using TRIzol (Thermo Fisher Scientific 15596018) according to the manufacturer’s instructions.

To determine the effect of Integrator subunit down-regulation on the output of the eGFP reporters, 1 x 10^6^ DL1 cells were first bathed in 2 μg of dsRNAs in 12-well plates in 0.5 mL Schneider’s *Drosophila* medium without FBS, penicillin/streptomycin, or L-glutamine. After incubating cells at 25°C for 45 min, 1 mL of complete *Drosophila* medium was added and cells were grown at 25°C. After 24 h, Effectene (4 μL of enhancer and 5 μL of Effectene reagent; Qiagen 301427) was used to transfect 500 ng of the indicated eGFP plasmid. Total RNA was isolated ∼48 h later using TRIzol (Thermo Fisher Scientific 15596018) according to the manufacturer’s instructions. When examining reporters driven by the Metallothionein A promoter, a final concentration of 500 μM copper sulfate (Fisher BioReagents BP346-500) was added to cells for the last 14 h prior to RNA isolation.

### Northern blotting

Northern blots using 1.2% denaturing formaldehyde agarose gels, NorthernMax reagents (Thermo Fisher Scientific), and oligonucleotide probes were performed as previously described in detail.^57^ Oligonucleotide probe sequences are provided in **Table S4**. Blots were hybridized overnight at 42°C with oligonucleotide probes in ULTRAhyb-Oligo (Thermo Fisher Scientific AM8663), washed two times with 2x SSC, 0.5% SDS, and viewed with the Amersham Typhoon scanner (Cytiva) followed by quantification using ImageQuant (Cytiva).

### Antibody production

The C-terminal region of *Drosophila* IntS6 (amino acids 1035-1284) and the N-terminal region of *Drosophila* IntS8 (amino acids 1-308) were individually cloned into pRSFDuet-1 using the BamHI and HindIII restriction enzyme sites to generate pRSFDuet-1 IntS6 AA 1035-1284 (Addgene #196904) and pRSFDuet-1 IntS8 AA 1-308 (Addgene #196905), respectively. The C-terminal region of *Drosophila* IntS11 (amino acids 300-597) was cloned into pRSFDuet-1 using the SalI and HindIII restriction enzyme sites to generate pRSFDuet-1 IntS11 AA 300-597 (Addgene #199329). Details of cloning, including full plasmid sequences, are described in the **Supplemental Material**. Plasmids were transformed into BL21 Star (DE3) *E. coli* and grown in terrific broth media supplemented with 50 µg/mL kanamycin. Expression of the His-tagged proteins was induced at OD_600_ ∼0.8 by addition of 0.3 mM IPTG. IntS6, IntS8, and IntS11 cultures were incubated at 16°C for 20 h, 25°C for 7 h, and 16°C for 20 h, respectively, before cells were harvested.

For purification of His-tagged IntS6, the cell pellet was lysed by sonication in lysis buffer (50 mM Tris pH 8.0, 500 mM NaCl, 0.5 mM DTT, 25 mM imidazole, 1 mM PMSF, 100 µM leupeptin, 10 µM pepstatin A, and 1 mM benzamidine). Cell debris was removed by centrifugation at 20,000 x *g* for 15 min. The supernatant was filtered and loaded onto a Ni-column (Cytiva 17524801). The column was then washed with wash buffer (30 mM Tris pH 8.0, 300 mM NaCl, 50 mM imidazole) and eluted by gradient of imidazole to 400 mM. Fractions containing His-tagged IntS6 were pooled, concentrated, and loaded onto a Superdex 200 column in PBS. Fractions containing His-tagged IntS6 were pooled, concentrated, flash frozen, and stored at -80°C.

His-tagged IntS8 and IntS11 were purified under denaturing conditions. Cells were resuspended and incubated in lysis buffer (30 mM Tris pH 7.5, 300 mM NaCl, 1 mM DTT, 1 mM EDTA, 100 µg/mL lysozyme, 1 mM PMSF, 100 µM leupeptin, 10 µM pepstatin A, and 1 mM benzamidine) on ice for 30 min, and then sonicated for a total of 5 min. Sample was pelleted by centrifugation at 20,000 x *g* for 15 min and then resuspended in denaturing buffer (30 mM Tris pH 7.5, 300 mM NaCl, and 8 M urea) at 4°C for 30 min while stirring. Cell debris was removed by centrifugation at 20,000 x *g* for 15 min. The solubilized IntS8 protein was diluted with buffer to lower the urea concentration to 1 M, loaded onto a Ni-column (Cytiva 29051021), and eluted by adding elution buffer (30 mM Tris pH 7.5, 300 mM NaCl, and 500 mM imidazole). Fractions containing His-tagged IntS8 were pooled, dialyzed against PBS, concentrated, flash frozen, and stored at -80°C. The solubilized IntS11 protein in denaturing buffer was flash frozen and stored at -80°C.

Purified His-tagged IntS6 and IntS8 proteins in PBS were shipped to a commercial vendor (Cocalico Biologicals) and used to inoculate rabbits. His-tagged IntS11 was purified from SDS-PAGE gels and the cut bands were likewise used to inoculate rabbits. The reactivity and specificity of antisera was confirmed with Western blots using whole cell extracts from DL1 cells treated with dsRNAs targeting either IntS6, IntS8, or IntS11.

### Western blotting

Cells were pelleted at 5,000 x *g* for 1 min and then resuspended in RIPA buffer (150 mM NaCl, 1% Triton X-100, 50 mM Tris pH 7.5, 0.1% SDS, 0.5% sodium deoxycholate, and protease inhibitors [Roche 11836170001]). After 20 min incubation on ice, lysates were cleared at 21,000 x *g* for 20 min at 4°C. Protein concentrations were measured using a standard Bradford assay (Bio-Rad 5000006). Lysates containing 20 μg protein were then resolved on NuPAGE 4-12% Bis-Tris gels (Thermo Fisher Scientific NP0323) and transferred to PVDF membranes (Bio-Rad 1620177). Membranes were blocked with 10% nonfat milk in TBST for 1 h before incubation in primary antibody (diluted in 1x TBST) overnight at 4°C. Membranes were then washed with 1x TBST (4 x 10 min) followed by incubation in secondary antibody (diluted in 1x TBST) at room temperature for 1 h. Antibody incubation conditions are summarized in **Table S5**. Membranes were washed with 1x TBST (4 x 10 min) and processed using SuperSignal West Pico PLUS Chemiluminescent Substrate (Thermo Fisher Scientific PI34080).

### RT-qPCR

5 μg of total RNA (quantified by Nanodrop) was digested with TURBO DNase (Thermo Fisher Scientific AM2238) in a 20 μL reaction following the manufacturer’s protocol. Samples were then incubated at 75°C for 10 min in the presence of 15 mM EDTA. 1 μg of the digested RNA was reverse transcribed to cDNA in a 20 μL reaction using TaqMan Reverse Transcription Reagents (Thermo Fisher Scientific N8080234) with random hexamers following the manufacturer’s protocol except that 4 μL of 25 mM MgCl_2_ instead of 1.4 μL was used. RT-qPCR reactions were performed in 15 μL reactions that contained 1.5 μL of cDNA (diluted up to 10-fold in H_2_O), 7.5 μL 2x Power SYBR Green PCR Master Mix (Thermo Fisher Scientific 4368708), and 6 μL 1.5 μM gene-specific primer pairs. Primer sequences are provided in **Table S4**.

Using the QuantStudio 3 Real-Time PCR System (Thermo Fisher Scientific A28566) and clear plates (Thermo Fisher Scientific 4346907), the following cycling conditions were used: 95°C for 10 min, 40 amplification cycles of 95°C for 15 s followed by 60°C for 1 min, and a final melting cycle of 95°C for 10 s, 65°C for 1 min, and 97°C for 1 s. Subsequently, a melt curve was performed to verify that amplified products were a single discrete species. Threshold cycle (CT) values were calculated by the QuantStudio 3 system and relative transcript levels (compared to RpL32) were calculated using the 2^−ΔΔCT^ method. RT-qPCR reactions were performed using at least three independent biological replicates, with each replicate having two technical replicates.

### Immunoprecipitation

Parental DL1 cells and DL1 cells stably maintaining inducible FLAG-tagged IntS6 or IntS12 transgenes (DL1 pMtnA IntS6 and DL1 pMtnA IntS12, respectively) were grown in T75 flasks (4 flasks/each) for 3 d and 500 μM copper sulfate (Fisher BioReagents BP346-500) was added for the last 24 h. Cells were harvested by centrifugation at 340 x *g* for 5 min and then washed with ice-cold PBS. Cells were resuspended in hypotonic buffer (50 mM Tris pH 7.5, 10 mM KCl, 1 mM DTT, and protease inhibitor mix containing 1 mM PMSF, 100 µM leupeptin, 10 µM pepstatin A, and 1 mM benzamidine) and incubated for 20 min on ice. Cell suspension was then homogenized using a glass Dounce homogenizer. Nuclei were pelleted at 1000 x *g* for 10 min at 4°C and washed once with hypotonic buffer. Nuclei were incubated in lysis buffer (40 mM Tris pH 7.5, 300 mM NaCl, 1 mM DTT, 10% glycerol, 0.75% Triton X-100 and protease inhibitors) for 20 min on ice. Insoluble proteins and cell debris were removed by centrifugation at 21,000 x *g* for 30 min and the supernatant was passed through a 0.45 µm filter (Thermo Fisher Scientific F2513-14). The supernatant was diluted with dilution buffer (20 mM Tris pH 7.5, 10 mM NaCl, 1 mM DTT, 10% glycerol, and protease inhibitors) such that the final NaCl and Triton X-100 concentrations became 200 mM and 0.5%, respectively. The diluted supernatant was then incubated with pre-equilibrated anti-FLAG beads (Sigma A2220) while rotating for 2 h at 4°C. The beads were washed five times with wash buffer (20 mM Tris pH 7.5, 150 mM NaCl, 1 mM DTT, 10% glycerol, 0.25% Triton X-100, and protease inhibitors). Bound proteins were eluted with 0.1 M glycine pH 3.5, TCA precipitated, and resuspended in 1x loading dye containing 5 mM DTT.

### RNA-seq library generation

To determine the effect of Integrator subunit over-expression on the endogenous transcriptome, DL1, DL1 pMtnA IntS6, and DL1 pMtnA IntS12 cells were seeded in 6-well plates (2 x 10^6^ cells per well) in 2 mL complete Schneider’s *Drosophila* medium (with FBS, penicillin/streptomycin, and L-glutamine) and grown for 3 d. As indicated, a final concentration of 500 μ copper sulfate (Fisher BioReagents BP346-500) was added to cells for the last 24 h prior to RNA isolation using TRIzol (Thermo Fisher Scientific 15596018). Total RNA was isolated from three biological replicates, treated with DNase I, depleted of ribosomal RNAs, and strand-specific RNA-seq libraries were generated by Genewiz/Azenta Life Sciences. Libraries were sequenced using Illumina HiSeq, 2 x 150 bp configuration.

To determine the effect of Integrator/PP2A subunit down-regulation on the endogenous transcriptome, 5 x 10^5^ DL1 cells were bathed in 2 μg of dsRNAs in 12-well plates in 0.5 mL Schneider’s *Drosophila* medium without FBS, penicillin/streptomycin, or L-glutamine. After incubating cells at 25°C for 45 min, 1 mL of complete *Drosophila* medium was added and cells were grown at 25°C for 3 d. Total RNA was isolated using TRIzol (Thermo Fisher Scientific 15596018) according to the manufacturer’s instructions. RNAs were treated with DNase I, depleted of ribosomal RNAs, and strand-specific RNA-seq libraries were generated by Genewiz/Azenta Life Sciences. Libraries were sequenced using Illumina HiSeq, 2 x 150 bp configuration.

### RNA-seq analysis

For paired-end RNA-seq samples generated in this study, Trimmomatic^58^ (version 0.39; parameters: PE -threads 3 -phred33 TruSeq3PE-2.fa:2:30:8:true LEADING:3 TRAILING:3 SLIDINGWINDOW:4:15 MINLEN:30) was used for quality control of RNA-seq datasets, including removal of adaptor sequences and low-quality bases at both ends of reads. Reads were next aligned to the *D. melanogaster* dm6/BDGP6.22 reference genome using HISAT2^59^ (version 2.1.0, parameters: --no-softclip --score-min L,-16,0 --mp 7,7 --rfg 0,7 --rdg 0,7 --dta -k 1 --max-seeds 20). Mapping summaries of RNA-seq datasets are described in **Figures S3C and S5A**. Fragment counts were calculated per gene using featureCounts^60^ (version 2.0.1, parameters: -s 2 -p --fraction -O -T 16 -t exon -g gene_id) in a strand-specific manner based on the *D. melanogaster* Ensembl gene annotation (BDGP6.22.96). Differentially expressed genes (DEGs) were then identified using DESeq2^61^ (version: 1.26.0) under R 3.6.3 with a threshold of an adjusted *P* value < 0.001 and |log2(fold change, FC)| > 0.585. *Pearson* correlation coefficient (PCC) of FC of 96 up-regulated genes between IntS6 over-expression, IntS4 depletion, and IntS6 depletion were calculated under R 3.6.3. GO enrichment analysis (biological process) for the 107 genes up-regulated with IntS6 over-expression was performed and visualized with clusterProfiler^62^ (version 3.14.3, parameters: pvalueCutoff = 0.05, qvalueCutoff = 0.2).

Fragment counts mapped to 3 kb downstream of snRNA gene bodies were calculated using featureCounts^60^ (version 2.0.1, parameters: -s 2 -p --fraction -O -T 16 -t exon -g gene_id) in a strand-specific manner based on the *D. melanogaster* Ensembl gene annotation (BDGP6.22.96). DESeq2^61^ (version: 1.26.0) was then used to compare differences for each snRNA between treatments under R 3.6.3. Next, to compare the overall endogenous snRNA readthrough levels between treatments, readthrough transcription from a given snRNA was calculated by dividing RNA-seq fragment counts aligned to 3 kb downstream of the snRNA gene body with the fragment counts aligned to the mature snRNA. Statistical difference between treatments was assessed using *Wilcoxon* signed-rank test under R 3.6.3.

For single-end RNA-seq samples downloaded from GEO (GSE114467),^26^ Trimmomatic^58^ (version 0.39; parameters: SE -threads 5 -phred33 TruSeq3PE-2.fa:2:30:8:true LEADING:3 TRAILING:3 SLIDINGWINDOW:4:15 MINLEN:30) was used for quality control, including removal of adaptor sequences and low-quality bases. Reads were then aligned to *D. melanogaster* dm6/BDGP6.22 reference genome using HISAT2^59^ (version 2.1.0, parameters: -k 1 --max-seeds 2). The mapping summaries of RNA-seq datasets are described in **Figure S5F**. Fragment counts were calculated per gene using featureCounts^60^ (version 2.0.1, parameters: -s 1 -p --fraction -O -T 16 -t exon -g gene_id) and then normalized to FPKM (fragments per kilobase of gene per million fragments mapped).^63^

Lengths of all genes were extracted from the *D. melanogaster* Ensembl gene annotation (BDGP6.22.96).

### ChIP-seq analysis

Published ChIP-seq data for IntS1 and IntS12 in DL1 cells (3 replicates/each) were downloaded from GEO (GSE114467).^26^ Raw sequences were filtered and trimmed using Trimmomatic^58^ (version 0.39; parameters: PE -threads 3 -phred33 TruSeq3PE-2.fa:2:30:8:true LEADING:3 TRAILING:3 SLIDINGWINDOW:4:15 MINLEN:30). Reads were then aligned to the *D. melanogaster* dm6/BDGP6.22 reference genome using bowtie2^64^ (version 2.3.5) with default parameters. Multi-mapped reads were removed using sambamba^65^ (version 0.8.0, parameter: view -F “[XS] == null and not unmapped”). Mapping summaries of ChIP-seq datasets are described in **Figure S4A**. IntS1 and IntS12 ChIP-seq peaks were next called using MACS2^66^ (version 2.2.7.1, parameters: -g 142573017 -q 0.001). ChIP-seq peaks were annotated by ChIPseeker^67^ (version 1.22.1), with the TxDb object of *D. melanogaster* Ensembl gene annotation (BDGP6.22.96) generated using the makeTxDbFromGFF function in GenomicFeatures package (version 1.38.0).^68^ Genes closest to peaks and with peaks located within ±1 kb of gene bodies in all 3 replicates were defined as IntS1 or IntS12 binding genes. Finally, IntS1 binding genes and IntS12 binding genes were merged to define a set of 3,932 Integrator bound genes.

### Quantification and statistical analysis

For Northern blots and RT-qPCRs, statistical significance for comparison of means was assessed by one-way ANOVA using GraphPad Prism.

