## Supplemental Material for "IntS6 and the Integrator phosphatase module tune the efficiency of select premature transcription termination events"

**SUPPLEMENTAL FIGURE LEGENDS**

**Figure S1. Validation of Integrator subunit depletion and over-expression effects in *Drosophila* DL1 cells. Related to Figure 1.**

**(A, B)** To quantify the efficiency of dsRNA-mediated depletion of target gene expression, *Drosophila* DL1 cells were treated with dsRNAs for 3 d and then total RNA or protein was isolated. **(A)** RT-qPCR was used to quantify expression of IntS4, IntS6, mts, or Pp2A-29B mRNAs and data were normalized to expression of RpL32 mRNA. Data are shown as mean ± SD, *N* = 3. **(B)** Western blotting was performed using an antibody that recognizes the C-terminal region of *Drosophila* IntS6 (amino acids 1035-1284). * denotes non-specific band. α-tubulin was used as a loading control. Data are shown as mean ± SD, *N* = 3.

**(C)** Equal loading of RNA was verified for the Northern blots shown in **Figure 1B** by re-probing the membranes for RpL32 mRNA.

**(D)** DL1 cells were transfected with 500 ng of the indicated FLAG-tagged expression plasmids and total protein was harvested after 48 h. A plasmid containing a multi-cloning site (MCS) was used as a control. Western blot analysis using an antibody that recognizes FLAG was used to confirm expression of individual Integrator subunits. α-tubulin was used as a loading control.

**Figure S2. IntS6 over-expression is sufficient to inhibit Integrator activity at select protein-coding gene promoters. Related to Figure 1.**

**(A, B)** *Drosophila* DL1 cells (5 x 10^5^ cells per well) were seeded in 12-well dishes and co-transfected the next day with (i) a constant amount (300 ng) of pMtnA eGFP **(A)** or pUbi-p63e eGFP **(B)** reporter plasmid and (ii) variable amounts of plasmids that express FLAG-tagged IntS6, IntS12, or IntS8 from the Ubi-p63e promoter (0, 20, 40, 60, 80, 100, 150, or 200 ng). Empty vector (pUb 3xFLAG MCS) was added as needed so that 500 ng DNA was transfected in all samples. Total RNA was isolated ~40 h after transfection (500 μM CuSO_4_ was added for the last 14 h in **A**) and eGFP mRNA expression levels were analyzed by Northern blotting (20 μg/lane). Data were normalized to the (-) samples where only the eGFP plasmid was transfected and data are shown as mean ± SD, *N* = 4. (*) P < 0.05; n.s., not significant.

**(C)** Schematics of plasmids that express FLAG-tagged full-length (FL) or shortened isoforms of the 1284 amino acid *Drosophila* IntS6 open reading frame from the Ubi-p63e promoter. Characterized protein domains in IntS6 are noted in red.

**(D)** 500 ng of each *Drosophila* IntS6 expression plasmid was transfected into DL1 cells and total protein was harvested after 48 h. A plasmid containing a multi-cloning site (MCS) was used as a control. Western blot analysis using an antibody that recognizes FLAG was used to confirm expression of each IntS6 protein isoform. α-tubulin was used as a loading control.

**(E)** DL1 cells (5 x 10^5^ cells per well) were seeded in 12-well dishes and co-transfected the next day with 400 ng of pMtnA eGFP plasmid and 100 ng of the indicated IntS6 expression plasmid. Total RNA was isolated ~48 h after transfection (500 μM CuSO_4_ was added for the last 14 h) and eGFP mRNA expression levels were analyzed by Northern blotting (20 μg/lane). Data were normalized to the (-) samples where only the pMtnA eGFP plasmid was transfected and data are shown as mean ± SD, *N* = 3. (*) P < 0.05; n.s., not significant.

**(F)** 500 ng of plasmid that expresses FLAG-tagged *Drosophila* (d), human (h), or zebrafish (z) IntS6/IntS6-like (IntS6L) protein was transfected into DL1 cells and total protein was harvested after 48 h. A plasmid containing a multi-cloning site (MCS) was used as a control. Western blot analysis using an antibody that recognizes FLAG was used to confirm expression of each isoform. α-tubulin was used as a loading control.

**(G)** DL1 cells (5 x 10^5^ cells per well) were seeded in 12-well dishes and co-transfected the next day with 400 ng of the indicated eGFP plasmid and 100 ng of the indicated IntS6/IntS6L expression plasmid. Total RNA was isolated ~48 h after transfection (500 μM CuSO_4_ was added for the last 14 h only when measuring eGFP production from the MtnA promoter) and eGFP mRNA expression levels were analyzed by Northern blotting (20 μg/lane). Data were normalized to the (-) samples where only the eGFP plasmid was transfected and data are shown as mean ± SD, *N* = 3. (*) P < 0.05; n.s., not significant.

**Figure S3. Identification of endogenous differentially expressed genes upon IntS6 over-expression. Related to Figures 2 and 3.**

**(A)** The parental DL1 cells as well as stable cell lines that express FLAG-tagged IntS6 or IntS12 from the copper-inducible MtnA promoter were seeded in 12-well plates (5 x 10^5^ cells per well) and grown for 3 d. As indicated, a final concentration of 500 μM CuSO_4_ was added to cells for the last 24 h prior to harvesting total protein. Western blot analysis was then performed using antibodies that recognize FLAG, IntS6 (amino acids 1035-1284), IntS12, IntS8 (amino acids 1-308), or IntS11 (amino acids 300-597). * denotes non-specific band. α-tubulin was used as a loading control. Subunit expression data were normalized to the parental DL1 cells without CuSO_4_ treatment and are shown as mean ± SD, *N* = 3. (*) P < 0.05.

**(B)** To confirm the specificity of the IntS8, IntS11, and IntS12 antibodies, DL1 cells were treated with dsRNAs for 3 d and Western blots performed. * denotes non-specific band.

**(C)** Parental DL1 cells or DL1 cells stably maintaining IntS6 or IntS12 transgenes driven by the copper inducible MtnA promoter were grown for 3 d. As indicated, 500 μM CuSO_4_ was added for the last 24 h prior to total RNA isolation. rRNA depleted RNA-seq libraries were then generated, sequenced, and analyzed. Quality control and mapping statistics of the RNA-seq data to the *Drosophila* genome (dm6) are shown.

**(D)** Analysis pipeline used to identify genes that are both differentially expressed upon IntS6 over-expression and have IntS1/12 ChIP-seq signal ± 1 kb of the gene body (also see **Figure S4**).

**(E)** GO enrichment analysis (biological process) for the 107 genes up-regulated upon IntS6 over-expression was performed and visualized with clusterProfiler^62^ (parameters: pvalueCutoff = 0.05, qvalueCutoff = 0.2).

**(F)** Length distribution of different gene sets as per the *D. melanogaster* Ensembl gene annotation (BDGP6.22.96). The vertical line indicates the median gene length of each gene set.

**Figure S4. ChIP-seq analysis to define protein-coding genes bound by Integrator. Related to Figure 3.**

**(A)** Quality control and mapping statistics of input, IntS1, and IntS12 ChIP-seq data from DL1 cells (GSE114467)^26^ to the *Drosophila* genome (dm6).

**(B)** Analysis pipeline used to identify genes that have peaks of IntS1 or IntS12 ChIP-seq signal ± 1 kb of the gene body.

**(C)** Densities of ChIP-seq peaks ±2 kb of transcription start sites (TSS).

**(D)** Proportions of gene elements with mapped ChIP-seq peaks.

**(E)** Comparison of genes bound by IntS1 and IntS12 identified 1,698 genes that are bound by both subunits along with a total merged list of 3,932 genes bound by Integrator.

**Figure S5. Identification of differentially expressed genes upon Integrator subunit depletion. Related to Figure 3.**

**(A)** DL1 cells were treated with control (β-gal), IntS4, IntS6, mts, or Pp2A-29B dsRNAs for 3 d. Total RNA was isolated and rRNA depleted RNA-seq libraries were then generated, sequenced, and analyzed. Quality control and mapping statistics of the RNA-seq data to the *Drosophila* genome (dm6) are shown.
 **(B, C)** The magnitude of change in mRNA expression compared with statistical significance (*P*-value) is shown as a volcano plot. Threshold used to define IntS4 **(B)** or IntS6 **(C)** affected mRNAs was (|log2(fold change)| > 0.585 and adjusted *P* value < 0.001.
 **(D, E)** The correlation of fold change for 96 up-regulated genes between IntS6 overexpression, IntS4 depletion **(D)** and IntS6 depletion **(E)**. The grey area indicates the confidence interval of linear regression. cor: *Pearson* correlation coefficient. **(F)** Quality control and mapping statistics of published RNA-seq data (GSE114467)^26^ to the *Drosophila* genome (dm6). DL1 cells were treated for 60 h with control (β-gal) or IntS11 dsRNAs and rescued using a stably integrated transgene expressing eGFP, wild-type (WT) IntS11, or IntS11 with a mutation that disrupts endonuclease activity (E203Q).^26^  **(G)** Heat map shows relative expression levels of the 107 genes that are up-regulated upon IntS6 over-expression in the RNA-seq samples described in **F**.

**Figure S6. Changes in pre-mRNA levels upon IntS6 or IntS12 over-expression. Related to Figure 3.
(A, B)** Parental DL1 cells or DL1 cells stably maintaining IntS6 or IntS12 transgenes driven by the copper inducible MtnA promoter were grown for 3 d. 500 μM CuSO_4_ was added to each for the last 24 h prior to total RNA isolation from four independent biological replicates. Expression of the indicated pre-mRNAs for genes whose mRNAs were up-regulated **(A)** or unchanged (**B)** upon IntS6 over-expression was quantified using RT-qPCR. Data are shown as mean ± SD, *N* = 4. (*) P < 0.05.

**Figure S7. Additional examples of protein-coding genes that were differentially affected by modulation of Integrator subunit levels. Related to Figure 3.**

**(A-F)** UCSC genome browser tracks (top) and validation RT-qPCR experiments (bottom) for protein-coding loci that are **(A-D)** or are not **(E, F)** affected by IntS6 over-expression. IntS1 and IntS12 ChIP-seq profiles in DL1 cells (GSE114467) are shown in black. RNA-seq data generated from DL1 cells treated for 3 d with control (β-gal), IntS4, or IntS6 dsRNAs are shown in blue. RNA-seq data generated from parental DL1 cells or DL1 cells stably maintaining inducible IntS6 or IntS12 transgenes are shown in red. 500 μM CuSO_4_ was added for the last 24 h as indicated. Green arrow, transcription start site (TSS). (Bottom) Expression of the indicated mRNAs was quantified using RT-qPCR. Data are shown as mean ± SD, *N* = 3. (*) P < 0.05.

**SUPPLEMENTAL TABLES**

**Table S1. Quantification of endogenous snRNA readthrough upon over-expression or depletion of Integrator subunits. Related to Figures 2 and 5.**
Comparisons of normalized RNA-seq fragment counts aligned to 3 kb downstream of each endogenous snRNA gene body in DL1 cells between treatments (depletion of IntS4, IntS6, mts, or Pp2A-29B using RNAi, or over-expression of IntS6 or IntS12). Statistical difference for each snRNA readthrough was assessed by DESeq2.^61^

**Table S2. Endogenous protein-coding and long non-coding RNA genes up-regulated upon IntS6 over-expression. Related to Figure 3.**
List of protein-coding and long non-coding genes up-regulated in DL1 cells upon IntS6 over-expression when compared to parental DL1 cells or DL1 cells over-expressing IntS12. All cell lines were grown in the presence of Cu^2+^ for 24 h prior to total RNA isolation and RNA-seq library generation. Threshold used to define differentially expressed genes was |log2(fold change)| > 0.585 and adjusted *P* < 0.001. An overlapping set of 107 genes was up-regulated upon IntS6 over-expression.

**Table S3. Endogenous protein-coding genes differentially expressed upon depletion of IntS4, IntS6, mts, or Pp2A-29B. Related to Figures 2 and 5.**
List of genes that were up- or down-regulated in *Drosophila* DL1 cells after RNAi depletion of the indicated factor as determined by RNA-seq. Threshold used to define differentially expressed genes was |log2(fold change)| > 0.585 and adjusted *P* < 0.001.

**Table S4. Oligonucleotide sequences. Related to Figures 1, 2, 3, 4, and 6.**
The oligonucleotide sequences used for Northern blots, RT-qPCR, and dsRNA synthesis are provided.

**Table S5. Antibodies for Western blots. Related to Figures 4 and 6.**
For each antibody used in this study, the source and experimental conditions are provided.

**SUPPLEMENTAL METHODS**

**Expression plasmid construction**

The following *Drosophila* expression plasmids were generated from the previously published **Hy_pPepck1 eGFP SV40** plasmid (Addgene #132644; Tatomer et al. (2019) *Genes Dev*, 33, 1525-1538):

**Hy_pRnrS eGFP SV40	(Addgene #195062)**

**Hy_pAna eGFP SV40	(Addgene #195063)**

**Hy_U5:34A eGFP SV40	(Addgene #195064)**

The Pepck1 promoter (marked in gray) drives expression of eGFP (marked in green) followed by the SV40 polyadenylation signal (marked in yellow). The full sequence of **Hy_pPepck1 eGFP SV40** is as follows:

GGTTAATAAGGAATATTTGATGTATAGTGCCTTGACTAGAGATCATAATCAGCCATACCACATTTGTAGAGGTTTTACTTGCTTTAAAAAACCTCCCACACCTCCCCCTGAACCTGAAACATAAAATGAATGCAATTGTTGTTGTTAACTTGTTTATTGCAGCTTATAATGGTTACAAATAAAGCAATAGCATCACAAATTTCACAAATAAAGCATTTTTTTCACTGCATTCTAGTTGTGGTTTGTCCAAACTCATCAATGTATCTTATCATGTCTGGATTAATTAAAAAAACAGAGCCATATTTACGTGCAACTTTCCCGACTGCTTCTGTTGAACTTTTTCTTCGAGTACCCAGAGGATTCTGGGTCTTCTGGTAAACTGGATTCGGGGTACAAAGTCACACCACCGGCAGCACCTTCATTTCTCCTTCCCCAAAGACAAAAACAAGAGTGGCCACATGACAACCATCCGGGCCATGCTATTCTTGGTGCGCCCCCAGAGTCACATGAATGTGAATACTGGCCTCCAAGGCGGCCATGGCCAATTGGGGAGGCCCCCGGCTCATCTGGATGCCGCGAGGGGGGCGGATTAGCATACAAATTTTCCGTCACCGATTCTACAAATAAGCGAAAGTTTTTTTCGATTTGGCAAATGTCCAAAAACGAGAGGCCAACACATGTGGTCAAACAGGCGTAAGTGGGCGGGCGCGTGTGATGTGGCCCACCATTCGATGTTGCACCAATGTAGCTATATTTGTGGCCAATGCAGCTATAGACACATCCAGAATTTACCCAGCTGCAGGGGACAACAACTGCTGTTGCCTCAGCTGTCCGTGAACTGGTTTCTATATTCGCAGCGGCCACATTTGTATCGCATAAAATCATAAATAGAATAATTAAATTAACAGCGCGAAAGTGGCAAATATCAGGGTGTTCAAAATAACAACAAACGCTTATCAATGCAAACGCACTAGAAAACGCTCTCAGCGCCACATGAAAACAACACTCTGCTGTTGCCATGTGGGGACATACCCCATATACGTATCTGTATGCCCATGCCCCCCATTCCGTAGACCCCCCCCCCCACCCACAACCAACTCATCGTAGCTGCGCCGCCTGTGAAAAACCAAAACAATCGCCGAGCCTCCGCCGGCAGCCAATCAGCTCGTCCGGAGTTTGACCCTCTCACTCGGCGAGGCGATCGTAAAACCCGATTTGCGTGGCGATCGGCCGACTTGTTCTAGTCGCCTCGATGGCCAGGCGTCGCCGGCTGGGTATAAAAGGCCCCGGATCGGAGCAGTTGGCATCAGTTACTCGTGGCCAGAGTAACTCGAGGTCGACGGTATCGATAAGCTTGATATCACCATGGTGAGCAAGGGCGAGGAGCTGTTCACCGGGGTGGTGCCCATCCTGGTCGAGCTGGACGGCGACGTAAACGGCCACAAGTTCAGCGTGTCCGGCGAGGGCGAGGGCGATGCCACCTACGGCAAGCTGACCCTGAAGTTCATCTGCACCACCGGCAAGCTGCCCGTGCCCTGGCCCACCCTCGTGACCACCCTGACCTACGGCGTGCAGTGCTTCAGCCGCTACCCCGACCACATGAAGCAGCACGACTTCTTCAAGTCCGCCATGCCCGAAGGCTACGTCCAGGAGCGCACCATCTTCTTCAAGGACGACGGCAACTACAAGACCCGCGCCGAGGTGAAGTTCGAGGGCGACACCCTGGTGAACCGCATCGAGCTGAAGGGCATCGACTTCAAGGAGGACGGCAACATCCTGGGGCACAAGCTGGAGTACAACTACAACAGCCACAACGTCTATATCATGGCCGACAAGCAGAAGAACGGCATCAAGGTGAACTTCAAGATCCGCCACAACATCGAGGACGGCAGCGTGCAGCTCGCCGACCACTACCAGCAGAACACCCCCATCGGCGACGGCCCCGTGCTGCTGCCCGACAACCACTACCTGAGCACCCAGTCCGCCCTGAGCAAAGACCCCAACGAGAAGCGCGATCACATGGTCCTGCTGGAGTTCGTGACCGCCGCCGGGATCACTCTCGGCATGGACGAGCTGTACAAGTAAGCGGCCGCAACTTGTTTATTGCAGCTTATAATGGTTACAAATAAAGCAATAGCATCACAAATTTCACAAATAAAGCATTTTTTTCACTGCATTCTAGTTGTGGTTTGTCCAAACTCATCAATGTATCTTAACTAGTGGATCCACTAGGGGCCGCCGACGCGAGGCTGGATGGCCTTCCCCATTATGATTCTTCTCGCTTCCGGCGGCATCGGGATGCCCGCGTTGCAGGCCATGCTGTCCAGGCAGGTAGATGACGACCATCAGGGACAGCTTCAAGGATCGCTCGCGGCTCTTACCAGCCTAACTTCGATCATTGGACCGCTGATCGTCACGGCGATTTATGCCGCCTCGGCGAGCACATGGAACGGGTTGGCATGGATTGTAGGCGCCGCCCTATACCTTGTCTGCCTCCCCGCGTTGCGTCGCGGTGCATGGAGCCGGGCCACCTCGACCTGAATGGAAGCCGGCGGCACCTCGCTAACGGATTCACCACTCCAAGAATTGGAGCCAATCAATTCTTGCGGAGAACTGTGAATGCGCAAACCAACCCTTGGCAGAACATATCCATCGCGTCCGCCATCTCCAGCAGCCGCACGCGGCGCATCTCGGGCAGCGTTGGGTCCTGGCCACGGGTGCGCATGATCGTGCTCCTGTCGTTGAGGACCCGGCTAGGCTGGCGGGGTTGCCTTACTGGTTAGCAGAATGAATCACCGATACGCGAGCGAACGTGAAGCGACTGCTGCTGCAAAACGTCTGCGACCTGAGCAACAACATGAATGGTCTTCGGTTTCCGTGTTTCGTAAAGTCTGGAAACGCGGAAGTCAGCGCTCTTCCGCTTCCTCGCTCACTGACTCGCTGCGCTCGGTCGTTCGGCTGCGGCGAGCGGTATCAGCTCACTCAAAGGCGGTAATACGGTTATCCACAGAATCAGGGGATAACGCAGGAAAGAACATGTGAGCAAAAGGCCAGCAAAAGGCCAGGAACCGTAAAAAGGCCGCGTTGCTGGCGTTTTTCCATAGGCTCCGCCCCCCTGACGAGCATCACAAAAATCGACGCTCAAGTCAGAGGTGGCGAAACCCGACAGGACTATAAAGATACCAGGCGTTTCCCCCTGGAAGCTCCCTCGTGCGCTCTCCTGTTCCGACCCTGCCGCTTACCGGATACCTGTCCGCCTTTCTCCCTTCGGGAAGCGTGGCGCTTTCTCATAGCTCACGCTGTAGGTATCTCAGTTCGGTGTAGGTCGTTCGCTCCAAGCTGGGCTGTGTGCACGAACCCCCCGTTCAGCCCGACCGCTGCGCCTTATCCGGTAACTATCGTCTTGAGTCCAACCCGGTAAGACACGACTTATCGCCACTGGCAGCAGCCACTGGTAACAGGATTAGCAGAGCGAGGTATGTAGGCGGTGCTACAGAGTTCTTGAAGTGGTGGCCTAACTACGGCTACACTAGAAGGACAGTATTTGGTATCTGCGCTCTGCTGAAGCCAGTTACCTTCGGAAAAAGAGTTGGTAGCTCTTGATCCGGCAAACAAACCACCGCTGGTAGCGGTGGTTTTTTTGTTTGCAAGCAGCAGATTACGCGCAGAAAAAAAGGATCTCAAGAAGATCCTTTGATCTTTTCTACGGGGTCTGACGCTCAGTGGAACGAAAACTCACGTTAAGGGATTTTGGTCATGAGATTATCAAAAAGGATCTTCACCTAGATCCTTTTAAATTAAAAATGAAGTTTTAAATCAATCTAAAGTATATATGAGTAAACTTGGTCTGACAGTTACCAATGCTTAATCAGTGAGGCACCTATCTCAGCGATCTGTCTATTTCGTTCATCCATAGTTGCCTGACTCCCCGTCGTGTAGATAACTACGATACGGGAGGGCTTACCATCTGGCCCCAGTGCTGCAATGATACCGCGAGACCCACGCTCACCGGCTCCAGATTTATCAGCAATAAACCAGCCAGCCGGAAGGGCCGAGCGCAGAAGTGGTCCTGCAACTTTATCCGCCTCCATCCAGTCTATTAATTGTTGCCGGGAAGCTAGAGTAAGTAGTTCGCCAGTTAATAGTTTGCGCAACGTTGTTGCCATTGCTGCAGGCATCGTGGTGTCACGCTCGTCGTTTGGTATGGCTTCATTCAGCTCCGGTTCCCAACGATCAAGGCGAGTTACATGATCCCCCATGTTGTGCAAAAAAGCGGTTAGCTCCTTCGGTCCTCCGATCGTTGTCAGAAGTAAGTTGGCCGCAGTGTTATCACTCATGGTTATGGCAGCACTGCATAATTCTCTTACTGTCATGCCATCCGTAAGATGCTTTTCTGTGACTGGTGAGTACTCAACCAAGTCATTCTGAGAATAGTGTATGCGGCGACCGAGTTGCTCTTGCCCGGCGTCAACACGGGATAATACCGCGCCACATAGCAGAACTTTAAAAGTGCTCATCATTGGAAAACGTTCTTCGGGGCGAAAACTCTCAAGGATCTTACCGCTGTTGAGATCCAGTTCGATGTAACCCACTCGTGCACCCAACTGATCTTCAGCATCTTTTACTTTCACCAGCGTTTCTGGGTGAGCAAAAACAGGAAGGCAAAATGCCGCAAAAAAGGGAATAAGGGCGACACGGAAATGTTGAATACTCATACTCTTCCTTTTTCAATATTATTGAAGCATTTATCAGGGTTATTGTCTCATGAGCGGATACATATTTGAATGTATTTAGAAAAATAAACAAATAGGGGTTCCGCGCACATTTCCCCGAAAAGTGCCACCTGACGTCTAAGAAACCATTATTATCATGACATTAACCTATAAAAATAGGCGTATCACGAGGCCCTTTCGTCTTCAAGAATTCACATTTGTACGAATTTTTTTTTTATCAAAAGTTCGAGTTTTTCACCAATTTCCTCATCAACCGAGCAAGGCAAACGGCTTTGAATAATATGGTGTTATATATACATATATCAAATCGCTGCTGACTGCGTGATTGATGGCCCCAAGATTACATATTATCGAATCAGGATTCAGAAGGAGATCAATGTCAAATGCGGACAGGAACATGAAAGACGCCTGTTATGCGCAATTAAAAATTTGGGTTTAATTGCTGTGGAAACTGTTGTTGGCGGCATCTTAAGTTCCTGTTTAACAACATCAACTACTTATGTACGTAGAAGCGTTTAAGCCATTTGCATACAGATGAGAACTGGCTTTTGTGCTAATCAGTCAAGATGACTCCGATGATGATGACTCATTACCTGACCAGTTTTCGCTGCTTTCTTTTCAACAACTACTTGTATATGTATTGTATCCAATAGCAATACATTGAATTTCCATGGTCTAGTCACGTATTATCATTTAATTGACACCAAGTCGTGTTATTGTTGAGCTATCGAGTTCAGCTCAAACATTTCTTATTCCCATGAATAAGCCGGCAAAAATATGCAATCTATGAAAGTTAATATAAGCAAACCTTACTTTGACTCAATACCAATGCACTTTGTGTCGATAGGTTCACGCAATTGAGGCGATTATTCCGATAACCCAAGCGATTGACTGTTCCCGTTTCGATTCCAATTGAAATTTGGAAATGTACAATAGTTTTGCTATATGCTGTCAAGTACGCTCTTATCTTCTCTGGGTTTTCTTCAGAGTTTCGAAACGCTTCTTCTTTTTTTTGTTTTTTTTTTTTTGGAATCTCGTATTTTGGAAGGGGCTCCCCTCTGGAATTTGTTACACTGTCGTTATCATTGCGAACAAGCGGCCCGAAGCTATCAGCGACTTTAACATTTACAATGCACTTTTTTACGACCAATTAAATGTACATTTTCCTTTCTTCGCCCGTTGATAAGCGAACGCGATGTGGCGCAGGCAATGTGTTGCTCTTGCGACACAAACGCAATCAAAATGGATTCAATTTCGCTTTTTCCCAGTGAAACGAAGAACGAACCGACCATCATGATATGCTCCTCTGCATGTTGCGTATTGAATCAATGACAATTTCAATTAAGCCGCCCGTTCGTCATGCGTTTTCGTGCGCTTCGAAATGCTGATAACGCTGCTGTCCTCCAACTGCTTTGCATGTGGACACAATTCCATTTATTTAATTCTTTTATTTGGATCGGTTAAATTAAAAAGCGCCTTGTTACGCATTTAACGTTGTTTCCGGTGCGTGGTGGTTTCATGCTTCTGGGAACGGCAAATGGGTTTAGGATTGGGAACCCCTCATCATCTGTTGGAATATACTATTCAACCTACAAAAGTAACGTTAAACAACACTACTTTATATTTGATATGAATGGCCACACCTTTTATGCCATAAAACATATTGTAAGAGAATACCACTCTTTTTATTCCTTCTTTCCTTCTTGTACGTTTTTTGCTGTAAGTAGGTCGTGGTGCTGGTGTTGCAGTTGAAATAACTTAAAATATAAATCATAAAACTCAAACATAAACTTGACTATTTATTTATTTATTAAGAAAGGAAATATAAATTATAAATTACAACAGGTTATGGACCTGCAGCCAAGCTTGGCGCTCGTCCGGGGGCAATGAGATATGAAAAAGCCTGAACTCACCGCGACGTCTGTCGAGAAGTTTCTGATCGAAAAGTTCGACAGCGTCTCCGACCTGATGCAGCTCTCGGAGGGCGAAGAATCTCGTGCTTTCAGCTTCGATGTAGGAGGGCGTGGATATGTCCTGCGGGTAAATAGCTGCGCCGATGGTTTCTACAAAGATCGTTATGTTTATCGGCACTTTGCATCGGCCGCGCTCCCGATTCCGGAAGTGCTTGACATTGGGGAATTCAGCGAGAGCCTGACCTATTGCATCTCCCGCCGTGCACAGGGTGTCACGTTGCAAGACCTGCCTGAAACCGAACTGCCCGCTGTTCTGCAGCCGGTCGCGGAGGCCATGGATGCGATCGCTGCGGCCGATCTTAGCCAGACGAGCGGGTTCGGCCCATTCGGACCGCAAGGAATCGGTCAATACACTACATGGCGTGATTTCATATGCGCGATTGCTGATCCCCATGTGTATCACTGGCAAACTGTGATGGACGACACCGTCAGTGCGTCCGTCGCGCAGGCTCTCGATGAGCTGATGCTTTGGGCCGAGGACTGCCCCGAAGTCCGGCACCTCGTGCACGCGGATTTCGGCTCCAACAATGTCCTGACGGACAATGGCCGCATAACAGCGGTCATTGACTGGAGCGAGGCGATGTTCGGGGATTCCCAATACGAGGTCGCCAACATCTTCTTCTGGAGGCCGTGGTTGGCTTGTATGGAGCAGCAGACGCGCTACTTCGAGCGGAGGCATCCGGAGCTTGCAGGATCGCCGCGGCTCCGGGCGTATATGCTCCGCATTGGTCTTGACCAACTCTATCAGAGCTTGGTTGACGGCAATTTCGATGATGCAGCTTGGGCGCAGGGTCGATGCGACGCAATCGTCCGATCCGGAGCCGGGACTGTCGGGCGTACACAAATCGCCCGCAGAAGCGCGGCCGTCTGGACCGATGGCTGTGTAGAAGTACTCGCCGATAGTGGAAACCGACGCCCCAGCACTCGTCCGAGGGCAAAGGAATAGAGTAGATGCCGACCGAACAAGAGCTGATTTCGAGAACGCCTCAGCCAGCAACTCGCGCGAGCCTAGCAAGGCAAATGCGAGAGAACGGCCTTACGCTTGGTGGCACAGTTCTCGTCCACAGTTCGCTAAGCTCGCTCGGCTGGGTCGCGGGAGGGCCGGTCGCAGTGATTCAGGCCCTTCTGGATTGTGTTGGTCCCCAGGGCACGATTGTCATGCCCACGCACTCGGGTGATCTGACTGATCCCGCAGATTGGAGATCGCCGCCCGTGCCTGCCGATTGGGTGCAGATCTTTGTGAAGGAACCTTACTTCTGTGGTGTGACATAATTGGACAAACTACCTACAGAGATTTAAAGCTCTAAGGTAAATATAAAATTTTTAAGTGTATAATGTGTTAAACTACTGATTCTAATTGTTTGTGTATTTTAGATTCCAACCTATGGAACTGATGAATGGGAGCAGTGGTGGAATGCCTTTAATGAGGAAAACCTGTTTTGCTCAGAAGAAATGCCATCTAGTGATGATGAGGCTACTGCTGACTCTCAACATTCTACTCCTCCAAAAAAGAAGAGAAAGGTAGAAGACCCCAAGGACTTTCCTTCAGAATTGCTAAGTTTTTTGAGTCATGCTGTGTTTAGTAATAGAACTCTTGCTTGCTTTGCTATTTACACCACAAAGGAAAAAGCTGCACTGCTATACAAGAAAATTATGGAAAAATATTCTGTAACCTTTATAAGTAGGCATAACAGTTATAATCATAACATACTGTTTTTTCTTACTCCACACAGGCATAGAGTGTCTGCTATTAATAACTATGCTCAAAAATTGTGTACCTTTAGCTTTTTAATTTGTAAAGG

To generate **Hy_pRnrS eGFP SV40**, the following sequence was inserted between the PacI and XhoI sites:

CGATTAAACGTACTTATCGATAAATATCAACAGCCAGCTGTTATGTTATTGTACTGTTATTTACATGATTCGCTGTAATGGAATTCATTCAAAGTTGTTGAATTGAGTATGTTAGCACCTGTTGCATTTCCAAATAATTCGAGATTTCCAAAAAAAAAATTAATTTTAAGTCAATGTCCTTGGAAAAAAATTTTAACTTTGTTTCACCTTGATTTTTTATCTAATTGTAGCAAGAGGACCAAATTTTTTCACTCAAAACTAATTAAGTGGCGTTTCATCAGCTGTTTTCTGGATTAGTCTAGGGTTGTCTCATTGCATGAAATATCGATGATAAAAAAATTTCAAAATTTATTTAGTATTTGAAACTATTAATATTAATATTTTTCAAGTGACAAGCTGGGAAGCTAAACATAAAATTGTGCAGTAAGGATTCGATTTATGGTTTAAGAAAAGAAAACTACCACCCCATAATTGCATTAGATTTACCCTAAATTTATAAAAAGTGAATTGACGCACTCGACAGCCCTGATTTTCCCATAGTTTTCCCATCACCAAAAATGGCGGCAAATCGAAACAGTTTGCCGCCCGGCATAAAACCCAATGTAGCTGTATTTCCCAGATCATTTGCCACACAACTTTCAAACTGTACACTTAATACACGTCGTGGTTTAAGTGAATTTTACCAGAGAATCAGAGAAGCGCCCCTACCTGCTAATAATAATCCATTCAAAAACATCTCAA

To generate **Hy_pAna eGFP SV40**, the following sequence was inserted between the PacI and XhoI sites:

ATATATCTTGTTTTCATATTCCCCCCACACCACACCACACCACACCACCACCCCACCTACCGGCGGGATTATGAATCTGGGGGCATTTGATGCGATTGCAATTTTACATAACACATATCACTCGACGTCTGGAGAAGCCCGCCCCTGCGACCCCACTCACCACTCTATTCAGAACTTAAACGGTGTGTGTAGACACATCACATGAATGGGCAGCGGGAAGAAGCCGAATCCAGACTTTATAAAGTGCTGCGAATTATACAATCATTTTGAAATTCAATTTGGTCCAGACGTCGGCGGAGTGCTGGAAAAGATGTGTTTATATGTTGCAAATAAATAAATAAATCAGGAACGTCCTTGAAGTCTGGCTGTTTGGATTTCCGTATGCGAATGGAATGAATAATATTCGAGGCTTGTGCTATTCCTCATGCGTAGTACCAGAAGTGGGCCAAAGGATGGGCTTTCTGGGGTTAAGCGCCAAATGCCCAAAGCCGTGCCAGGCACCGCACGCACCTCTAATAGGATATATCCGTCCAATTCACCTTCCAATTGATGGACTACGCACTTTCGGGCCTATCTCGTTGTCAGATGATTAGTCCGCACGCACAAATGCCGGCGACAGCTGAAAGTCAAGAGCAAAGGCATCTTGCCCCATCGTGCCCCAAAAAATAACAGCAACCGATAACGGACATAACTGGCGTATCAAAACAATTTGATTAGGTCATACTAATTGTGCGCTTGCGAGGAAAATCACTTAGGTTAATCTAATCGCCCACGCACGTAGTAATTGAATAAATAGTGCCCGTCTTTACGGACATTCGAACGAATCCGATGGGTGCAAAGCCGTCCAAAAGATTGACACAGGTTGTTCATAATGTTTGAAGTGCCCCGAGTGTTTGGAGTAATAGATAGATAGATTGATATGCCTAGCAGTTTTCGGTTGATGGCCCTGTTTGATAGCAGCCACTTGAAAGGAAGTCGCCAAACAAACCTAGGTTAGGGCCGGAGTATCGCTTTCGGCCCTAAAGATAAGCCCCATCTCAGGCGCGCAGGCACAGGCCAAAGATACAGATACACCCATACAGCCGCGTTTCGTACGTATCGCACGCACGGCCACTGTAGAGCACCACGTCGACGTCAGCAGCGCCGGCAGATCCTCAAGCGGCGTTGATTGGTCGGCCACCGTGTATCCGAGCGCCAGCCGAACCGCCAGCCGAAGGCCCCAAGCGCTACACTGCAAACTTAGTTGGGGGGTTTTGAGTGGCGAACAAATGCGATAATAATATATAATGTAAAAAAACTAGTATAAAGAACTAAGCTAAGATTTAATGTACGCTTTCAAGCTCAGGTCACCTTGAATAGTACCTTTCAATCTCAATATTATTATATAATATTAATCCAGTAGTAAAATATTTATTTTGTGCCAATTTATTTAAGTAAGCCTTTTATTATATAGATAAATGTTATGTTAAGATATTACATGTATTATTTAATGTAATTTTATTAAGATATTAATAGTTTGCAATCAGCTAAAAGTAAGTTGTTCTTTTTAAGGCGGTCGCATTTTCTCACAGTGCCGGCCGCAACGCAGCGTTTACGACCATAACAAAGTAGATATGCCCGGCACAACCAAATTTTTCTTATTTAAGACTAAAAAACCGACTCGGGCAAATCACTTCATACTTGGTTCTTAGAGCGAACACATCGAGAAGCTGAACACAACAGCTACAAAAATTAACAAATTCGCATGTCCGTATCCAGCTGAAGTGTAAAAGATAGCATTTTAATTGCTGGCAAAAATGCAGTGAACAAACGTTGAATTTAAGTGAAGAAGAAAACTACAACTATTTCCGTGTGAAAACATTTAGTTTGGCAACAGACCAAGAGCAGAACACAAAA

To generate **Hy_U5:34A eGFP SV40**, the following sequence was inserted between the PacI and XhoI sites:

AACTTGAAACGGAAAACGAAAAGGGTCTCGATGGGCAAACGGCCAGAACCATTTGGGAATTATAAATGTGAAAGTAGAAAGTTATCGTACTTTCTATTGCATATTTAAGGGGTTTATGGTTCATATACTACAGACATTCCCGTTCCAGGTAGCCACATGATATAATATGACTAAGATCCCTACCATAAACATACTTTAGGGGTTGTTTGCGTTTTACAATCATTATGATTCCCAACATGTTCAAGCTCGTTCTAAATGATCGGGCTGATACTTTTCGTTGTAAATTTACTCTGGTTTCTCTTCAATTGTCGAATAAATCTTTCGCCTTTTACTAAAGATTTCCGTGGAGAGGAACACTCTAATGAGTCTAAAATAATCTTTTGTAGTGCCCGGCGACTTCGGTAGCTGGGCCATAAGGAATTCATTGTAAAGTTTTTTGCTTATTATTATAAGGTCTGGTGGTCATATACATGGACTACTCTACTAAGGTTACCAGAAGAGTGATGAAATTTTACTCGAATCGAGTTTGTACCGTAGACGGCGCTAAATGTGCAATAAATCAGCTTATTATATTGTGCAGTCAATTATATATTCTTAATGTCTATAGTGACTG

The following *Drosophila* expression plasmids were generated from the previously published **pUb 3xFLAG MCS** plasmid (Chen et al. (2012) *RNA*, 18, 2148-2156):

**pUb FLAG-Mts	(Addgene #195065)**

**pUb FLAG-Pp2A-29B	(Addgene #195066)**

**pUb FLAG-Cka	(Addgene #195067)**

**pUb FLAG-tws	(Addgene #195068)**

**pUb FLAG-wdb	(Addgene #195069)**

**pUb FLAG-wrd	(Addgene #195070)**

**pUb FLAG-IntS6 AA 1-400	(Addgene #195071)**

**pUb FLAG-IntS6 AA 1-600	(Addgene #195072)**

**pUb FLAG-IntS6 AA 101-1284	(Addgene #195073)**

**pUb FLAG-IntS6 AA 1197-1284	(Addgene #195074)**

**pUb FLAG-IntS6 AA 101-1200	(Addgene #195075)**

**pUb FLAG-Human IntS6	(Addgene #198408)**

**pUb FLAG-Human IntS6L	(Addgene #198409)**

**pUb FLAG-Zebrafish IntS6	(Addgene #198410)**

**pUb FLAG-Zebrafish IntS6L	(Addgene #198411)**

The Ubi-p63e promoter (pUb; marked in gray) drives expression of an N-terminal 3x FLAG tag (marked in blue). Start codon is marked in green. The full sequence of **pUb 3xFLAG MCS** is as follows:

GACGAAAGGGCCTCGTGATACGCCTATTTTTATAGGTTAATGTCATGATAATAATGGTTTCTTAGACGTCAGGTGGCACTTTTCGGGGAAATGTGCGCGGAACCCCTATTTGTTTATTTTTCTAAATACATTCAAATATGTATCCGCTCATGAGACAATAACCCTGATAAATGCTTCAATAATATTGAAAAAGGAAGAGTATGAGTATTCAACATTTCCGTGTCGCCCTTATTCCCTTTTTTGCGGCATTTTGCCTTCCTGTTTTTGCTCACCCAGAAACGCTGGTGAAAGTAAAAGATGCTGAAGATCAGTTGGGTGCACGAGTGGGTTACATCGAACTGGATCTCAACAGCGGTAAGATCCTTGAGAGTTTTCGCCCCGAAGAACGTTTTCCAATGATGAGCACTTTTAAAGTTCTGCTATGTGGCGCGGTATTATCCCGTATTGACGCCGGGCAAGAGCAACTCGGTCGCCGCATACACTATTCTCAGAATGACTTGGTTGAGTACTCACCAGTCACAGAAAAGCATCTTACGGATGGCATGACAGTAAGAGAATTATGCAGTGCTGCCATAACCATGAGTGATAACACTGCGGCCAACTTACTTCTGACAACGATCGGAGGACCGAAGGAGCTAACCGCTTTTTTGCACAACATGGGGGATCATGTAACTCGCCTTGATCGTTGGGAACCGGAGCTGAATGAAGCCATACCAAACGACGAGCGTGACACCACGATGCCTGTAGCAATGGCAACAACGTTGCGCAAACTATTAACTGGCGAACTACTTACTCTAGCTTCCCGGCAACAATTAATAGACTGGATGGAGGCGGATAAAGTTGCAGGACCACTTCTGCGCTCGGCCCTTCCGGCTGGCTGGTTTATTGCTGATAAATCTGGAGCCGGTGAGCGTGGGTCTCGCGGTATCATTGCAGCACTGGGGCCAGATGGTAAGCCCTCCCGTATCGTAGTTATCTACACGACGGGGAGTCAGGCAACTATGGATGAACGAAATAGACAGATCGCTGAGATAGGTGCCTCACTGATTAAGCATTGGTAACTGTCAGACCAAGTTTACTCATATATACTTTAGATTGATTTAAAACTTCATTTTTAATTTAAAAGGATCTAGGTGAAGATCCTTTTTGATAATCTCATGACCAAAATCCCTTAACGTGAGTTTTCGTTCCACTGAGCGTCAGACCCCGTAGAAAAGATCAAAGGATCTTCTTGAGATCCTTTTTTTCTGCGCGTAATCTGCTGCTTGCAAACAAAAAAACCACCGCTACCAGCGGTGGTTTGTTTGCCGGATCAAGAGCTACCAACTCTTTTTCCGAAGGTAACTGGCTTCAGCAGAGCGCAGATACCAAATACTGTTCTTCTAGTGTAGCCGTAGTTAGGCCACCACTTCAAGAACTCTGTAGCACCGCCTACATACCTCGCTCTGCTAATCCTGTTACCAGTGGCTGCTGCCAGTGGCGATAAGTCGTGTCTTACCGGGTTGGACTCAAGACGATAGTTACCGGATAAGGCGCAGCGGTCGGGCTGAACGGGGGGTTCGTGCACACAGCCCAGCTTGGAGCGAACGACCTACACCGAACTGAGATACCTACAGCGTGAGCTATGAGAAAGCGCCACGCTTCCCGAAGGGAGAAAGGCGGACAGGTATCCGGTAAGCGGCAGGGTCGGAACAGGAGAGCGCACGAGGGAGCTTCCAGGGGGAAACGCCTGGTATCTTTATAGTCCTGTCGGGTTTCGCCACCTCTGACTTGAGCGTCGATTTTTGTGATGCTCGTCAGGGGGGCGGAGCCTATGGAAAAACGCCAGCAACGCGGCCTTTTTACGGTTCCTGGCCTTTTGCTGGCCTTTTGCTCACATGTTCTTTCCTGCGTTATCCCCTGATTCTGTGGATAACCGTATTACCGCCTTTGAGTGAGCTGATACCGCTCGCCGCAGCCGAACGACCGAGCGCAGCGAGTCAGTGAGCGAGGAAGCGGAAGAGCGCCCAATACGCAAACCGCCTCTCCCCGCGCGTTGGCCGATTCATTAATGCAGCTGGCACGACAGGTTTCCCGACTGGAAAGCGGGCAGTGAGCGCAACGCAATTAATGTGAGTTAGCTCACTCATTAGGCACCCCAGGCTTTACACTTTATGCTTCCGGCTCGTATGTTGTGTGGAATTGTGAGCGGATAACAATTTCACACAGGAAACAGCTATGACCATGATTACGCCAAAGCTTGTCGCCGGAACGCAGCGACAGAGATTCCAATGTGTCCGTATCTTTCAGGCTTTTGCCCTTCAGTTCCAGACGAAGCGACTGGCGATTCGCGTGTGGGGTCTGCTTCAGGGTCTTGTGAATTAGGGCGCGCAGATCGCCGATGGGCGTGGCGCCGGAGGGCACCTTCACCTTGCCGTACGGCTTGCTGTTCTTCGCGTTCAAAATCTCCAGCTCCATTTTGCTTTCGGTGCGCTTGCAATCAGTACTGTCCAAAATCGAAAATCGCCGAACCGTAGTGTGACCGTGCGGGGCTCTGCGAAAATAAACTTTTTTAGGTATATGGCCACACACGGGGAAAGCACAGTGGATTATATGTTTTAATATTATAATATGCAGGTTTTCATTACTTATCCAGATGTAAGCCCACTTAAAGCGATTTAACAATTATTTGCCGAAAGAGTATAAACAAATTTCACTTAAAAATGGATTAAGAAAAGCTAGCTTGTGTAAGATTATGCGCAGCGTTGCCAGATAGCTCCATTTAAAACACTTCAAAAACAATAAGTTTTGAAAATATATACATAAATAGCAGTCGTTGCCGCAACGCTCAACACATCACACTTTTAAAACACCCTTTACCTACACAGAATTACTTTTTAAATTTCCAGTCAAGCTGCGAGTTTCAAAATTATAGCCGGTAGAGAAGACAGTGCTATTTCAAAAGCAAACTAACAAGGGTCTTAAATTCCAAAACACCAATCCTAACAAGCCTTGGACTTTTGTAAGTTTAGATCAAAGGTGGCATTGCATTCAATGTCATGGTAAGAAGTAGGTCGTCTAGGTAGAAATCCTCATTCAGCCGGTCAAGTCAGTACGAGAAAGGTCTCAATTTGAAATTGTCTTAAAAATATTTTATTGTTTTGTACTGTGGTGAGTTTAAACGAAAAACACAAAAAAAAAGTGATACACAGAAATCATAAAAAATTTTAATACAAGGTATTCGTACGTATCAAAAACATTTCGGCACAATTTTTTTTCTCTGTACTAAAGTGTTACGAACACTACGGTATTTTTTAGTGATTTTCAACGGACACCGAAGGTATATAAACAGCGTTCGCGAACGGTCGCCTTCAAAACCAATTGACATTTGCAGCAGCAAGTACAAGTAGAAAGTAAAGCGCAATCAGCGAAAAATTTATACTTAATTGTTGGTGATTAAAGTACAATTAAAAGAACATTCTCGAAAGTCACAAGAAACGTAAGTTTTTAACTCGCTGTTACCAATTAGTAATAAGAGCAACAAGACGTTGAGTAATTTCAAGAAAAACTGCATTTCAAGGTCTTTGTTCGGCCATTTTTTTTTATTCAACGCTCTACGTAATTACAAAATAAGAAATTGGCAGCCACGCATCTTGTTTTCCCAATGAATTGGCATCAAAACGCAAACAAATCTATAAATAAAACTTGCGTGTTGATTTTCGCCAAGATTTATTGGCAAATTGTGAAATTCGCAGTGACGCATTTGAAAATTCGAGAAATCACGAACGCACTCGATCGAGCATTTGTGTGCATGTTATTAGTTAGTTGTTTAGTTAATTGAAGTATTTTACCAACGAAATCCACTTATTTTTAGCTGAAATAGAGTAGGTTGCTTAAACAAAGCCACGTCTGAAAATTTCTTATTGCTTGTAGTTGTGACGTCACCATATACACACAAAATAATGTGTATGCATGCATTTCAGCTGTGTATATATACATGCACACACTCGCAACACGAAAACGATGACGAAGCAACGGAACAAAGGTTTCTCAACTACCCTTTGTTCCCTGTTTCTTCGCTTTCCTTTGTTCCAATATTCGTAGAGGGTTAATAGGGGTTTCTCAACAAAGTTGGCGTCGATAAATAAGTTTCCCATTTTTATTCCCCAGCCAGGAAGTTAGTTTCAATAGTTTTGTAATTTCAACGAAACTCATTTGATTTCGTACTAATTTTCCACATCTCTATTTTCTTCCCGCAGAATAATCCAAACTGCAGGTCGACTCTAGCTAGAGGAAGCTTATGGACTACAAAGACCATGACGGTGATTATAAAGATCATGATATCGATTACAAGGATGACGATGACAAGGATCCACTAGTCCAGTGTGGTGGAATTCTGCAGATATCCAGCACAGTGGCGGCCGCTCGAGTCTAGAGGGCCCGCGGTTCGAAGGTAAGCCTATCCCTAACCCTCTCCTCGGTCTCGATTCTACGCGTACCGGTCATCATCACCATCACCATTGAGTTTATCTGACTAAATCTTAGTTTGTATTGTCATGTTTTAATACAATATGTTATGTTTAAATATGTTTTTAATAAATTTTATAAAATAATTTCAACTTTTATTGTAACAACATTGTCCATTTACACACTCCTTTCAAGCGCGTGGGATCGATGCTCACTCAAAGGCGGTAATACGGTTATCCACAGAATCAGGGGATAACGCAGGAAAGAACATGTGAGCCATATGGGCCCATGTGAGCCATATGGTGCACTCTCAGTACAATCTGCTCTGATGCCGCATAGTTAAGCCAGCCCCGACACCCGCCAACACCCGCTGACGCGCCCTGACGGGCTTGTCTGCTCCCGGCATCCGCTTACAGACAAGCTGTGACCGTCTCCGGGAGCTGCATGTGTCAGAGGTTTTCACCGTCATCACCGAAACGCGCGA

To generate **pUb FLAG-Mts**, the following sequence was inserted between the SpeI and XbaI sites:

Cgaggataaagcaacaacaaaagatcttgatcaatggattgagcagttgaacgaatgcaatcagttgacagagacacaagttcgcaccctctgcgacaaggccaaggagattctctccaaggagtccaatgtgcaggaggtaaaatgcccggtgacagtgtgcggagatgtccacggtcagtttcacgacctcatggagctcttccggataggcggcaagtctccggacaccaactacctgttcatgggcgactacgtggaccgtggatactactccgtggagaccgtgacccttctggtggccctgaaggttcgctatcgcgagcgcatcaccatcctgcgcggtaaccacgagtcgcgccagatcacacaggtgtacggcttctacgacgagtgcctgcgcaagtatggcaatgccaacgtttggaagtacttcacggatctgttcgactacttgccactgacggcactcgtcgacgggcagatcttctgcctgcacggaggcctcagtccctcgatcgacagtctggatcacattcgggccctcgatcgcttgcaggaggttccgcacgagggtcccatgtgcgatctgctctggtccgatcccgatgacaggggtggctggggaatctcgcctcgtggcgccggttacacctttggccaggatatttcggaaacctttaacaacacaaacggcctgacactggtgtcgcgcgcccatcagctggtgatggagggctacaactggtgtcacgatcgcaatgtggtcacaatattctcagcgccaaactattgctaccgctgtggcaaccaagcggctcttatggaactggatgattcacttaaattttcattcctacaatttgatccagcacccaggcgcggggagcctcatgttacgcgaagaacacccgattatttcctttaa

To generate **pUb FLAG-Pp2A-29B**, the following sequence was inserted between the BamHI and XbaI sites:

Agcagcaagcgacaaatcggtcgacgattcactatatcccattgcggttctaatcgatgaactgaaaaacgaggacgttcagcttcggttgaactccatcaagaaactgtccaccattgcactcgctttgggcgaggagcgcacacggtccgagttgattcccttcctcaccgagaccatatacgatgaggacgaggtactgctggccctggccgaccaactgggcaactttactagtctcgttggtgggccagagtttgccatgtacttgattccgcccctcgagagtttggccaccgtagaggaaaccgtggtgcgagacaaggctgtggaatctctacgcaccgtggccgctgagcacagcgcccaggatttggagatccatgtggtgccgacactgcagcgattggtttccggtgactggttcacctcacgcacctctgcctgcggcctcttctcggtctgctatccacgcgtcacacagccagtgaaggccgagctgcgcgccaacttccgaaagctctgccaggatgagacacccatggtgcgccgtgcagcggccaacaagctgggcgagtttgccaaggtcgttgagacggagtatctgaagtccgatttgattcccaactttgtccagctggcacaggatgatcaggactctgtccgtctgctggctgtagaggcatgcgtaagcattgcccagctgctgcctcaggatgatgtagagcacctggttctgcccacgctgcgccagtgcgccagcgactcttcctggagggtgcgttacatggtggccgagaagtttgttgatctgcaaaaggctgtgggcccagagattactagggtggacttggtgcctgccttccagtacttgctcaaggatgccgaggccgaggttcgcgctgcagtggccaccaaggtgaaggacttctgcgccaatctggacaaggtcaaccaggtgcaaatcatccttagttccattttgccctatgtccgcgatcttgtctcggaccccaatcctcatgtgaagtcagctctggcctcagtgatcatgggcttgagtcccatgctgggcgcctatcagactgtggagcaattgctccccctgttccttattcaactcaaggatgagtgcccagaagtgcgcctaaacatcatctcaaacctggattgcgttaacgacgtcatcggtatccagcaactgtcacagtcgcttctgcccgccatcgtcgagctggccgaggactccaagtggcgtgtgcgtctagccatcatcgagtacatgcctgctctggccggtcagttgggtcaggaattctttgaccaaaaactgcgcggtctctgcatgggatggctcaacgatcacgtgtacgccattcgtgaggcagccaccctcaacatgaagaagctcgtcgagcagttcggagctccctgggccgaacaggccataattccaatgattctggttatgtcgcgcaacaagaactatttgcacagaatgacttgcttgttctgcctgaatgttttggcagaggtctgcggcacagatatcaccaccaagttgctgctgcccacagttctcctgcttgccgctgatcccgttgccaatgttcgtttcaacgtggcaaagaccctgcagaagatctcgcccttcctggaggccagcgtcattgatgcccaagtaaagcccacactcgacaaactgaacacagacacagatgtggatgtcaagcattttgctgcacaggccattgccggcatagctgcagcgtaa

To generate **pUb FLAG-Cka**, the following sequence was inserted between the SpeI and NotI sites:

Cggcaccaattcgggagccaccgctggcataaacaacaagccggttggcggtgcaacaggagccggcgtccttgtaggcggcggtgtgggcggtgccaattcctcgatcggcggtgtcctgtcgaacagcctgggcggtggcggcagcggcggtctgagcatcagcggcctcaacgctggtggacagaacgccaatgtgggcggaatgggcaacgttggcggcgacgacggcggaaacgggatggtgggcggcggtgtaaataaccagcaggccacaacgccccaatacacaataccgggcatcttgcacttcatccagcacgagtggtcgcgcttcgagctggagcgatcacagtgggacgtggacagggccgaattgcaggctcgcattgctatgctgcttggagagcgcaagtgcttggaaagcctcaaatcggatttgacgcgccgcatcaagatgctggagtacgcactgcgccaggagcgtgcgaaattctatcgtttaaaatacggcaccgatccgccgcagctcaatgagttcaagccatcgaacgaggatgccggactggccggtgaggtggccactgactcggaggtgccctacagcagtgtctcgaacactacgtggcggcaggggcgtcagatgctgcgacagtatctggccgagattggctacacagacaacatcatcgatgtgcgttcaaatcgggtgcgttcgatccttggcctaaacaataatgccgagcacgacggcagcggcggaggactgggtggtggacttggcggcggcaccggcggcgagaaccttagcccaaacatcaatggaaacgagagcaacaaacgcgcatcggagacggaaggccgtcatactcccgctaaaaaagtgcaacaatcgatcgatgaaataatcgtggacaccgaggcagctgtgatggcgaactttgagttcctgggcgccaccgagatgtccgacgacgacgagatatctgacgatctggaaatggtggccaccgacaatgacgacacagacgtaaagctggccaagcgcgccaagtctggcaaggatatgctcaccgaggaggtggacggctcgttgggactgggtgagcttgcccagctgacggtgaataacgaatctgatggcgcctacgatgcgaactccaaggatggaactggcggcagtgccggcggtgctggttaccgcaaaacctggaacgccaagtacacgcttcgctcgcactttgacggagtacgatcacttatctttcacccagaggagccggtgctgatcacggcttccgaggatcatacgcttaagctgtggaaccttcagaagacggtgcaggccaaaaagtcggccagcttggatgtggaaccgctgtatacgttcagggcccatacgggtccagttctgtgcctgggcatgtcgtccagcggcgagacatgctattcgggcgggctggacggtaacattgaatgctggcaattgccgtcgccgaatatcgatccgtacgattgctatgatccgaacgtgcattccggcacgctcgaggggcatacggatgccgtgtggggccttaccaccatgcaaagcaacattgtgtcatgctcagcggatggaacggttaagctgtggtcgccatacaataaggaaccgctgctgcgcacatacacagcctccgaggcagagggtgtaccctcatccgttgactttgttcgcaatgaggttgaccatatcgtggtggcctacaacagtgcccactgcatcgtgtacgatacggagacgggcaagcaggtggtgcgtctggaggcggcgcaggagatgtctggcaacacggggaagttcattaacaaggtggtgtcacacccaacgctacccattacgataaccgcgcacgaggatcgtcacatccgcttctgggacaacacatccggcacgctggtacactccatggtggcgcatttagagccggtcacctcgctggccgtcgatgctcatggcctgtatttgctatccggctcgcacgactgctcgatacggctgtggaacctggacaataagacgtgcgtgcaggagatcacagcgcatcgcaagaagtttgacgagagcatattcgatgtggccttccatgccacaaagccgtacattgctagtgccggggccgatggcctcgccaaggtctttgtctaa

To generate **pUb FLAG-tws**, the following sequence was inserted between the SpeI and NotI sites:

Cgccggtaatggagaggcgtcttggtgcttctcacagattaaaggcgccctagacgatgatgtcacggatgcagacatcatatcctgcgtggaattcaatcacgatggcgagctgctggccactggtgacaagggcggtcgcgtcgtcatctttcagcgtgatcctgcctcaaaagccgccaatccgagacgcggcgaatacaatgtctattcgacattccaatcgcacgagcccgaattcgactacctcaagtccctggagattgaggagaagataaacaagatccggtggctgcaacaaaagaatcccgtgcactttctgctctcgaccaacgacaagacagtcaaattgtggaaggtcagtgagcgtgacaagtcgttcggcggctacaacacgaaggaggagaacggactgatccgggatccacagaatgtaacggccctccgagtgccatccgtgaagcagataccactgctggtggaggcctccccacggcgcaccttcgccaatgcacacacctatcacatcaattcaataagcgttaattcggatcaggagaccttcttgtcggccgacgacctgcggatcaatctgtggcacttggaggtggtcaaccagagctacaacattgtcgacatcaagccaaccaacatggaggagctaacagaggtgatcaccgcggccgaattccatccgaccgagtgcaatgtgtttgtctactcgagctctaagggcacaatacgattatgtgatatgcgttcggcggcgctatgcgaccggcacagcaaacagttcgaggagcccgagaatccaacgaatcgcagctttttcagcgaaataatcagctccatcagcgatgtgaagctaagcaactcgggtcgctacatgatctccagggattacttgagcatcaaagtgtgggatctgcatatggaaacaaagcccattgagacctatccggttcatgaatacctgcgcgccaagctgtgctcgctgtacgagaatgactgcatcttcgacaagttcgagtgctgttggaacggcaaggacagctcaataatgaccggaagctacaacaacttcttccgcgtcttcgatcgcaactcgaaaaaggatgtgacgctagaggcgtccagggacatcatcaaaccgaaaacggtgcttaagccacggaaagtttgcactggcggcaaacgaaagaaggatgagatcagcgtggactgtctagatttcaacaagaagatcctgcacaccgcctggcaccccgaagaaaatatcatcgccgtggctgcgaccaataacctcttcatatttcaggataaattttag

To generate **pUb FLAG-wdb**, the following sequence was inserted between the SpeI and NotI sites:

ctcatcgggcacgtttgtggatcgaatcgacccgttcgccaaacgttcgctcaagaagaagggtaaaaagagtcaaggttcctcgcgctacagaaattctcaggatgttgagctgcagcaattgccgccgctgaaagctgattgctccagcctagaacaggaggagctgttcataaggaagctgcgccagtgttgcgtgtcctttgactttatggatcccgtgacggacttgaagggcaaggagatcaaacgggccgcgctcaatgacctatccacctatataacacatgggcgcggcgtgcttacggaaccggtttatccggaaataattcgtatgatttcttgcgacctatttcgcacgttgccgcccagtgagaaccctgatttcgatcccgaggaggacgatccgacactggaggcgtcctggccacacctccaactggtgtacgaggtcttcttgcggtttctggagtcgcaggatttccaagcaactatcggcaagcgtgtgatcgatcaaaagtttgtcttgcagcttttggaacttttcgactcggaggatcccagggagcgggactttctgaagacggtattgcatcgtatatatggaaaattcttaggactgcgggcttttatacgaaaacagattaacaacatattcttgcgattcatttacgaaacggagcactttaatggagtcggtgagctcttggaaatccttggcagtatcatcaacggattcgccttgcccctgaaggccgaacacaaacagtttctggttaaggtcctattgccactgcacaaagtcaaatgcctgtcgctctaccacgctcaactggcgtattgcattgtacaattcctagagaaggatccattccttaccgaaccggtggtgcgtggcctgctgaagttctggcccaagacgtgttcccagaaggaggtcatgttcctgggcgagattgaggagatcctcgatgtgatcgatccgccgcagtttgtcaagattcaggagccgctattccgtcagattgcaaaatgcgtgtcgagtccacattttcaggttgctgagcgagctctttatctgtggaacaatgagtacgccatgtcgctaatcgaggagaacaatgcggtcatcatgccgatcatgttcccggccctataccgcatcagcaaggaacactggaatcaaactatcgtggcgctcgtatacaacgttttgaagacgttcatggagatgaattcgaaactgttcgacgagcttacctccagctacaaggcggagcggcaaaaggaaaaaaaacgtgagcgtgaccgcgaggaactgtggaagaaactgcacgaactcgaatccaatcgttctagtgggcgcaccgccggcggcagcgccacgacatcgaacagcgcagcttccgcggcgtcaacgtccctgcagccacccagctccgctggcctgaactctcatcagcagcagtcgaatagcggcagcagcggtagcctgtccagcggcggtgccggcggcgacaataatcctgcgaccacaaatgcgaaaatcaaacaggataaggcggacaactaa

To generate **pUb FLAG-wrd**, the following sequence was inserted between the SpeI and NotI sites:

Cgataacgaggcgttagatccaacaataaaaagcagtacgtcggcagcgacgccaacagcagcagcatcagaaacgacaacaacagcagcatcttcagttgtggaaaccacaacaacaatcgcggcggccacagcctcagcggcagaatcaaaaaacgaaacaacggccacaaacaacaatagcaacacaagcggcagcatcagcagtagtagcagcaacaatatagtcataccggcatcggccactaacggtatcaaagagagtaacagtaacttaagtacaacaacaacagcagcagcagtagcggcagcaacaactgtagaaggagtagctccagcaataacgtccacaatcgtggtgaccggcggcactcctccattgagcagtttggctaacaaactcaaggacaacacaccaccgtacgatgcaccgccgcccacgccgatcagcaaggtcctgaacatcaccggcaccccgatcgttcgcaaggagaagcgccagaccagcgcccggtacaatgcctccaagaactgcgaactgacggccctcattccgctaaacgagaagaccgccgccagtgaacgtgaggagctgttcatacagaagatccagcaatgctgcacactgttcgacttctccgagccgctcagcgacctcaagttcaaggaggtgaagcgggcggctctgcacgaaatggtcgatttcctcaccaaccaaaatggcgtaataaccgaagttatttacccggaggcgatcaatatgtttgctgtcaaccttttccgaactctgccgccatcgtccaacccgaatggtgccgaattcgatccggaggaggatgagcccacgttggagtcctcgtggccgcatctgcaactcgtttacgagctgttcttgcgcttcttggagtcaccagattttcaaccaagcatggcaaaacgttttatcgaccatcaatttgtattacaactattggatttattcgattcggaggatccacgtgaacgtgatttcctgaagactgttttacatcgcatctatggaaaatttttgggcttgagagcatttattagaaagcagatcaacaatgtcttttacagatttatttatgaaacggagcatcataatggcatagccgaattgttggaaatcctgggtagcattatcaatggctttgctctgccgcttaaggaggagcataaacaatttttacttaaggtattgctgccattgcacaaagccaagagcctctcggtctaccatccgcagctcacctattgtgtggtgcagttcctggagaaggatcctagcttatcggaggcggtcatcaaaagcctacttaaattttggcccaagacgcacagtcccaaggaggttatgtttttgaacgagctggaggagctgttggacgtaattgagccggccgagttccagaaggtgatggtgccgctgttccgccaaatagccaagtgcgtctcttcgcctcatttccaggtggccgaacgtgcgttgtactattggaacaacgagtacattatgtcgctgataacggataactcggcggtgatattacccattatgttcccagcgctcaatcgcaactcaaagacgcactggaacaagaccatccatggtctgatctacaatgcactcaagctgttcatggagatagatcagcggctttttgacgagtgcagcaagaactacaagcaagagaagcagatggagcgtgagaagctgtcgcaaagggaggagctctggcaacaggtggagagcttggccaagaccaacccggagtggacaaaggcgcgccggtttaacgactgcctgccggtcagcgacagccgggccctgtgcgatcaatatagtgagaacagcgattcagcgtatgatcagagcgagcagagggcgcgccagccgccgcctccgctgccgccacagaaacaggcgcaccaggagccccgagaggtgagacaggcacttgccacattaacaacactaaacaactactaa

To generate **pUb FLAG-IntS6 AA 1-400**, the following sequence was inserted between the SpeI and XbaI sites:

cacaatcatactcttcctggtggacacctcgtcgtccatgtgccagaaggcgtatgtgaatggggtacagaaaacgtatctggacattgccaagggagccgtggagacgtttctcaagtatcgccagcgtacgcaggattgcctgggagatcgctacatgctgctcacattcgaggagccaccggcaaacgtgaaagctggatggaaggagaaccatgccaccttcatgaacgagctgaagaacctgcagagtcacggcctcacctcgatgggtgaatcgctgcgcaatgcgttcgatttgttaaatctgaatcgcatgcagtcgggcatcgatacgtacgggcagggcaggtgcccgttttatctggagccatcggtcatcattgtgattacggacggcggtcgctattcgtaccggaatggtgtccatcaggagatcatactgccgctaagtaaccaaataccgggcacaaagttcactaaggagccatttcgctgggatcagcgcttgttttcgcttgtcctccgcatgccgggcaacaagattgacgagcgagtggatggcaaggtgccgcatgacgattctcccatcgaacggatgtgcgaggttaccggcggacgttcatatcgagtgcgaagccactacgtactcaaccagtgcattgaaagtcttgtccaaaaggttcagcctggtgtggtgctgcagttcgagcctatgctgcccaaggaggctacttccgctaccgctggcgaagcggcgggtgcgagcacgatatccggcatgggaataccctcaacgtccggagcgggtcccgctcccgatattgtcttccatccggtaaagaagatgatctacgtgcagaagcacatcacgcagaaaacctttcccatcggctattggccactgccggagccctattggccggactccaaggcgattacattgccgcctcgcgacgctcatcccaagctgaaggtcctgacgccggcggtggacgagccacagctggtgcgcagcttccccgtcgacaagtacgagattgaaggctgtccgttaacgctgcagattctcaacaagcgcgagatgaacaagtgctggcaggtgattgtgaccaatggcatgcatggattcgagctgccctttggctacctaaaagcggcgcccaacttttcgcaggtacacctctatgtactggcctactaa

To generate **pUb FLAG-IntS6 AA 1-600**, the following sequence was inserted between the SpeI and XbaI sites:

cacaatcatactcttcctggtggacacctcgtcgtccatgtgccagaaggcgtatgtgaatggggtacagaaaacgtatctggacattgccaagggagccgtggagacgtttctcaagtatcgccagcgtacgcaggattgcctgggagatcgctacatgctgctcacattcgaggagccaccggcaaacgtgaaagctggatggaaggagaaccatgccaccttcatgaacgagctgaagaacctgcagagtcacggcctcacctcgatgggtgaatcgctgcgcaatgcgttcgatttgttaaatctgaatcgcatgcagtcgggcatcgatacgtacgggcagggcaggtgcccgttttatctggagccatcggtcatcattgtgattacggacggcggtcgctattcgtaccggaatggtgtccatcaggagatcatactgccgctaagtaaccaaataccgggcacaaagttcactaaggagccatttcgctgggatcagcgcttgttttcgcttgtcctccgcatgccgggcaacaagattgacgagcgagtggatggcaaggtgccgcatgacgattctcccatcgaacggatgtgcgaggttaccggcggacgttcatatcgagtgcgaagccactacgtactcaaccagtgcattgaaagtcttgtccaaaaggttcagcctggtgtggtgctgcagttcgagcctatgctgcccaaggaggctacttccgctaccgctggcgaagcggcgggtgcgagcacgatatccggcatgggaataccctcaacgtccggagcgggtcccgctcccgatattgtcttccatccggtaaagaagatgatctacgtgcagaagcacatcacgcagaaaacctttcccatcggctattggccactgccggagccctattggccggactccaaggcgattacattgccgcctcgcgacgctcatcccaagctgaaggtcctgacgccggcggtggacgagccacagctggtgcgcagcttccccgtcgacaagtacgagattgaaggctgtccgttaacgctgcagattctcaacaagcgcgagatgaacaagtgctggcaggtgattgtgaccaatggcatgcatggattcgagctgccctttggctacctaaaagcggcgcccaacttttcgcaggtacacctctatgtactggcctacaattatccagcgttgctgcccattctgcatgacctcattcacaagtacaatatgagtccgccgagcgatctgatgtacaaattcaacgcctatgtgcgttcaatacccccgtattactgcccgttcctgcgcaaggcgctggtcaacatcaatgtgccgtatcagctgctgcagtttctgctgccggagaatgtggacaactatctgtcgccaacgatcgccaatcagcttaagcacatcaagaatacggccaagcaggatcaggagaatctttgcatgaaggtgtacaagcagctgaagcagccgaagccaccatatcgccaagtggagacgggcaaactgttcacaggtgcgccgttacgcagagatttggtgcggcatccactgctccgggatatattttccaagttgcatggggaaatagatcccgtggagaattatacgattgtggtgccgcagccaacgcatcagtcgtcggccaagtcgcatcgcaatcccttcgatataccgcggcgggatttggtggaggaaatagccaagatgcgtgagaccttactgcgacccgtctcgctggtggccaaggactcgggccattgcctgtaa

To generate **pUb FLAG-IntS6 AA 101-1284**, the following sequence was inserted between the SpeI and XbaI sites:

cttaaatctgaatcgcatgcagtcgggcatcgatacgtacgggcagggcaggtgcccgttttatctggagccatcggtcatcattgtgattacggacggcggtcgctattcgtaccggaatggtgtccatcaggagatcatactgccgctaagtaaccaaataccgggcacaaagttcactaaggagccatttcgctgggatcagcgcttgttttcgcttgtcctccgcatgccgggcaacaagattgacgagcgagtggatggcaaggtgccgcatgacgattctcccatcgaacggatgtgcgaggttaccggcggacgttcatatcgagtgcgaagccactacgtactcaaccagtgcattgaaagtcttgtccaaaaggttcagcctggtgtggtgctgcagttcgagcctatgctgcccaaggaggctacttccgctaccgctggcgaagcggcgggtgcgagcacgatatccggcatgggaataccctcaacgtccggagcgggtcccgctcccgatattgtcttccatccggtaaagaagatgatctacgtgcagaagcacatcacgcagaaaacctttcccatcggctattggccactgccggagccctattggccggactccaaggcgattacattgccgcctcgcgacgctcatcccaagctgaaggtcctgacgccggcggtggacgagccacagctggtgcgcagcttccccgtcgacaagtacgagattgaaggctgtccgttaacgctgcagattctcaacaagcgcgagatgaacaagtgctggcaggtgattgtgaccaatggcatgcatggattcgagctgccctttggctacctaaaagcggcgcccaacttttcgcaggtacacctctatgtactggcctacaattatccagcgttgctgcccattctgcatgacctcattcacaagtacaatatgagtccgccgagcgatctgatgtacaaattcaacgcctatgtgcgttcaatacccccgtattactgcccgttcctgcgcaaggcgctggtcaacatcaatgtgccgtatcagctgctgcagtttctgctgccggagaatgtggacaactatctgtcgccaacgatcgccaatcagcttaagcacatcaagaatacggccaagcaggatcaggagaatctttgcatgaaggtgtacaagcagctgaagcagccgaagccaccatatcgccaagtggagacgggcaaactgttcacaggtgcgccgttacgcagagatttggtgcggcatccactgctccgggatatattttccaagttgcatggggaaatagatcccgtggagaattatacgattgtggtgccgcagccaacgcatcagtcgtcggccaagtcgcatcgcaatcccttcgatataccgcggcgggatttggtggaggaaatagccaagatgcgtgagaccttactgcgacccgtctcgctggtggccaaggactcgggccattgcctgcccatcgccgagatgggcaactaccaggagtacctcaagaacaaggacaatccgctgcgtgaaatcgagccgacgaatgtgcggcagcacatgttcggtaatccgtacaagaagaacaagcacatggtgatggtggacgaggcggatctgagcgatgtggcgccaatgaagtcgccaaatggcaatggaccgggtggagcgccacccggctcaccttccggatcgggctcgcccacaggaccgggtccctcatctccgggctcttcgcccggcggtggttcgggtcccgggatgcccggaatgcccggaatgggcggtggaatgtcaggactgatgctgggcgcaggtggcagtggcggatcctctaaaaaactggacggcacgtctcgtgggcgaaagaggaaagcaggcccgctgccacgtagctttgaattccgtcgatcatctacggactcgcgatcttcgagctccggctccgaattatccacaacgggtagtgcaccaggatcaccgataccaggcgccacgtcatccatatccggctgcgattccgccatgggcgctgatggcggtcccagttccagctgtttcgacgaggacagcaatagcaacagcagcttcgtgagctccacatcggaggcctcggccagtgattcagggatgtcaaactcgcatttggatcccttgcgaatcagtttcgttttcatcgaggaggagaccagcgatcaggatacgccgctcaatggttatgtgcacaatttcatcaatggcatcagcgacgatgcagaaacgggcgaaacggaagcagttccaggcggggcatccctgccgggtgcgtcttcagctaacgaaccgtcgtcaattggagcatcaccagcagtttcagccacgccagcaacatctgtgatccccgccagcaatggcagcgaagtttgctccgccgtcgcggccggcagcagcagttccatcagctccagcgccagttccccgggcgttctctacgtgcccctcaacggactgaccacccatgcggcctacagcagtaatctgggtggtccgagctcgccgttggacaaccttagctccctgggtggagcggcgggtggcggaggatcgggactactcggtgtcagcaagccgggatcgctgccactggccaatggttatggacacggtcatagttcagccatttccccaccttcgccattgtccttggtgcctctatcaccattgggaacaagtacgcccgctgccagcagctcgggactatcgtctagttccgcctacttcagccacaatgacatcaacgatgcctcggagatctcgcgcatactgcagacctgtcacaatggaaccagcagtggctcctcggtcggagcgggcgcaagcaacagcaatttgaatggcaatggcagcacggagagtggcggaggagtctcatccgacgatcatgcctcgctgggtggcaacagtttcgatgccctgaaacggacagcgggtactgcgggcagcactgcagcgggaaatccccactactactgggccagtggtctcaatcatcacaacaataacaacagcagcgccgctggagcagctagcggttcgacgttgagcaacaatcacaaccaccggggtcacaacaacagcagccccaacaaactggaggtgaagatcaactccagctgcggctcctcaccgacgcacaacaacgggggcatgggctcaccgggacagagcctgccgctcattttcaccgaggagcagcgcgaagcggcccggttgcacaacgtcgaactacgattgcagatcttccgtgatatacggcgacccggccgagactacagccagctgctggagcacctgaatctggtcaagggcgaccaggatatgcagtcggatttcgtggacatgtgtatcgtggaatcgctgcgcttccgccgccatcgcatggcgtccagcatacaggagtggtgggatcgcaagcagcagttaactgccgtgggaaccaatgaagcgacccccagcgcggagcaggccgtcgccaagagttaactcgag

To generate **pUb FLAG-IntS6 AA 1197-1284**, the following sequence was inserted between the SpeI and XbaI sites:

Caacgtcgaactacgattgcagatcttccgtgatatacggcgacccggccgagactacagccagctgctggagcacctgaatctggtcaagggcgaccaggatatgcagtcggatttcgtggacatgtgtatcgtggaatcgctgcgcttccgccgccatcgcatggcgtccagcatacaggagtggtgggatcgcaagcagcagttaactgccgtgggaaccaatgaagcgacccccagcgcggagcaggccgtcgccaagagttaa

To generate **pUb FLAG-IntS6 AA 101-1200**, the following sequence was inserted between the SpeI and XbaI sites:

tcttaaatctgaatcgcatgcagtcgggcatcgatacgtacgggcagggcaggtgcccgttttatctggagccatcggtcatcattgtgattacggacggcggtcgctattcgtaccggaatggtgtccatcaggagatcatactgccgctaagtaaccaaataccgggcacaaagttcactaaggagccatttcgctgggatcagcgcttgttttcgcttgtcctccgcatgccgggcaacaagattgacgagcgagtggatggcaaggtgccgcatgacgattctcccatcgaacggatgtgcgaggttaccggcggacgttcatatcgagtgcgaagccactacgtactcaaccagtgcattgaaagtcttgtccaaaaggttcagcctggtgtggtgctgcagttcgagcctatgctgcccaaggaggctacttccgctaccgctggcgaagcggcgggtgcgagcacgatatccggcatgggaataccctcaacgtccggagcgggtcccgctcccgatattgtcttccatccggtaaagaagatgatctacgtgcagaagcacatcacgcagaaaacctttcccatcggctattggccactgccggagccctattggccggactccaaggcgattacattgccgcctcgcgacgctcatcccaagctgaaggtcctgacgccggcggtggacgagccacagctggtgcgcagcttccccgtcgacaagtacgagattgaaggctgtccgttaacgctgcagattctcaacaagcgcgagatgaacaagtgctggcaggtgattgtgaccaatggcatgcatggattcgagctgccctttggctacctaaaagcggcgcccaacttttcgcaggtacacctctatgtactggcctacaattatccagcgttgctgcccattctgcatgacctcattcacaagtacaatatgagtccgccgagcgatctgatgtacaaattcaacgcctatgtgcgttcaatacccccgtattactgcccgttcctgcgcaaggcgctggtcaacatcaatgtgccgtatcagctgctgcagtttctgctgccggagaatgtggacaactatctgtcgccaacgatcgccaatcagcttaagcacatcaagaatacggccaagcaggatcaggagaatctttgcatgaaggtgtacaagcagctgaagcagccgaagccaccatatcgccaagtggagacgggcaaactgttcacaggtgcgccgttacgcagagatttggtgcggcatccactgctccgggatatattttccaagttgcatggggaaatagatcccgtggagaattatacgattgtggtgccgcagccaacgcatcagtcgtcggccaagtcgcatcgcaatcccttcgatataccgcggcgggatttggtggaggaaatagccaagatgcgtgagaccttactgcgacccgtctcgctggtggccaaggactcgggccattgcctgcccatcgccgagatgggcaactaccaggagtacctcaagaacaaggacaatccgctgcgtgaaatcgagccgacgaatgtgcggcagcacatgttcggtaatccgtacaagaagaacaagcacatggtgatggtggacgaggcggatctgagcgatgtggcgccaatgaagtcgccaaatggcaatggaccgggtggagcgccacccggctcaccttccggatcgggctcgcccacaggaccgggtccctcatctccgggctcttcgcccggcggtggttcgggtcccgggatgcccggaatgcccggaatgggcggtggaatgtcaggactgatgctgggcgcaggtggcagtggcggatcctctaaaaaactggacggcacgtctcgtgggcgaaagaggaaagcaggcccgctgccacgtagctttgaattccgtcgatcatctacggactcgcgatcttcgagctccggctccgaattatccacaacgggtagtgcaccaggatcaccgataccaggcgccacgtcatccatatccggctgcgattccgccatgggcgctgatggcggtcccagttccagctgtttcgacgaggacagcaatagcaacagcagcttcgtgagctccacatcggaggcctcggccagtgattcagggatgtcaaactcgcatttggatcccttgcgaatcagtttcgttttcatcgaggaggagaccagcgatcaggatacgccgctcaatggttatgtgcacaatttcatcaatggcatcagcgacgatgcagaaacgggcgaaacggaagcagttccaggcggggcatccctgccgggtgcgtcttcagctaacgaaccgtcgtcaattggagcatcaccagcagtttcagccacgccagcaacatctgtgatccccgccagcaatggcagcgaagtttgctccgccgtcgcggccggcagcagcagttccatcagctccagcgccagttccccgggcgttctctacgtgcccctcaacggactgaccacccatgcggcctacagcagtaatctgggtggtccgagctcgccgttggacaaccttagctccctgggtggagcggcgggtggcggaggatcgggactactcggtgtcagcaagccgggatcgctgccactggccaatggttatggacacggtcatagttcagccatttccccaccttcgccattgtccttggtgcctctatcaccattgggaacaagtacgcccgctgccagcagctcgggactatcgtctagttccgcctacttcagccacaatgacatcaacgatgcctcggagatctcgcgcatactgcagacctgtcacaatggaaccagcagtggctcctcggtcggagcgggcgcaagcaacagcaatttgaatggcaatggcagcacggagagtggcggaggagtctcatccgacgatcatgcctcgctgggtggcaacagtttcgatgccctgaaacggacagcgggtactgcgggcagcactgcagcgggaaatccccactactactgggccagtggtctcaatcatcacaacaataacaacagcagcgccgctggagcagctagcggttcgacgttgagcaacaatcacaaccaccggggtcacaacaacagcagccccaacaaactggaggtgaagatcaactccagctgcggctcctcaccgacgcacaacaacgggggcatgggctcaccgggacagagcctgccgctcattttcaccgaggagcagcgcgaagcggcccggttgcacaacgtcgaactacgataa

To generate **pUb FLAG-Human IntS6**, the following sequence was inserted between the SpeI and XbaI sites:

ccccatcttactgttcctgatagacacgtctgcctctatgaaccagcgcagccatctgggcaccacctacctggacacggccaaaggcgcggtagagaccttcatgaagctccgtgcccgggaccctgccagcagaggagacaggtatatgctggtcactttcgaagagccgccctatgctatcaaggctggatggaaagaaaaccatgcaacgtttatgaatgaattgaaaaaccttcaggctgaaggacttacgactcttggccaatccctaaggacagcttttgatttattaaatttaaatagattagtaactggcatagacaactatgggcagggaagaaacccttttttcttggagccagcaataattatcacaattactgatgggagcaagttgactaccaccagtggagtccaggatgagcttcatttacctcttaattctcctttgcctggaagtgaattgaccaaggaaccttttcgttgggatcagagactctttgcattagtgttgcggttgcctggcaccatgtcagtagaatcagaacagttgacaggtgtgcctttagatgactctgcaatcacaccaatgtgtgaagtgacaggcggccgttcatattctgtgtgttctccaagaatgcttaatcagtgtctggagtccttggtgcagaaagtacaaagtggggtggtaataaactttgaaaaagcaggaccagatccttcccctgtagaagatgggcagccagatatatcaaggccttttggatctcagccttggcatagctgtcacaaactcatatatgtcagaccaaatcctaaaactggggttcctataggtcattggcctgttccagagtctttttggccagatcaaaattcgccaacactaccacctcgtacatctcatcctgtagtgaagttttcctgtacagactgtgaaccaatggttattgataaacttccttttgacaaatatgagttggaaccttcaccactgactcaatttatcctggaaaggaaatctcctcaaacatgttggcaggtgtacgtgagcaatagtgcaaaatacagtgaacttggtcatccttttggttacttgaaagccagtacagcactgaactgtgtcaacttatttgtgatgccttacaattatccagtccttcttcccctcttagatgacttgtttaaagtgcataaagcaaaaccaacattgaagtggagacagtcatttgaaagttatttgaagacaatgcctccctactatcttgggcccttgaagaaagctgttaggatgatgggagcacctaacctaatagcagacagtatggaatatggacttagttacagtgtcatttcatacctcaaaaaactgagtcaacaggccaaaatagaatctgatcgagtcattggatctgtaggcaaaaaagtagtacaggagactggaataaaagtccggagccgatcacatggtttatcaatggcatataggaaagattttcaacaactcctccagggaatttcagaggatgtccctcacagactgctagaccttaatatgaaggaatacactgggttccaagttgctttgctgaataaggatttgaagccacagacatttagaaatgcttatgacataccaagacgaaatcttttggatcacttaacaagaatgagatctaatcttttgaagagcactcgcagatttctgaaaggacaggacgaagatcaagtgcacagtgttcctatagcacaaatggggaactaccaggaatacctcaagcaagtaccttctccactaagagaacttgatcctgatcagccacgaaggttgcatacatttggcaacccctttaagctggataagaagggtatgatgatagatgaagcagatgaatttgtggctggacctcaaaataaacataaacgacccggagaaccaaatatgcaagggatccctaaaagacgtcggtgtatgtctccactactaagaggcagacagcagaatcctgttgtaaacaatcatattgggggaaaaggaccacctgcacctacaactcaagcacagccagatcttattaaacctcttcctcttcataaaatttcagaaaccactaatgattcgataatacatgatgtggttgaaaatcatgttgcagaccaactttcatcagacattacaccaaatgctatggatacggaattttcagcatcttctccagccagtttactggaacggccaaccaatcatatggaggctcttggtcatgaccatttaggaaccaatgacctcactgttggtggatttttagaaaatcatgaggagccaagagataaagaacaatgtgctgaagagaacataccagcatcttcactcaacaaaggaaagaaattgatgcattgcagaagccatgaagaggtcaatactgaactaaaagcacaaataatgaaagagatccgaaagccaggaagaaaatatgaaagaatcttcactttactgaagcatgtgcaaggcagtttacaaacaagactaatatttttacaaaatgtcattaaagaagcatcaaggtttaaaaaacgaatgctaatagaacaactggagaacttcttggatgaaattcatcgaagagccaatcagatcaaccatattaatagcaattaa

To generate **pUb FLAG-Human IntS6L**, the following sequence was inserted between the SpeI and XbaI sites:

ccccatcctgctgttcctcatagacacgtccgcctctatgaaccagcgcactgacctgggcacctcttatttggacattgccaaaggcgctgtggagttattcttgaagctgcgcgcccgggacccggccagccgtggagacaggtacatgctggtcacctacgacgaacccccgtactgcatcaaggctggttggaaggaaaatcatgcaacattcatgagcgaactaaaaaatcttcaggcttctggactgactactctcggtcaggctctaagatcctcatttgatttgttaaatctcaatagattaatatctggaatagacaattatggacaggggagaaatccattttttttagaaccatctattttaattaccatcacagatggaaacaagttaacaagtactgctggtgttcaagaagagctccatcttcctttgaattcccctctgcctggaagtgaactaaccaaagaaccttttcgttgggatcaaaggttatttgccctggtgttgcgtttgcctggagtggcttctaccgaaccagagcaactagggagcgtaccaactgatgaatctgccatcacacagatgtgtgaagtcacaggaggtcgctcctactgtgtgagaacacaaagaatgttgaatcaatgtttagaatctctagttcaaaaagttcagagtggtgtagttattaattttgaaaaaacaggaccagatccacttcctattggagaagatggacttatggattcatccaggccaagcaattcatttgctgctcagccatggcatagttgtcataaactcatttatgtacgacctaactctaaaactggtgttcctgttggacattggccaattccagaatctttttggccagatcagaatttaccttcactacctccacgaacatctcatcctgttgtgaggttctcctgtgtagattgtgagccaatggtaatagacaaacttccttttgacaaatatgaacttgaaccttcgcccttaactcagtatatcttggaacgaaagtctccccatacctgctggcaggtatttgttactagcagtggaaagtacaatgaacttggatatccatttggttatttaaaagccagtacaactttaacttgtgtaaacctctttgtgatgccttacaactacccagttttacttcctcttttagatgacttgtttaaagttcacaagcttaagccaaatctgaagtggcgacaggcttttgacagctacttaaaaactctgcctccatactacctattaaccaaactagagtcagaacgaatactagcatcagtggggaagaaacctccccaggaaattggaattaaagtgaaaaatcattctggaggtggcatgtccttgactcacaataaaaattttagaaaactattgaaagaaatcacaggggaaactgcacttagactgacagaattgaacaccaaagaatttgctggcttccaaattgggctcttaaacaaggatttgaaacctcagacatacagaaatgcttatgatattccccgtagaggtcttttagaccagctgaccagaatgagatccaatctgctgaaaacgcacaagtttattgttggacaagatgaagattcccttcatagtgttccagttgcacaaatgggtaactatcaggaatatctgaagacattggcttctccactgcgagagattgatccagaccaacccaaaagactgcatacttttggcaatccgtttaaacaagataagaagggaatgatgattgatgaagcagatgagtttgtagcagggccacaaaacaaagtgaaacgtccaggggaacccaacagtcctatgtcatctaagagaaggcggagtatgtccctgctgttgaggaaaccacaaacaccacctactgtaactaaccatgtgggcggaaagggaccaccctcagcctcgtggttcccatcttatccaaacctcataaaacccacccttgtacatacagatgctactatcattcacgatggccatgaggagaagatggaaaatggtcagatcacacctgatggcttcctgtcaaaatctgctccatcagagcttataaatatgacaggagatcttatgccacccaaccaagtggattctctgtctgacgacttcacaagtctcagcaaagatgggctgattcaaaaacctggtagtaacgcatttgtaggaggagccaaaaactgcagtctctccgtagatgaccaaaaagacccagtagcatctactttgggagctatgccaaatacattacaaatcactcctgctatggcacaaggaatcaatgctgatataaaacatcaattaatgaaggaagttcgaaagtttggtcgaaaatatgaaagaattttcattttgcttgaagaagtgcaaggacctctggagatgaagaaacagtttgttgaatttaccatcaaggaagccgcaaggtttaaaagacgagtcctaattcagtaccttgagaaggtactagaaaaaataaattcccaccaccttcacaacaacattagtcacatcaacagcagatcatcatgttaa

To generate **pUb FLAG-Zebrafish IntS6**, the following sequence was inserted between the SpeI and NotI sites:

ccccgtgctactttttctaatagacacgtccgcctccatgaaccagcgctcccatctgggcactagctatctggacatagcgaagggcgcagttgagactttcctgaagctgagaagcagagatccggccagcaggggagacagatacatgttagtgagtttcgaggaagcgccggctggcattaaggctggatggaaggacagtcatgccacttttatgactgagctaaggaacctgcaagcagttggactgacatcgtttggccaagctttaaggacagcttttgatttgctcaatcttaatagattagtttcagggatagacaactatggacagggtagaaacccttttttcttggaacctgctatcattgtggccataactgatggcagcaagctgactggcagctctggggtacaagatgagttacatttaccgttaacaacaccattacctggcagtgaactgaccaaagagccctttcgatgggaccagcgactcttctctttagtgctgcgtgttccaggccatgcctctgctgaccctgaaccagtgggtggagttccactcgactcctcccctattacacccatgtgtgaggtcaccggaggccgatcatacagcgtgttttcccagagaatgctgaaccagtgcctggagtctttggtacagaagatacagagtggtgtggtcatcaattttgagaagactggacctgatccttctcctactgaagatggtccaatagaggtgaagcatggacctcaagtatggcacagctgtcataaattgatctatgtgagacccaaccctaaaactggtgtacctgttggccattggcctatccctgaatccttctggcctgaccagaactctcctactttgcctccacgtgcagctcaccctcatgtgaagttttcatgtgtggattcggagcccatggtggtagataaagtgcccttcgacaagtatgagctggagccttccccgctcacccagtacattctagagaggaaatcaccacatacctgctggcaggtttttgtgtgtaacagtgctaaatatggagaactgggtcaaccgtttggatatctgaaggctagcacagctctcaactgtgtcaacctgtttgttatgccctataactatcctgtggtgcttccattgcttgatgacttaattgttgtgcataagttcaagcctccagtgaagtggcgacagtcttttgaaaattatcttaaaacaatgcccccctactacatcccagccctgcggaaagcattaagaatgatgggggctccaaacctgttagcagacaacatggaatatggcctcagctacagtgttgtgtcttatcttaaaaagctcagtcagcaggcgaagatcgaggcagaccgagtttgtgcttcagtgggaaaaaaagtggcaccagaaggtggaattaaacttcgttgtaggagctctgctctatccctggcaaacaggaaggatttcacacagcttctgcagagcatcatgggagatggaccagcacttccaatggaggcaaacaccaaggagtttgctggctttcagctggcttccttgaataaagcattaaaaccacagggcttgcgcaacccttacgatatcccaagagctcatcttctggaccagctgagccgcatgaggaggaaccttctgcatgccagtatctgtattctcaaagggcaagatcaagatcagcttcacagtgtgcctatagctcagatgggcaactatcaggacttcttgaagcattgcccatcacctctgagagaggcagaccctgatcagcccaaacgtctgcacacctttggtaacccctttaaactggacaagaaggccatgatggtggatgaggcagatgagtttgttactggcactcagggcaaaggcaaacgtcctggtgagtccagcagtcctactgtgggaggggcacctaaacgcaggcgctgcatgtctccattgcttcggcctggacgggcatatacacctcctagaacaccatccagaactcccgataacaatcacttggaattggaaaaccatatcaccaatcatctcgagccaaactgcgactcggacagcgatataaatcctgtgccggagtctgaagtgatccagcaaagcaaccatctccagacagaagaggttgagaaccagcaggactgcgtgcaggagaatggccagtccagtgacagtgagctgtctttgggaagcgagggggaagaggaggccccgcaccgctaccagtacccctgccagctgaagaagatcagaaaccaggagagccaggagctcaactctgaactccgagtgctcatcaccaaggagatcaggaaacctggaagacgttatgagaaaatcttctaccttctcaagcaaatccagggcagtttggaaacgcacctgatcttcttgcagagcataatcaaagaggctgcaagatttaagaagagggtcttaatcgagcagctggagattttcatagaggagattcacttaagagccaacggcatgaatcatctggacagtttctga

To generate **pUb FLAG-Zebrafish IntS6L**, the following sequence was inserted between the SpeI and NotI sites:

ccctattttacttttcctgatagacacgtccgcttctatgaatcagcgcacttatttgggtacgacgtatctggacattgctaaaggcgcggttgagatctttatgaagctgcgtgcccgagacccggctagtagaggcgacaggtacatgctagttacattcgatgatccaccatacggcgtaaaggcgggctggaaagagaatcatgccacattcatgagtgagctgaagaacctgcaggcatctggactgactacactgggtcacgctcttcgtgccgccttcgacctgctcaatctcaacagacttgtctctggcatcgacaactatgggcagggtcgaaatccgttctttctggagccgtcagtcatcattacaattacagatggaaacaaactgacacacagctcgggagtggctgaggagcttcacctgccgctgaattccccattgcctggtagtgagctaactaaggagccgttccgatgggaccagagactcttcgccctggtcctgcgtctgccaggagtggccgtgcccgacagtgagcagctgggaagtgtgcccactgatgagtctgccataacccagatgtgtgaggtcactggaggccgctcttactgtgtaagaacccaaaggatgctgaaccagtgtcttgagtctctcgtccaaaaggtgttgagtggagttgtaatacattttgagaaaaccggtccagatccaccagtgattggtgaagatggtcttgtggatccagctcgtcctttgacgtccttcagtccacagccttggcacagctgccacaagctcatttacgtccggcccaaccccaagactggcgtacctgtgggtcactggccaatctcagaatccttctggccagaccaaaactcccctactttgcctcctcgttctgcccatcctgtggtgcggttttcctgtgttgactgtgagccgatggttatcgacaagctgccttttgacaaatatgagctggagccgtccccgctcacccagtacatcttagaaaggaaatctccgcacatgtgctggcaggtgtttgtgaactgcagtggcaaacatagtgatgttgcccacccgttcggctacctgaaggccagcaccacactcacctgcgtcaacctcttcgtcatgccttacaactatcctgtcctgttgccactcctcgatgatttgtttaaagtgcacaaactaaagccaaacctcaagtggcggcagtcctttgaaatgtacctgaaatccatgcctccgtactatctactgcctctcaagaaggcgctaaggatgatgggagccccgaatctcatctctgataacatggattgtggcttgagctacagtgtcatttcctacctgaagaaactcagccagcaggcaaagattgagtcagaccgtctgattgtgtctgtcggtaagaagcctcctccagagacgggcatcaaggtgaagaaccactcgaatgccttgtctctggcacaccgccgggatttcaaacagctcctgcagggcatcacgggtgaggttcccctccggctcatcgacatgaacttcaaagagtttgctggtttccagatcgcacttctcaacaaggatttgaagccccaggcctataggaatgcttatgatattccacgacgaaaccttctagatcaagtgactcgaatgcgttccaaccttttgaggacaacgcaaaagctcatcagaggccaggatgatgactctttgcacagtattccagtgggacagatgggaaattatcaggagtatctgaagatgatgccatcacctctgcgtgagattgatccagaccagcccaaacgactgcacacattcggcaacccattcaaacaagacaagaagggaatgatgatcgatgaagccgatgaatttgtgacaggtcctcaaaataagaagcggggtaacacgggggatctaaactcgggtacagctctaaagaggaggaggagtatgtcccctctgctccgccggccccaaacccctccaataataaccaatcatgtattgggcaaagggcccacagggactcaaggccaacaaggaatcatcaagcccattccactacacaaaggagctgaggggaacaatgtaggaggcacagaaagcaatggcgagcgagtggcaggtactgacgctggagactgctggcctggagaagtggacggagaatcaggagaacccgcacctgtggaggacagagaagacgctgcggcaccggacggagaggaggagatactagcactggaaaattgtcttgatgacagatcccctgaccacacgcaaaactgtgaggagctcagtcctccgggccaggagggggagatggaagtcaatgaaggagacactccagcacagggcacgattgtcatgattcccctggaggggagtaacgcagagctgcgcactcgggttatcaaagaggtccgcaagcctggccgcaattatgaggcgatatttaggctgctggaggaggtgaaagggcctgtatcagtccagaggtactttattcatcatgccatcaaagaggcagccaggtttaaaaagcgtatgctgattcagcagctggaaaccgctctggaggagattgaagacagacagatgctacccgcacaaatcaacaatgtccacagcagatagt

The following *Drosophila* expression plasmids were obtained from Eric Wagner’s group (University of Rochester) and were generated from the previously published **pUb 3xFLAG MCS** plasmid (Chen et al. (2012) *RNA*, 18, 2148-2156):

**pUb FLAG-IntS1**

**pUb FLAG-IntS2**

**pUb FLAG-IntS3**

**pUb FLAG-IntS4**

**pUb FLAG-IntS5**

**pUb FLAG-IntS6**

**pUb FLAG-IntS7**

**pUb FLAG-IntS8**

**pUb FLAG-IntS9**

**pUb FLAG-IntS10**

**pUb FLAG-IntS11**

**pUb FLAG-IntS12**

**pUb FLAG-IntS13**

**pUb FLAG-IntS14**

To generate **pUb FLAG-IntS1**, the following sequence was inserted between the SpeI and XbaI sites:

cgatcgcgggaaaggaagcggctccaaccgatcgcagaagaaggttccgctgggcggagaactctttgcgctcggcaagagcgtacgcgatgactccaagtcgaagatcctgcccattaagggcatgtcctcgtcggaccgcaagcgtgaagcctccaccgcactggccagctcgagcaagcggttccgcggcaacctgaaggacgcgggcgccccggacatgtcctccggcagtagtcagtgcgagacgtgggagcaattcgccgtcgactgcgatctggataccgtggtcgagaccatatacgcggctttggagcagaacgacagcgaaactgtgggtcggcttgtatgcggcgttataaagcagacaacgacgagctcctcgcgctccaaggtggacaacattgcccttctggccctgatctacgtggccaaggtgcagccgactatattttgcacggacatagtggcctgcgcgctgctgtccttcctgcggcgggaggccaacgtgaaaatgaggtacaacaccaacctgcacattctgtttgccaatctgctgacccggggcttcatggagatttcccagtggccggaggtgctgctccgcatttacatagacgatgcggttaacgagcgctactgggcggacaatgaactgtgtgccccgctcgtgaagaacatatgcgcagcctttaaaacccgcacgccacacatcagtctcctccgctgggatgtgagctcgtccctgccttctggtcaagcacacagagacagcatgaccgtggacgacgattccggggataattccacacagagcttggatgcaagtcccttaaataccgaatcggaaccaattcccgacgccatgtgcaccactaaatccagattcagcgatgctgtggtgcagaagcatgtcagcgatgccattcgcgatcagctcaacaagcgacagcagcaggataactacaccagaaacttcctaaagttcctgtgcaccacttctggcattgcagaggtacgatgtctgagcatttcccgcctggagctgtggatccacaacggaaaactggttaagtttgcccagcaattgctctcgtacatctgcttcaacatcaagggacgcaacacacaggacaacgaagtgctcttggtgctggtgaagatgcggctcaagacgaagcccctaataaaccactacatgtcctgcttgaaggagatgatatttctgcagccggaaatcttgagtaccgtaatgaaactggtcgtgcaaaatgagctatccaacaccaggaatccaaacaatatggggatgctggccacaatgtttcaaacatctgctgatcaatcagctgccactctggctgagatttaccaggaattcttgcttcagcgggacgattgcctgcgcaccctgcgcgttttcttacgggagcttgtgcggatgctgcgcttcgatgtgaacctggtcaagttttgcaagacgtttctcagtgagcgggaggatctgactcctcaaatcgaaatgttcgagtttaaggagcgaattttcaactccatggtggacattgtgtgtctctgcatgtttctgtcggccacgccacaagctagggaagccagtttgtccctgaaaaccaatcgggacaccaaaaacaaccatgctctgctcaagctgtacaatcagatgtcacagattcagttggacacagtgtcctggatgtacgaaacagtacccacgctcttcaagattccggctgcggagtaccaccaggcgctgcacaagctgctactgctggacagtccggagcagtattcgcgctgcgatcagtggccgtcagagccggagcgaggagcaatcctgcgtattatctctgagacaccgattcacgaggagaccctgctgcgaattatattgattggcatcaccaaggatattcctttctccatcgcaaacacctttgatgtcctgctgctggtcataaagcgagtttccgggatgaaggctaccaatattccggcggtgcaggccaacaaattcgacatcatcgacttcctgttcagcatgtcagagtaccatcatccggagaacattcgcttgcctgcggagtacgagccaccaaagctggccatcatcgccttctactggaaggcgtggctcatattgcttatgatatccgcacacaacccctcctcgtttggagccttctgctgggaccactatcccacaatgaagatgatgatggagatttgtataacgaatcagttcaacaattcatcggccaccaaggatgagttgcagataataaccatggaacgggatcacattctgcagttcgagacgtacttggctgcccaaacatcgccgcatgctgttatcactgaggaaactgcgattctaatcactcagctcatgctgatggacccgatgggcactccgcgcaaagtgccgtccatggtcctggatcagctaaagttccttaatcagacctacaagcttggccacttattctgccgctgccgcaagccggacttgctgctggacataattcagaggcagggcaccacgcaatccatgccctggctttccgacttggttcagaacagcgagggcgatttcagccatctcccagtgcaatgcctgtgcgagttcctgctcttcaacgcgcacatcatcaacgaggagaatagccgcgatgccgagttggtcaatttcctgagaaacctgatatttgacggcaatcttagccaccaaatcgtttgcgagcttttggactacatatttagacgcttgtcgtccacggtgaagcaatcgcgagtcgccgctctttcgggtctgaagataatcttcaggcactctggcgactttgaaaacgaatggctcctgaagtcgctccagcaaattccgcacttctacgaggtgaaacccttcataattccccagctgcgagctgcctgccaggtggagaattgcccggaactgattatggcgtacattcagttcatcactgcgcacacgctcaacgatccggtcaacgaaatgctggatcatgtcatcgatatggcgcagctgattgtagagcgcagcacaatgtttcagcacattatcatttcgcaggaggactacgactatgtgcccgacgagaaccgcattcagactctgaagtgtcttttcgtcatgttcaacaactatataattaagctaagggaataccacgaaccgtacgagtggaccgagtatccggacctgctgatggttcagtttgacgatggagtacagctaccgctccacataaacatcatccacgccttcattatcctgctcacctattccaacagcaacatgcccgaatcgatacccattttggactattggtttccgcccggacgaccagcgcccgtcgcctttctgcccagcatgtcgcaggagcaggtgcaactgctgcccgactggctgaagctgaagatgatccgctcatcggtggacagactaattgaggcagctctcaacgatctaacgccggatcagatcgtgctctttgtgcaaaactttggcacgccggtcaactcgatgtccaagctgctggcaatgctggacaccgctgtactggagcagtttgatctggtgaagaatgccattctgaacaaggcctatctggcccaactaatcgagattcagcaggcgcggggtgctaagaatgggcactacaccgtacaggccttggatctgcattcccactcgcagactgtgccagatctgcccaagatcagtgtcgttattcaggaggccgttgaaattgatgattacgattcttcagactcggatgatagacccactaacttcctggccaccaaggaggtagcccaaaccattctcacgcagcccgaccagttgactgagtcgcgaagtgactgccgatccttgattcaaaaactgttggacatgctagcaagcccgaacagcaatagagcggatgtggttaacgccataacagaggtgctagccgtgggttgcagtgtcacgatgagccgtcacgcctgcacatttttaaggactttcttcagctgcatgctgcacagcgacaagtatcacatactggagaacgctctccaaaagaacttaagcatgtttaagcacacattcgccgactccagtctgctgcagaaatccgaactctatcatgagagcttggtgttcatgctgagaaactctcgcgagatttatgcgcagcagtttaaggcgaataccgccttggtggcacgaaagcggatcgtacgggcgatcgtccaaagcttcgatcagaccaaggacagcaagaccgtcgccaagtccaagagcgaccagctcttccacaacgggctcttcatcgactggctgtccgaaatggatcccgagatagtttccactcagctaatgaaggagcgctttctgttctcaaagtcctgcagcgagtttaggttctatctgttgtccctgatcaaccaccaaaccaactgggacacgatcgaaaggattgccgagtacctgttcaagaatttccatgaagactacgactacgccaccgtcctcaactacttcgaggcactgaccaccaatccgaagctgtggaagggacgcgagaagtacatgtcgaagaacgtacggccggatgccttcttcatgctgaggacttctgaactggagccgttctcccacttcatccttcacgaagggctttccgaggtcaagctggacagcaaaaactatgatttcaagctctgctcgcgaatgaacctactgtttaagctcacagagaagcgacgagacctcatggtcaaggtaatggagcacgtggagaaaagttcggtgtccgactacttgaaactgcaggtcctccagcagatgtacatcatgtatccgcgcatcaagtttctgaaaccgggcaaaaccggcgagcaggcgtacaaattgcagaatctgaagggctgccaggcggacaaggtgtccaacaatctgatcacctgcttgggcagtctggtgggcaaaaaggactttgagaccctgtccacggacaccgagctgctccttcgcaagctggctgcctcccacccgcttctcttccttcgccagctgggcgtgctgtcgtccattatgcaaggtcgggcccagctcagcatgaaggccctgcgcgaggagcaccacttccaccgatttgtacagatcctgaggacgctggagctgctgcaaccaaccatctttgaggaggcgtacaagaacgagatccaaaacacgctgtcgtgctacttcaacttcttcaagcaccacagcaacgtaaaggaggcctgccaaatgctgaacaagtttgtgcagatgctccaggcctacatcaactacaatccctcaagtgctctgctcttcatcgaacagtacgtgggcatcctcaaggaacttgcagccaagtacacctcactaggcaaactgcaggttttggtccaggccgttgccctgctgcagcacaagtcgcactcggcgacggaattggacgacgaggaagttaagtacgagtacgatctggatgagcatttcgatgtaaagccatcggccagcaagcccgtggtaacagaggatcccatcgaagtgaatccgcaaacacccatcgatccaagcagcagtaggggtcccttatcggttctaaccttgggctcgtacagccggtcgaactacacggacatatcgccgcacttcctcgatctggtcaagatcataaagcagtccaacacggaggacgtggtcttgggtcccatgcaggagctggagtgcctcacttccaagagatttgtgttcatcaacgagctgttcgaacgactgctcaaccttatattctcgccgagtgcccagatccggtccatcgccttcatcatactgatcaggcatctgaagcacaatcccggcaactcggacatcaacctgtgcacccttaacgcctacattcagtgcctgcgggacgagaactcctcggtggcagcgacggccatcgacaatctgccggagatgtcggtgctgctgcaggaacacgcaatcgacatcctaacggtggccttctcgctgggcttgaagtcgtgcctgaacactggccaccagattagaaaggttctccagactctagtgatccagcatggctattaa

To generate **pUb FLAG-IntS2**, the following sequence was inserted between the BamHI and XbaI sites:

aatgccggtgaggatgtacgatgtatcgccgcgcgttttctgcgccatgcagaatctggacatcaccctgctggccagctatccggaggcggagattcggcccgtgctgccctcgctggtgcggatgagcctgctatcgccactggacaacaccgaatcgtcgatggagtcgcgcaaagagatcctggccgtgctcatcggcatcgaggtggtgaacagcatcgtctcctacctgcaagtcaactaccatgagctggagaatgagctgaagaaggagctgcaggcccgccagaagtcggccttcttcgagggacagcaacacgagtacggcctccagtcgggcatcgcactgggctttgagcgggcggatgtggcgcgcaaggtgcgcgtggtgctctccgagatcttcaatctgcagcagcaggtctccgagcagaaaccggccgcccattcggagatgctggacgacggcatctatctggaggaggtggtcgacattctgtgcattgctctcgccgagctgccctccctgctgaacatactggagctaacggacgcccttgtccatgtgcccaacggacatcgcatcatctgcgcactggtcgccaactttccggattgctatcgcgacgttgtgtcccatgtgatcgccaattgcgacgaggatggcagcgatggcaagcaccggttaatgctgctcatggggctcagcgagatgaatccctcccaggcgctggccaacaggtccatgtgcgtggatatgctgaaggtgccgtcgtttatgcttaagctcacgctcaagcatcccgaggatctgatagccttcctcacgggcctgctgctgggcaatgaccagaatctgcgctcctggttcgccgcttacatccgctccagccagaagcggaagggggacgcactgaatctggtgcgcgtggagctgctacagaaggtgattcaaacgacgaccaatgcggcggagctgcgcgacttcaatctgcagggcgcggtgctgctgcgtctgtactgtgccctgcgcggcatcggtggcctgaagttcaacgacgacgagatcaatgctctgtcgcagctggtcacgagttgtccgcaggcgacgccctcgggcgtgagatttgtaacgctggcgctgtgcatgctcatcgcctgtccctcgctggtgtccaccattccactggagaacaaggctgtggagtggctgcagtggctcatacgggaggatgcctttttctgcaagcgccccggtactagcacctcgctgggtgagatgctcctgcttctggccattcactttcacagcaatcagatctcggccatcagcgagatggtctgctccacgctggccatgaagatacccatccggccgaacagtacaaacagaatcaagcagctcttcacacaggacctgttcacggagcaggtggtggcgctgcatgcggttcgcgtacccgtcactccgaatttaaatggcaccatactgtgctacctgccggtgcactgcatccagcaactgctgaaatcgaggacatttctcaagcacaaagtgcccattaagtcgtggatcttcaagcaaatatgcagttcggtgaggccagtgcatcctgtgatgccggccctggtggaggtctttgtcaacacgctgataatacccaatcccaccggaaaagtgaatattgatcacatgcaccggcccttcactgaggcggaaattctgcacgtgttccgcacatccaagctcacattcttcgccgaggagctgccaccgatggcggaaagtcaggagctcaaccagatcgaggtgacgtgtccactaaccgcccagctgctgatgatctactacctcatgttgtacgaggatacgcgactgatgaacctcagcgctttgggtggtcgcaagcagaaggagtactccaacaacttcctgggcggactgccactgaagtatctgctccagaaggcgcaccactaccacaacgactatctctcgctcttccatccgctgctgcgtttgattatctccaactatccgcatctcagcatggtcgatgactggctggaggagcacaatctggcccagggcaattccacggtggtggtaagcaagcatgagctcaagccggagaccctggatcgtgcactggccgccatccaaacgaaaccgcatctagccatccgcgtgtttaagcagctactccagatgcccccagaaacgcaggcgcagtatggacaacagctggtgaagcatctgccgatggtgtttgccaagtcggtgccgcgatacgtcaaggatctgtacaacgatatttggctgcgcctgaatgccgtgctgcccaccaccctgtggatcatgtcgctgcgagcgattaccaacggctcggatacgatggatcgacgtacctttgccaacgagagtcttctggagcccatggaggtgctaagttgtccacgcttcgtattctgttcgccctatttgctgatgatcctgctgcgcattctgaagggcagcctagccgcctcgaagacctatctcaacgtgcacatgcagatgcagcagaagcaggtgctggacaagaatggcatgatgcagacggatgcgatttgggaggacctaaggaccacgctgattgcctcgcaggagagcgccgccgtgcacattctgctggaggtgctggactacatagccagcaaggcaacggatcgggtgtcgcatctggagctgcgtgagatacagggcatcattggcacctatgtgcatcaggcattcatttcggagccttcgctggcgaaactagtgcacttccaaacgtatccaaagtcggttattcccatgatggtggccagtgtgccatcgatgcacatatgcattgattttgttcacgaattcctcaacgtgacggaaatggaaaaacagatctttaccatcgatctgacctcccatctggtgctcaactattcgatacccaagagtctgggcgtctcgaaattctgcctgaatgtaatccagacgacgttgtccatgctgaccgcatcgaccaagtgtagattcctgcgcaatgtgatgccagcgatggttcgtttcgtggaaacgtttcccattctggccgacgactgcgtcaacatcctgatgaccaccgggcgcattctgcattcgcagtcatcgctgggcatgaccacaatggaaatgccactcaccgatagcgacaagctctgcacgtatcgggatgcccagctgcacatcatcatgatcgaggacgcgtttaaggcactggtcacggcggtcatgaaaaaatcggacctgtattag

To generate **pUb FLAG-IntS3**, the following sequence was inserted between the BamHI and NotI sites:

agaacagcagcaatcaaaaaataatgctcacgtctcaaagttgttcatttgtacggctgttgattgcaaagatgatattgaggagaaattcgagcgttcgtttgtcactctgcagatgcaaatttctggccttagtgacaaagagatgcatgatatgttgtcacaagctgtgtgtaaagaaaaacagcatgaagaaatatcaatcggctttctctacataatgcttaccgacccttctatggcttccaaaacatatcgggacgttactctcgtatctagagacggaatgaatggcattgtcacaaatctcacgtttctggtcgctgagaaatatacaaagctcacggaggtagcgagaaggcaattaatttggttgttacgggaattcgtaaagcaccaggttctaagtgttgaaaatgttatatggaactgtttgcgtcaggctggcggaggagatgtttcctccagaaacctttttcttatcgagagccttttggacatattcatcgaatttagaacgtggttggaaaccaacccatttctcgttcagtctacagtttacagcttcgtacgcctaatagaagatcacgctaaccctgctctcttgtcgcttcggcagaaagaagtgaagtttacgatatcattgatccgagaaaggttccatgatattatacctctaggaagagattttgtacgcctcctacaaaacgtagctcgaattcctgagtttgaacaactgtggcgggatattcttttcaatcctaaaatgttgcatcaaacttttaacggaatttggcaactactgcatatacggacaagtcgaaggtttctgcaatgccgtctgctgcctgagatggaacgaaaaataagcttcttagcatcttctgttaagtttggaaatcagaagagataccaagattggtttcaggacaaatactttgctacacctgagtctcatagcttgcgatctgatctaattcgatttatcataaatgtcatccatcccacaaacgatatgttgtgctctgatattataccgcgatgggcaatcataggttggctcatctcgtcctgcaccaatccaatcgcttctgctaatgcaaagttgtcattgttttatgattggttattttttgatcctgccaaggataatataatgaatatagagcctggaattttagttatgtaccactctataagaaatcatccatttgttagtagcacgctgttggatttcttatgccgtattacaaaaaactttttcgtaaaacatgaagataaaattcgcattggtgtttataattcattgaaattaattcttgaaaagcaggtgataccaaatctgcaaccactttttgagtcacccaaacttgacagagaactgcgtaatttaataagggacaattttagagagtttttatcaccgccggctaatcacggacagctactgttcacttccctgcatccagttcagggtcatattttaaaaaaagagagcgatcaaagaattttacattgtgaaaacatggaccttcatgaaactgggctcataaatattagtgggacaatcgatgaagacaaaaagatcttactcgtgccaacggatcaagaaattgaaagtgtttttagcgacgaaactgctgaaaatttaagaagagttcacaacattgaggacaatacggatgacgacgatgacctaccattatcagaggtaagacttaaagaaaagccaaaggtagaactagcagaagcaatagccgaatcttttgatgcatttgttacgaagcgtaactcatacacttgggaggcgtttttgaaagatttccggccattaccagcgtctgcgtttgaagagtttcagctgaactatgtgatatcaaatactgttttaattttacgggagactttgcctcaacaaaatatattttccgagagcaaaaccgaagaaaaacatttggcaaaaagtataagttatcctttgtacggtttgtttcgatttttatatgaaaacgacgaaaaaagcaaaaaaccatttcaaaccttgctgtcagaaatctgtgaaggaatacctgaaattggctatttgcttctgtactttatgaaaatttattgtaagcttcaaacgcgcaaaaactcccaacaatcatatcagtttaaaacaactatttatcgacaaatttgcgatgcggcagatgagaaaatcggcaattgtctgttacgggatcttgatctgctagaaaaggaaaacacaaatatttttttatggctcctgcctgatatatatagagagtttaagtcaatagcaacaaataacactgatcttttgcgaataacactgcgttgcatagatgcaaaaaatgtgcgtgacatattatactcaatcgcacaaggaaaacttacaatatttaaacaagatggtctcattgattgcataaggcaaagtctcgagtttgagacatacgaacagttttgcctttggcaaattgttcaagctcacgatgtacccttaaggtgtatacaggacatattgccagaactggaagccggaagtcatcccgaagctttaagccattttctgttattgcttaaaaatgaagaacccacaaatgaaattattcgattaatgcttagcagagaatcgaagtcgaaaggtgatccattcgtaacctcagccttaagattctggtgtcaacgctatgaagaaaaactgtctgaaatcattgcttccctgcttacatcaaaataccccagttcatctcccaacaaaagaaaacgaccccccaaggggatttctgtctcgactagcactccttctgctgatcaagtattaaatcatctggagcattatagaagaagttgcagacatggcactggaactggcctttacgttcatgatatgatgcagagagcacttcaatcggcttactctcacagtaatgatagtactaagaaacaattttgtgatttgttcgccctggcagcggaagaagataccacagttggccgacgtgggggaagcggacgtggacgtaaacaaccagggagtaagaaagatgtcaataatcatgggacaagtaaaaaaatcccgagatggttaaaacaatatattcctccgatgacaattctagcgaggaggattggtcaaaatctaaaatattgcaaacggctaagagaaggaaaaaagctaacaatgattctgactga

To generate **pUb FLAG-IntS4**, the following sequence was inserted between the SpeI and NotI sites:

cgcgctggccattaagaagcgcgtaggcacctacgtggaaacggtggacgggtcgccgccggtgaagaagctgcgtctgcaaacgctggccgctgatgccaagggcgggaaatccggaaaagtgggcaatgtggagcgcaagctgacggcgctcaaccaattggacgcctatgtgggcaacctgcccgccggtgcgttggtcctgcccactgggacaccagttgcttcgacgggagccccgtcaacaggtgtcatcggaaatccgccggctgcggcgactggagctcccccaatgacggccgcgaatagcagggaactgctggagctcttggtcaagataaccgatgagatatcctatgaggacgtggaaatgggcgagcttaaggaggtggccagcaagatcttccaactgtatcagctgcaggaacgcgatagcgacacctccatccgggtgaagctgctcgagctgctgtccggattgggttgcgagtgtgccaccgagcaggcgcttaccatgatcatcgactactttatcttcttgttgaggaaggaggtgtcccaaaaggtgctcgcccagggcatgatgtgcctgtttcgaatcggggagcgcaggaagcacatgttgcccatatcgtacaaaacccaggtggcccacctagccaaggagcagctgcgctccggttcggcgcatacgcagaagaacgccatgctggtaatcggtcgttttgccaccaaaatggagggtgagcgtcattacgtatggaagttagccttctacatcgactcgcaggactcgagcgttcgtgcccaggctctgcacgccctgctgaccctcggcgaacggggatcccaactgccggcggtgctctataagcgtgccgtcgaggccatgaaggatgactacgagtgtgtgcgcaaggaagctctacagctggtcttcatgctgggcaatcggcatccggactacattttaccctcggaccgccagcaggaggagctgcgcatgatagacgctgcgttcagcaaggtctgcgaagccctgtgcgatctatcgctgcaaattcgcgtgctcgcagcggaattgctcggtgggatgaccgccgtaagcagggagttcctacaccagacactcgacaaaaagctgatgagcaatctgcgccggaagcgaacggcgcacgagagaggtgctcgcctggtggccagcggtgagtggtcatcgggtaaacggtgggcggacgatgcgccgcaagagcacctcgatgctcagagcatctcaattatagccagcggcgcctgtggcgcccttatacatggcctagaggatgagttcctcgaggtgcgtactgcggcagtggcctccatgtgcaagctggccctctctcgaccggatttcgcggtgaccagcctagatttccttgtggacatgttcaacgatgagatcgaggatgtgcggctgaaggccatctatagtttaacggccattgccaagcacatagtgctgcgcgaggatcagctggagataatgcttggatcgcttgaggattactccgtggacgtgcgtgagggactacatctcatgctgggcgcctgtagagtgtccacacaaacgtgcctgctaatggtggtgcagaagctgctagatgtgctggccaagtatccacaggatcgtaactccacctatgcctgcatgcgcaagattggccagaagcatccacatttggtgatggctgtcgccgttcacctgctttacgtgcatccctttttcgagacgcccgaacgcgatgtcgaggatccagcgtatctgtgcgttctgatacttgtcttcaatgctgcagaacatttggtgcccattatcagtttgctgcccacggccacccatcgccattacgcctatctgagagactcgatgccgaatctggtgccccagttgcccattgagggtgcgtcctcggcgagtgcgacgcacaggatcgattcggctatgcaccaagcgggcagttcggcggagtacctgcagatgattctcagtcacatcgaagagatcttcaccatgactgacgagcgcttggagctgctccagactgcccaatctaatctgcagcgcctgggatccattgatgcgggcatgtacggcacctcaaactttctagagaccttcttggcggcccagatccagatcgaacagatgcagcgttgcgccagcacgcaacgaagcagggtgcccttgaaggaatcactggcagcactcatccgaaactgtctcaagctgcagcacactttctccggcctcaattacggcgatatcttgcaggtgaagcagctgcgcctgagggcttgcgcgcttcacttggtcctcgtggtcagggatcgctcacagagtgcattgggcccgtgccagatgcttttgcaaaccgccggcgatataagtgagttcatcaaggcgaatacgaaggatgaggaggagaagccaccggtggtagagaccgatatgcccatgaaagagtcggtcagtagggatgcccagccggatagcttcacccgccagctgctgatcaaactggatggcatatcggatcccaagccgggtcgcgtctttcgcgaaattctgccgctggtccagcaggctcctccgcttgcattgccacccgccaatgacaaaatacgccgctgtgtggccaacatccttgagccgtgtccgctgcaatcgcaggataatgtaattaaggtcacggctggcctgatagcagccgttccttttgttgcggaaatcgacaatctgctggagtcccagaaggcggacatgcggatcaagatcaagtatccggaccagcacatgcacaccgtggtgccgaagcagagtgacttcaagcccattatgacggagcagggcgagcacaagaccaatgtgcgcctgcgcaccacaatcctgctgtcgcacagcgtttggacggagtcgtcgctggtggagatccaactgtgcctggccgtgcgtcccggcagcgaactggagctctgcaagccggcaaaggtgctgttcgcacccaaaccagtcaggcggggcatttag

To generate **pUb FLAG-IntS5**, the following sequence was inserted between the BamHI and XbaI sites:

actgcgccagaacctgttggatcagcttaagcacttcatagagacggtgagcaatgggcacagttgtccgcagctgctgaccagccccaatctcatcaagctcgcactgggattcctggaggagctgcccgccacgcgagacattgtgttcgaatacttcgccctgctggccgaaatcagtgtgcagctgtatgtgtcgccggagatggccgacccaaagaccggtatgccggtatcccaggtgaagctggccgggaatcgccagcagcagcaacgtgctccggaatacgaggctttcaatctggtaaagaccgcgctgcagagtctggtgtggaagggaccgcccgcctggtcgccgctcatcgccaactggagtctggaactggtggccaagctgtccgataagtacacccagcggcggatgaccatcacggccagctgcaactattggctggagtgcagcgccatgcacgggctgatgacccttataaatagttgtttccggaagttaacccaaccggaggaggaggcctgcgtggagatcatgctcaatgcctttcaccggtttcccatgaccttcgactggattgtggcgagattgggcggctgtttcccttacaagatcatcatgcagattctacagtgcggcatcaagcgattcgtcgatgactaccgctgccacctggactcggaggcgggcatcctcgattatatgacctcgtgccacgaacagcacctgagggccgcctttcgtgagatgcttagggagggattcgcaccgaagaagccgctggacgtggccgtcgttcctttcctgctgatcaccaccaactactcggacacgatcctgcagagcttggtcaatgtgctagtcgaaatctataccgaggacatgtgcgaggtgatagtgcagaaggcgccgttgtggctgagcaacaaaatgtttgccgatatgcagcccacgctgaataatgccgtgctgcgcctgaacgagcgtggagcgacgttgctccttacggcagccaaaatggccgagaagtatgtttggtgccaggatttcctggacaattcgatgcaggagctggaacagtgggtgctcaaccagcggaacttcccgttgctggccgatctggcctatgaggaaaccaagtacatgctgtggaagagctgcctcagcaccaatctcttcgagcagcaaacagcggtgagattactgctggtggtctcatcccagcacccgaatatttactaccaaactatatcccaacttctaaagaagtcgtatgctcaaaatcccaatggcattggcgcattaattcgcttacttggcggacaaagcggcatggttaactttccgggattcacgccaggtttcaagatggtcttggaggacatcacgttggatgttcaagttaacaatcgattgccagttccgccgggcacacccacagaagcctttaacactttctccaacctaaacatacttgccaggatgcacaagagcaagaatgtggctccgtacatcaaagcgcagcatttgaaccaagccctcaacgaatgtcttcccaacatccttcaaatcttcgactgcaccgttaacaaactggtcctgaggatagacagggatgccgccgagcggatagcggataaattccgggcgcagcaaagtaaaaatagcaataacaacaatgaattgtgcaatggcaaggattacggaaagcgcacaaaattggagccgggagaagataaggtggatgatgaggacgctacccgcatgcgcttggcccatctcattgtggatctgttgaacaacattgaagccggcagccggacaactgtcctgcgcactccattagtcctcaagctggccacgctaagcgtaaagtacttctttgtgggcttaaccgagaagactgtgatccgtcgagcagcggcttcgcatcgctcctacacgctgttgcagaggcagtgttccgcccggaaaatcgcgagaaccgtctgcctgcgcgagctggtggagcgcgcgctcttttatcacggtcacttgcttggtcagctggaggtgtatcagctcgatgagcttgagatacccgagcacgaacacctcatcctgcagaatctgcacaccagctctggcgccaactcgaatcgctccgtgctgcattccggtattataggcagaggtctgcgaccagtgctgccccccagcgagcggaactgcgatgcggagaagcaggcgctgtatcttaaggcattgaatgcatgctgcgccgatctggagaagcccaacaacgtggagggctactcgctggtctcgctgctgctggtcgagctggtctccacggatgtcatgtacaacggcctgccctttccggatgaggagttcaccagggtcacaatggagcgggacatgctaatcaggagagcgttcatcaactccccagtgctgtgggctgttttgggactgatcgccggtcataggccggcgctttgctattcctccgtgctactgcgcgcactgtgcgccacctgcttgcatcactggcggggcaagaatgtcaatcgattccagcccactgctgcgaacgatgagctgatgctgtgcaccaagaagatgctgcaactgctggctatgagccaactaattccacctccactcaccaatctgcatctcatcatcgagcactttgaatcggcagagattgccctgctgctgcgcgagtgcatctggaactatcttaaagatcacgtgccctcgccggcgctgtttcacgtggacaacaacgggctgcactggcgcaacacgaatacacagctggccaaggtgccgccgcaatatgtggacccgttgcgccacctaatgcagcgtaagctgtccaccctgggcccccactaccatcagatgttcatcatgggcgagctgatggagggcgactcagagccagatccgacggcccggctgcagatcgttgaaatagattaa

To generate **pUb FLAG-IntS6**, the following sequence was inserted between the SpeI and XhoI sites:

cacaatcatactcttcctggtggacacctcgtcgtccatgtgccagaaggcgtatgtgaatggggtacagaaaacgtatctggacattgccaagggagccgtggagacgtttctcaagtatcgccagcgtacgcaggattgcctgggagatcgctacatgctgctcacattcgaggagccaccggcaaacgtgaaagctggatggaaggagaaccatgccaccttcatgaacgagctgaagaacctgcagagtcacggcctcacctcgatgggtgaatcgctgcgcaatgcgttcgatttgttaaatctgaatcgcatgcagtcgggcatcgatacgtacgggcagggcaggtgcccgttttatctggagccatcggtcatcattgtgattacggacggcggtcgctattcgtaccggaatggtgtccatcaggagatcatactgccgctaagtaaccaaataccgggcacaaagttcactaaggagccatttcgctgggatcagcgcttgttttcgcttgtcctccgcatgccgggcaacaagattgacgagcgagtggatggcaaggtgccgcatgacgattctcccatcgaacggatgtgcgaggttaccggcggacgttcatatcgagtgcgaagccactacgtactcaaccagtgcattgaaagtcttgtccaaaaggttcagcctggtgtggtgctgcagttcgagcctatgctgcccaaggaggctacttccgctaccgctggcgaagcggcgggtgcgagcacgatatccggcatgggaataccctcaacgtccggagcgggtcccgctcccgatattgtcttccatccggtaaagaagatgatctacgtgcagaagcacatcacgcagaaaacctttcccatcggctattggccactgccggagccctattggccggactccaaggcgattacattgccgcctcgcgacgctcatcccaagctgaaggtcctgacgccggcggtggacgagccacagctggtgcgcagcttccccgtcgacaagtacgagattgaaggctgtccgttaacgctgcagattctcaacaagcgcgagatgaacaagtgctggcaggtgattgtgaccaatggcatgcatggattcgagctgccctttggctacctaaaagcggcgcccaacttttcgcaggtacacctctatgtactggcctacaattatccagcgttgctgcccattctgcatgacctcattcacaagtacaatatgagtccgccgagcgatctgatgtacaaattcaacgcctatgtgcgttcaatacccccgtattactgcccgttcctgcgcaaggcgctggtcaacatcaatgtgccgtatcagctgctgcagtttctgctgccggagaatgtggacaactatctgtcgccaacgatcgccaatcagcttaagcacatcaagaatacggccaagcaggatcaggagaatctttgcatgaaggtgtacaagcagctgaagcagccgaagccaccatatcgccaagtggagacgggcaaactgttcacaggtgcgccgttacgcagagatttggtgcggcatccactgctccgggatatattttccaagttgcatggggaaatagatcccgtggagaattatacgattgtggtgccgcagccaacgcatcagtcgtcggccaagtcgcatcgcaatcccttcgatataccgcggcgggatttggtggaggaaatagccaagatgcgtgagaccttactgcgacccgtctcgctggtggccaaggactcgggccattgcctgcccatcgccgagatgggcaactaccaggagtacctcaagaacaaggacaatccgctgcgtgaaatcgagccgacgaatgtgcggcagcacatgttcggtaatccgtacaagaagaacaagcacatggtgatggtggacgaggcggatctgagcgatgtggcgccaatgaagtcgccaaatggcaatggaccgggtggagcgccacccggctcaccttccggatcgggctcgcccacaggaccgggtccctcatctccgggctcttcgcccggcggtggttcgggtcccgggatgcccggaatgcccggaatgggcggtggaatgtcaggactgatgctgggcgcaggtggcagtggcggatcctctaaaaaactggacggcacgtctcgtgggcgaaagaggaaagcaggcccgctgccacgtagctttgaattccgtcgatcatctacggactcgcgatcttcgagctccggctccgaattatccacaacgggtagtgcaccaggatcaccgataccaggcgccacgtcatccatatccggctgcgattccgccatgggcgctgatggcggtcccagttccagctgtttcgacgaggacagcaatagcaacagcagcttcgtgagctccacatcggaggcctcggccagtgattcagggatgtcaaactcgcatttggatcccttgcgaatcagtttcgttttcatcgaggaggagaccagcgatcaggatacgccgctcaatggttatgtgcacaatttcatcaatggcatcagcgacgatgcagaaacgggcgaaacggaagcagttccaggcggggcatccctgccgggtgcgtcttcagctaacgaaccgtcgtcaattggagcatcaccagcagtttcagccacgccagcaacatctgtgatccccgccagcaatggcagcgaagtttgctccgccgtcgcggccggcagcagcagttccatcagctccagcgccagttccccgggcgttctctacgtgcccctcaacggactgaccacccatgcggcctacagcagtaatctgggtggtccgagctcgccgttggacaaccttagctccctgggtggagcggcgggtggcggaggatcgggactactcggtgtcagcaagccgggatcgctgccactggccaatggttatggacacggtcatagttcagccatttccccaccttcgccattgtccttggtgcctctatcaccattgggaacaagtacgcccgctgccagcagctcgggactatcgtctagttccgcctacttcagccacaatgacatcaacgatgcctcggagatctcgcgcatactgcagacctgtcacaatggaaccagcagtggctcctcggtcggagcgggcgcaagcaacagcaatttgaatggcaatggcagcacggagagtggcggaggagtctcatccgacgatcatgcctcgctgggtggcaacagtttcgatgccctgaaacggacagcgggtactgcgggcagcactgcagcgggaaatccccactactactgggccagtggtctcaatcatcacaacaataacaacagcagcgccgctggagcagctagcggttcgacgttgagcaacaatcacaaccaccggggtcacaacaacagcagccccaacaaactggaggtgaagatcaactccagctgcggctcctcaccgacgcacaacaacgggggcatgggctcaccgggacagagcctgccgctcattttcaccgaggagcagcgcgaagcggcccggttgcacaacgtcgaactacgattgcagatcttccgtgatatacggcgacccggccgagactacagccagctgctggagcacctgaatctggtcaagggcgaccaggatatgcagtcggatttcgtggacatgtgtatcgtggaatcgctgcgcttccgccgccatcgcatggcgtccagcatacaggagtggtgggatcgcaagcagcagttaactgccgtgggaaccaatgaagcgacccccagcgcggagcaggccgtcgccaagagttaa

To generate **pUb FLAG-IntS7**, the following sequence was inserted between the SpeI and XhoI sites:

catgtctcacctgaccggcacccgcgtgagcaccttcaacgagagcttccttaacgaaaatgagcacgactcgaatgcggtgctcatggaactggacaagggactgcggagcaccaagcagggaatccagtgcgaggcggtcgtccgctttccgcgcctcttcgagaagtatccgtttcccattctcatcaactcgtcgttcattaagctggccgattactttgtcagcggatcgaatctgctgcgcttctgggtgctccgtgtttgccagcagagcgagaatcatctggacaagatcctcaacattgacagctttgtgcgctgtatctttgtggtgatgcactccaacgatccggtggcgcgtgctctcctcctgcgcaccttgggcgccgtttcccgtgtgattcccgagaagcaacaggttcatcatgccattcgacgtgctctggatagccacgataccgtcgaggtggaggctgccatctatgccagcagctgctttgccgcccagtcaagttcctttgccatcagcatgtgcgccaagatttcggacatgatcgaatcgcttcaggtgccggtgccaatgaagctgctcctgattcccgtgttgcgacacatgcatcacgaggctaccacagcgtcactggttagcagactgtgcatggatctgctgcccaaatatccagcccagagctttgtggtggccatcatagacacgctcactcaattgtcatcacgaacgctggtgggagtgcccggtcagctggatgttctgctggatttcatgcaggacttaaggacaccagtacgcatacaggtgctgcgatctttcaacgagttggccggacgtcagagcgttcatgcctggcccaagccagccatcaaagcgctcattgatcgttttgaattgtgtaccaattccaaggagcagttcctcttcctctcaatcctgctgaagttaagcgagtgcccgcttacttgccagcagctgcttcgcgagcatcgcgtagccctgctccgtttgtgcatccagtgcattagcaagctggacgactacaccaccgccactcaggccatggcagttttgtcggttctagtagcatttggtttaaaaaaaaAgggcagcggcgagcaggtggacgatattcttcatatggtcaacctgcacatggaaggattgctcctttgcacggccaagcggtcggagtgcacccgcgatctccgtagggtgctgacatacggtataaggatcactaaagccaacgctgaatttggaacatccttcattgggatcgtgaccaatagcctgggagataagggtgcctatccgccagccaatgcggaacttatgtgcgaggcactggcgggtctgtgtgagcactttcagctgcgtaagtacgccttttccaccgcagaagatttgatagtagacgaaaacgcgatggacaccgatgagttgcccccgcccgagataaatcccatgctagctcgtttgccgctcattctgcacaagctgaacaccattatcgaccaggagaattgcgatcaacagttgcgcagcgtggagatcctgagttctctggttctacaaaccaccatgggctgttacctgccgcaaaaggttgttcagtgctttgagaaatgcctcggtaggcttaactgctggacgctgtataggattgcacgcactgctagtcgctatggtcaccactatgtggctgctcacatttacaccaaggtctcccagattgtgatcagcgatcatatgcactatttcctggtggcactgtcgcagatctcgcaggcggagtgcattcttaactatggcttggaatacgcctacatgcgggacaattatgcaccaaaggtggcaccggaacctctaattccgctgatgaagcgcttggaaatggctagtaatctctatcagcaagcgctggccagcctgcgagctggctcctcgccccagcatccgtgcacctttcagttggagtatctgaagataagggcgcagtttctgcagacccttcacctggctgtgaccgtaaagaatgcccaggtgattgtgccgccaccggcaatagctggaagtttggcccagaactcgcgcgactatctgcaaaagtttggtcatgtcacgaaccagctgcgcaagctggtgaaggcactaaaggcctgcgaggagacttacgctaggctatacaaatcctctttcgatgcggatcacgtgacgctggagtttctggaggtggctgaatttcagtgcgctcttttcgcccacatcatcgagtcgatttgctacgccacgcctccggaaccgcctgtatttctaaccacaggcgatcacccggagacgcgctactttgccgcaagctgccagcgcatggagcagatgcaaaaaaacctgccacaggagccggccaatgccaagaccattagtaaccgacatttggatgtcatcattgcgcagatcgagatcataacaaagacaccgctctgcctgccgcgttacttcttccagattctgcagtccacacagatcaagctatcggtgagtccgcagccaaggagcgccaccgaacccgtgaatgtccagtcgggcagtaatctggttatcaaggtcgagggtgtcctgcagcactttagcaagcagaagaaacacttccgtcgcgtggagtccgttcaattaagcctcacctcacagctaatcacgccgccaccgcgatcctcgcaagagctgcccaagcagggcgccaacgatactgtgaccctcaatcagattgtaaagccccagcgcgatttcctctccggcagcttcctgttgcccatctccaatggcgggcacttccaggtgacgctggaaacattcgtggtggacgagaatggcattacctggtgcactggccccaaatcctcgatggtggtgcgggtgctggaggatcctagcaagcagggcgctccagcgccgagcacttcccaggcagtgggacagacgaggaggttttaa

To generate **pUb FLAG-IntS8**, the following sequence was inserted between the BamHI and BstBI sites:

agacgatcctcttaagcccaagccagttccactggctgccgaaacggtactgtggttcgagttcctgctcgatccgcacaagattacgcagcatctgcagcgcccgcatcccgagcccagcgccatggagctgatcgtgcagttcatcagcatgacgccgaacacggcgcaggagtcagtggggacgcctggcagtgatttgcagaatctaaaccagacgccatcgaattcgggacccattcccggcgtggttggtggtgcccctgcgccgacaacgcccactgcctccggcggagtgggcatgccccatagcccacaaaggcccgcggagaagggcctgcaattgaaccgcaagcagttggcactgaaaatcctcgagctgaaggtagccacttggctgaagtgggatctggatgcattggagaagaacctgcctgtgattatgcagctggccttgctgcgcgatctgtgcaccataagctacggctgctccttaagtattcccctgcccaatgattttgacgcgagaatttccgctgctggaaacgaaagagcagcaagattcgccttgaccatataccaccgcatgctgctgcgaatgcaactgataaaggagcaggcgttaaaagcgccacgtcctcaaaacacaatgtaccaaactgtggaccagctccagcagtttctggacacaccgactcaaccatccatcgagtatttgcaacaactctgcgcctcgacaaaacctttctacatatttcactacgacagctttgttccgctgcgatgtgatgatatcggcaatggccaaaactacgatgttatgcacctaataacaccccaggaattgagagcccaactgcactacgagctcgcccaatattacctgtacaccaaacagtatgttttggcccgggaagcggccgccgcgtgtaatactaatctgcaggcaatcccgccacagacaactctatattactgccacatacgaccgtcagaactggaaggtctactgcaggcctgtggaattagcgccgaggagcagtcactgctggagaagttccagcaatcgctacttaacaactacacagacattgtatcaattctgcgaatggacaacagaaggcgagaaatcccctttataagccggcggcaagtggaactggacattgagggttccatctccacgggcatactcaaggagacagttcagctgcaattgcaggtggctgcattgaatgtggtgcgaaacatctttgagtggggcagcatctttggcagcgttgagtactttgagaagtaccgggaactggactgcctgccaccgctggtggaggctttgcaggaaatgcttccccactgtactttcaaagaacaggcggcactcaagcacttcctcatcgactgtttgcttcatcagggtgggcagtcacgacagttgctacaaacggtgcgaggatttggactgttctcgtcagatgaactccaggatattgacgaacagatgctgcaggcgacgccgccggtgcccaccaactcgttggcctcgctttccgactggatgtgtcactccaagatgagcagggttgacgtgggcgctctggagcgacaactgataagctgcaccaatgccaacacagtgaggatattgctagtgaaattatgcgcaactgctcctggaaaacctctgtgggccattaatcccagttgggacgtaccacagcccctgaaaactcttattatggccatgccggttagctttttgcaagacttcagctatgttctgctcggcaaggcgcgagaattggccacaaggggcaactacatcgatgccgtttcaatgctgagtgtactgaagtcggaaaatcagcgccaggagatggcggccaatgtgcagctgatgtgtaagctgatcacctgggaaatcctgcacattcagataacacagtgcctggaggagtggcaccagaagccactcgatctgcaatccttgggaggaagatgcaagcaatgtttgggagccctgcaggcaggagattcaatcgtcccacgtccggatatcctagagagttgcgctatcatgctacttaacctcacagaatttccacctctcctttacctggataaacgggcgggtcccctggaactacccctggccttcgccgccaccttcatcgagatggagaagatgaagggccccaagaaggtatgtcgcgatgcctgggaactgatgctaagcatgttccttaacgttccgaagcgtggctcctcaggagttggcggtatcagttccctgcaggccttcctacagcgaatccgccaccaaagcgttttcggactcgccatctccatgataggcaaagtccacaacatacttaaggatgaccccaatcatgatttaagctgcgagtacatgcagctgtggcctaccagcattaacaaccctgtcagctacagtttgaggagcgtttgcgagaccctgcaatggcttctttccgaggctttgtcctactatccgcaaaccatttcatggctgaagatgaaaggcgatctagacctggcgattggaaacaacgaatcggccatgcgctgttacgtgaacgccctcgtgaccggaacggattattgtacgatgcccctccaacgcaacgtcgcggatgattatgtcattcggaaaatgatccgctgcgcggcgaacctcggatgccatatgcaagcgacggtgctctgccaatttctggacgaaattgattatggaatcgtctttaaaaacctgtccgaaaagagcagcaactttacggacgcgatggacgcctactattcctgtatttgggacaccaccctcctcgaatttattgtgaacctccatgccaaacgcggagagcattcccgcaagctggaggccatctccatgatgggaaccctcgaactcaacgccaataataacgaagaaattaagcgcgagtccgcgatggtccgcaagagccgcttcctccgcgcgctcgcgaaacaatacctgctctaatctagagggcccgcgg

To generate **pUb FLAG-IntS9**, the following sequence was inserted between the BamHI and XbaI sites:

aatgcgattgtattgtctcagcggggacctggccaagccatgttatatcattaccttcaagggcctgcggattatgctggactgcggtctcacggagcagacagtcctgaatttccttccgctgcccttcgttcagtcactgaagtggtccaatctgcccaacttcgtgcccagtcgggatcatgatccccaaatggatggcgagttgaaggactgctgtggccgggttttcgtggactcgacgcccgagtttaatctacccatggacaaaatgctggatttcagcgaagtggatgtgatacttatctcgaattatcttaatatgctggccttgccgtatatcacggaaaacacggggttcaagggcaaagtctatgccacggagccaacgctgcagatcggccgatttttcctggaggagctggtggactatatcgaagtgtcgcccaaggcgtgcaccgcgcgcttgtggaaggagaagcttcatctgctgcccagtcccttgagcgaagcctttcgcgccaagaagtggcgcactatctttagtcttaaggatgtccaaggcagtctgtccaaggtgaccatcatgggctacgacgagaagctggacatcctcggagcttttatcgccactcctgtcagctcgggctactgcctgggctcgagcaactgggtactgagcacggcgcatgaaaagatttgctatgtcagtggttcctccacgttgaccacacatccgcggcccatcaatcagtcggcgctgaagcacgccgatgtactcattatgaccggactgacgcaggcgccgacggtaaatccggacacgaagctgggcgagctctgcatgaatgtggccttgacgatccgcaacaatgggtctgccttgattccctgttatccttcgggcgttgtctatgacctctttgagtgcctcacccagaatttggaaaacgctggcctaaacaatgttcccatgttctttatttcccctgtggcagacagctccttggcgtattctaatatcctggccgagtggcttagttccgccaagcagaacaaagtgtatcttccagacgatcctttcccacatgccttttaccttcgcaacaacaagctgaagcactataaccatgtgttctccgagggctttagcaaggacttccggcagccctgcgtggtcttttgtggccatcctagcttgcgtttcggcgacgctgttcactttatcgagatgtggggcaataaccctaacaactccatcatattcacggagccggactttccgtatctgcaagtactggcgcccttccaaccactggccatgaaggctttttactgtcccatcgacacctctttgaattatcagcaggcaaataagctgatcaaggagttgaaaccaaatgtgcttgtcatcccagaggcatacacaaaaccacatccctcggccccaaatttgttcattgaacagccggataagaagataataacgttcaagtgcggcgagatcatacgtctgccattaaaacggaagctggatcgaatttacattacctcggaactggcccaaaagatatcgccaaaggaggtggcggccggcgtgaccttctccacattgacaggagttttgcaagttaaggacaaggtccactgcatccagccatgtgcggatagcgttaaagacgagaccatctcgagcaacagcgctccgacaaaggaggatgtgctgaagaacgtcaaatacgagtatggcagcattgatgtagatgcagtgatgaagaagctggcacaggatggcttctccaacatcaagctagaccgcactggcggcgctctgactctgaaccttgtcaatgaggacacagtcattaagttcgaggacaacgagacgcatattatctgcggcggaaagccaacaacccggctcaagctgcgagacaccatcatgaaatgcttacagagtttttaa

To generate **pUb FLAG-IntS10**, the following sequence was inserted between the BamHI and XbaI sites:

aatgccgagccaagaggaaaatgagttgtacatggtcaaggaggcgcaaagactccggaaaagcgatccttgcgcggccatggcctggattatcacagctaaaactttatatcccaatgcattcaacctgcaatacgaggcttatctactggaacgcgatgctcagaattatgaggaggcagcaaagtgtttcagtgctatagccaccaatttccagaatcaacatacggagctgtggcaggagatcaactccctaacaaatgccctcagaaatgagaatgaaactacgccggaacacgagttctatgtaaaaatgtacaagcacctaactccggaagtacagcacaacatattcatgcacaccatcaaccacagtgccgataatctggagcgcatctatatctacatactgatgttcaacaagtttcccaaatcagccataactcaggctcccaggctactggaaatgctggcggagggcatgaaaaccgaaccggatctctatcagcggattctggtggaggaggtattgcccatgatccagaataagcccccggaactttcgccaaatctggcctgcagactttataccagttccttggagttttatttgcggcaaataatggatgaatcggatacagcagatgcctggaagaacatctttaaagtgctgatgatctgcggtcagatgatgggctgggagcccttcctgcccttcagcaagcatgtcaaccagaacgtctactgggaaaagctggtggacatactttccggcagtcctgcgggcagctcccaggttttgttctacgccaccaccttgttcatttactcccttcatggttatatacgaaattgtaagctgaggatcgaggatgccgatgtaactcatgttttggtagaaggattcatggaatggtcgccggaaggtgatggctccgaagtgcccagcatggagccaccgaaattctccctgaccacggccataagtccagagttatccaaggccttcctacacgccgcccagtgctggcagctgctcaatacggatcagttccaaagggatttcagtcaacttatgctggctcttcccttggcgccttggatctcaagattcctcttcgacttggccatatatttcggacaccgggatgaggccaacaagcttatggcggacatgaccacccagagcagtctggtgcaaagcctgcagatcttgagccttaatctaatgcaaggcagcatgacgctccagggcttccagtgcattttaaagatcctgtcagaactccccaccacccagggtcaacttttggagaatatgtcgctgaagggccacaggcacatggtttttctgcccctcactcgatcagcgttggttcagtactgcgtgggagccatcatcagcagactgagtcgcaaggtcttcgaaccgaacgttccagatcgattgctaggggatatcctagtgctgcagcaacttaatctactcaacgatgttctgctcactcaacagatatttaacctgatcaagcaacggaaatcgttcaatctgcgcaccctatccacctacattattaatatcgatttgctcgaggaactctcgcacatttggaactcccagcaggaggataactttgagttgaccagctcgccaaattcgagtggcacacctactgcgaccactgtagctggtggttcccaaagccgaagaattggcacccgtggagcggacaaaggagcccgagatgaattcagggcgataacacgccagcagattgcccgctgcaatgagaacgtgatcactttgcttgcgaatttcattaaccaagaacacttgatgctggcgcagcacatctttggaatcagtcagcccgtgggacgattgtgattaagtga

To generate **pUb FLAG-IntS11**, the following sequence was inserted between the SpeI and XhoI sites:

cccggacatcaagataacgcccttgggcgccggccaggatgtgggccgcagctgtctgctgctctcgatgggcgggaagaacataatgctcgactgcgggatgcatatgggctacaacgacgagcgccgcttcccggacttctcctacatagtgccggagggtccgatcaccagccacattgactgtgtgatcatctcgcacttccacctggatcactgcggggccttgccctacatgtcagagattgtgggctacacggggcccatctacatgacgcatccgaccaaggccatagcacccatcctcctggaggacatgcggaaggtggccgtagagcggaaaggcgagtcgaacttctttaccacccagatgatcaaggactgcatgaagaaggtgattcccgtcacacttcaccagagtatgatggtggacacggaccttgagatcaaagcctactatgcaggccatgtcctgggcgccgccatgttctggatcaaggtgggctcccagagcgttgtctacacgggggactacaacatgactccagacaggcatctgggagccgcctggatagacaagtgccggccggacttgctgatctccgagagcacctacgccactaccattagggactcgaagcggtgtcgcgagagggacttcctcaagaaagttcacgagtgcgtggctaagggcggcaaggttttaatccctgtttttgccctgggtcgcgcccaggagctgtgcatcctgctggagacgtactgggagcgcatgaacctcaagtaccccatatactttgctctgggtcttaccgagaaggccaatacctactacaaaatgttcatcacctggacgaaccagaagatccgcaagaccttcgttcaccgcaacatgttcgacttcaagcacatcaagccctttgacaaggcctacatcgacaatcctggcgcgatggtagtgttcgccacgccaggcatgctgcacgcaggtctttccctgcagatcttcaagaagtgggcgccgaatgagaacaacatggtgattatgcccggctactgtgtgcagggaaccgtgggcaacaagattctcggtggcgccaagaaggtggagttcgagaaccgccaggtggtcgaggtcaaaatggccgtggagtacatgagcttctcggcccatgcagatgccaaaggcattatgcagctaattcagaactgtgagccaaagaacgtcatgctcgtccatggtgaagcagggaagatgaagttcctgcgctcgaagatcaaagacgaattcaacctggaaacctacatgccggccaacggcgaaacctgtgtgatttctacgcctgtgaagataccagtagatgcgtccgtttccctgcttaaagcagaggcccggtcctacaacgcccagccaccggatcccaagaggcggcgactaatccatggcgtcctcgtaatgaaggacaatcgaataatgctgcaaaacctaaccgatgccctgaaggaaatcggaattaatcgacatgttatgcgatttacatccaaggtcaaaatggacgactccggaccggtcattcgcacaagcgaaagactgaagactctgctcgaagaaaagctagccggctggacggtgacgatgcaggagaacggctccatagccatcgaatccgttgaggtgaaggtggaggaggacgagaaggatcccaagcagaagaacattttgatatcgtggaccaaccaggacgaggacattggagcctacatactgaatgtgctgcagaatatgtgctag

To generate **pUb FLAG-IntS12**, the following sequence was inserted between the SpeI and EcoRI sites:

Cgcggccaacatcgcggcggccgccgccgccgcgcaggaggtcgacccggtcctcaaaaaagcgattaaactgctccatagctccaacccgacgagcgccgcggagctccgcctcctgctggatgaagccctgaaagcccgcttcggccccgaaaagtccctgacgaataatatgaccccccgcatgctcgaagacgaggccaatttctcgggccgcgccgcgaccccgccccaacaaccgattaacgccgatgaaattattaacctgactaattcccccgataaagaaccgtcggacagcgtcgataccatcgccgacagcgacgatggcctgtccgcggtcggcatcgtgaatacgggcgatacgggagatttcggagatctgaactgttgtgtctgtggagaaatggtgtttacggccaccaatcggctgattgagtgctccaagtgcggtgccatgtaccatcaggagtgccacaagccgcccataaccaaggaggaggcggccgatgaccaggagcagaactggcagtgcgacacgtgctgcaacaagccaacgagcagcgggaggacaacatcctctgcagcggccgtaacgccaactgtcttcatagccgacgaacccatgccgttgaccagcaaagccaaatcgtcggtcgcgtcgtcgcgctcatcgaactcttccaactcctcgtcacccttctaccggcccgagcccagcagttccacgaatgccagcagcagtagcagcagcaagcatggccacaagtcctcctcttcgtcgtcgtccaaatcgcacaaggaagaacgatcctctaagtccacggcggcgtcttctcttagcgccatcggcggaatggagaagcacaacagcagtggcacctcatcgcgacgcagcggctccagcaccaaatccagctccaagagcagttcctccaagcatcacgaaagcggcagcagcagcaagcgcagatccaagcagtaa

To generate **pUb FLAG-IntS13**, the following sequence was inserted between the EcoRI and XhoI sites:

tgttcgaacgcaaccagaagaccatctttgtgctggaccacacccgatactttagcatcgccagcgaggagtacatctcgatggacttcctaaagggaaaaccatctgcagacggcggcgcaacaggagcggcgggaaatgcaaccggcagcggtggatcccaattcagcaagagcctgtggacatgcgcctgcgaatcctccatcgagtactgccgcgtggtctgggatctctttcccggcaaaaagcatgtgcggtttattgtctcggacacggcggcgcatatcgtgaacacctggaggcccagcacgcaaaatatggcgcacgtgatgaacgcaatgctaattgtgggcgtgccgtcgcgcaacgtacccacatcttctgactactcggtaatccatggcctgcgggccgccatcgaagcgctggcggagcccaccgatgagcagttggcggcaatggccgactttgggaccgacgaactgccgcgcattcccaacaagggtcgcgtcatctgcataacttcagcgcgtgacaataccagcatgaaaagcctggaggacatctttaacaccgtgctggtgcagcagaacaccctagcagctccgccctccaagaagggactggtcatcgaccattgccacctagttatcctcaacattgtgccgctgggcgttgaatcgttggttaccaatcgcagcctgctcaagatatcgccgctgcttgatgtcgagatccacacagtcagtgccccggatatttcctataagctgacgcacctcatactgaaccattacgacctagctagtaccacggtgactaacatacccatgaaagaggagcagaacgccaactccagcgccaactacgatgtggaaatcctgcacagccgaagggcccactccatcacctgtggcccagatttcagtctacccacgagcatcaagcaaggtgccacatacgaaacagtgactcttaagtggtgcacaccgcgcggttgcggctctgcagatctacagccctgtctgggtcagttcctcgtgacaccggtggacgtcacctcgcggcccagttcttgcttgattaactttctgcttaacggacgttcagtcttgctggaaatgccgcgcaagaccggctcaaaggccaccagccatatgctttctgcacgcggtggcgagatctttgtgcactccctgtgcatcacgcgctcctgcatggatgaggctccatcaattactgatggccccggaggacgtgtctcggactatcgcaccgccgaacttggtcaactgatcaagatgtcgcgcgtggttccgctcaaggttaaggatccatctgctccgcccttgacgcgccggttgcctcgctacttccccctgaccaccagctcttcgattctgttccatctacagcgacacatcagttggcttccgcattttctccatcttttagtgaaggaagacatggataagcaggacgaggtgcggtgccagcagcacattcacgagctgtacaaaagcgcctcgcgtggcgatgtcctaccatttacccacaccaatggagcaagactaaagctttccaaggccaaagaccagtaccgcttgctgtacagggaactggagcaattaattcaactaaatgccaccacaatgcaccataaaaatctgctggaaagcctacagagtttgcgagccgcatacggcgatgccccgctgaagtcggaacctggagcaagtctcctgcgcacctacacggaatccccactctctcccgaacggctggagcccatcagtagcgtcggtgctagtggcagcagcagctccaatagccttctcaaggcgagcaagcgtcgcatgtctagttgtgggcaaagatccctgctagatattatatcctcagcagagcgaagccagtccaacaaacgattggatttctcgggacgcctttgcactccgttgggtcaagtggccaagctgtatccggattttggtaccaaggacaaagacacagttacgacgggggccagtattacgcccaatgttaaggaggaatccgtacgtagttaa

To generate **pUb FLAG-IntS14**, the following sequence was inserted between the SpeI and NotI sites:

ccccaccttaatagcgctggatgcatcgctatcgatgctgcgcccggtgccgggaagaaatgagcacacctaccagtcgctggccaccaagggcatccagcatctgctggacaatctcacggcggccggcaaattggagcacgttgctctgctctcctattcgacgacggcggaactgaaggtggacttcacgcgggactacgaccaggtgcggcaggcggtcaagaaggtggagcccgtggacaaggcctgcctgatgagcatgctcaaggcggtggtgtccataatgtcgccgtggggcaaccagaacatcctgcaggtggttgtctttacggactgcggcctcggctttggcaacacctcgatcacgggcttcctggaggcctacgccgaaaaggagtcggagccggagttcggcttcctgaagactctggccaactacaacctgaatttcatctgcctgggcctgcacggcgattactacttcaccaggggattggcggtgtaccagcagctgctggacaaggtctcactgaagggccagctcttcatgacaaagcccgccaagagcagcgatgcagtcgagggaaatcccaatcccaacccgaaccccagccacaaaagcgaactgggccgcactacggtcttcgaactaattgaacgtctctgcgaggccagctacaagagttcggaggtgaccctgaaatgcggcagctacttccgcatggaggcatctgttctgctctggccaccgacggctccgtatgagcaaaagtcccacatattcggccgagaacccaccattcgccacacagaccagaagattgaggtgtgtggcttcctgtccctttcggacattggatcaccggccaccctgagtcgacactgggtgctgcccaaagtggagcgggagaagagcggcagcagtcggaggtctggaaacctgagcgcagcagccaagccaccgaagctcaatctagataccagcaatcccaactacgagctggagaagctggaagccgatatcaaagagttttacgccaaggattcaaaagacacggaggaaagtggcgacgacgatgtgaccattgtcttgaaacccggcccccagacagagcaacaaaaagaaaacttgtgcgtcctccttcacggagccctcaaaatggagaacatggcagctctggttcgagtgggtgacaagtggtacggattcatatatgcattcaccgacagcaagaagaagtcaaacctgatgctgaacatacttccacccggcacgaacgtgattccctggctaggagacctggaatcgctgggcttccccgaggatttggcgcccggagagacggccagttttccggtgcgtgccgaccgcagatcttattcgcagagcagtgtggtgtggatacgacaggctagcctgcagtccgacgtccagaaggtactgcgacacgccaagaagatgcccgacaagacgcagcacttctacaaggagctcaatcgcatccgccgcgctgccctggcgctgggattcgtcgagttgctggaggctttggccatgctgctggagaaggagtgcgcccatctcagcctgaacggagccagcaacgactgcaccctgcagctgcagcacgcggctacggagttgcggaagacctcgaacagagacatgaagtccatgattgttccgctccagaaagttggtgcatcggatgcgggcagcacggctgccacggctcctgctcctgcatacatgtattga

The following inducible *Drosophila* expression plasmids were generated from the previously published **pMT FLAG MCS puro** plasmid (Elrod et al. (2019) *Mol Cell*, 76, 738-752):

**pMtnA FLAG-IntS6 puro	(Addgene #195076)**

**pMtnA FLAG-IntS12 puro	(Addgene #195077)**

The MtnA promoter (pMtnA or pMT; marked in gray) drives expression of an N-terminal 3x FLAG tag (marked in blue). Start codon is marked in green. The full sequence of **pMT FLAG MCS puro** is as follows:

gcgtcaatacgggataataccgcgccacatagcagaactttaaaagtgctcatcattggaaaacgttcttcggggcgaaaactctcaaggatcttaccgctgttgagatccagttcgatgtaacccactcgtgcacccaactgatcttcagcatcttttactttcaccagcgtttctgggtgagcaaaaacaggaaggcaaaatgccgcaaaaaagggaataagggcgacacggaaatgttgaatactcatactcttcctttttcaatattattgaagcatttatcagggttattgtctcatgagcggatacatatttgaatgtatttagaaaaataaacaaataggggttccgcgcacatttccccgaaaagtgccacctgacgtctaagaaaccattattatcatgacattaacctataaaaataggcgtatcacgaggccctttcgtctcgcgcgtttcggtgatgacggtgaaaacctctgacacatgcagctcccggagacggtcacagcttgtctgtaagcggatgccgggagcagacaagcccgtcagggcgcgtcagcgggtgttggcgggtgtcggggctggcttaactatgcggcatcagagcagattgtactgagagtgcaccatatgcggtgtgaaataccgcacagatgcgtaaggagaaaataccgcatcaggcgccattcgccattcaggctgcgcaactgttgggaagggcgatcggtgcgggcctcttcgctattacgccagctggcgaaagggggatgtgctgcaaggcgattaagttgggtaacgccagggttttcccagtcacgacgttgtaaaacgacggccagtgaattaattcgttgcaggacaggatgtggtgcccgatgtgactagctctttgctgcaggccgtcctatcctctggttccgataagagacccagaactccggccccccaccgcccaccgccacccccatacatatgtggtacgcaagtaagagtgcctgcgcatgccccatgtgccccaccaagagctttgcatcccatacaagtccccaaagtggagaaccgaaccaattcttcgcgggcagaacaaaagcttctgcacacgtctccactcgaatttggagccggccggcgtgtgcaaaagaggtgaatcgaacgaaagacccgtgtgtaaagccgcgtttccaaaatgtataaaaccgagagcatctggccaatgtgcatcagttgtggtcagcagcaaaatcaagtgaatcatctcagtgcaactaaaggggggatctagatcggggtaccatggactacaaagaccatgacggtgattataaagatcatgatatcgattacaaggatgacgatgacaaggatccactagtccagtgtggtggaattctgcagatatccagcacagtggcggccgctcgagtctagagggcccgcggttcgaaggtaagcctatccctaaccctctcctcggtctcgattctacgcgtaccggtcatcatcaccatcaccattgagtttaaacccgctgatcagcctcgactgtgccttctaagatccagacatgataagatacattgatgagtttggacaaaccacaactagaatgcagtgaaaaaaatgctttatttgtgaaatttgtgatgctattgctttatttgtaaccattataagctgcaataaacaagttctagggtcggtgggcctcgggggcgggtgcggggtcggcggggccgccccgggtggcttcggtcggagccatggggtcgtgcgctcctttcggtcgggcgctgcgggtcgtggggcgggcgtcaggcaccgggcttgcgggtcatgcaccaggtgcgcggtccttcgggcacctcgacgtcggcggtgacggtgaagccgagccgctcgtagaaggggaggttgcggggcgcggaggtctccgaggaaggcgggcaccccggcgcgctcggccgcctccactccggggagcacgacggcgctgcccagacccttgccctggtggtcgggcgagacgccgacggtggccaggaaccacgcgggctccttgggccggtgcggcgccaggaggccttccatctgttgctgcgcggccagccgggaaccgctcaactcggccatgcgcgggccgatctcggcgaacaccgcccccgcttcgacgctctccggcgtggtccagaccgccaccgcggcgccgtcgtccgcgacccacaccttgccgatgtcgagcccgacgcgcgtgaggaagagttcttgcagctcggtgacccgctcgatgtggcggtccgggtcgacggtgtggcgcgtggcggggtagtcggcgaacgcggcggcgagggtgcgtacggcccgggggacgtcgtcgcgggtggcgaggcgcaccgtgggcttgtactcggtcatggaaggtcgtctccttgtgaggggtcaggggcgtgggtcaggggatggtggcggcaccggtcgtggcggccgacctgcaggcatgcaagctatccaattcctgcagcccgggggatctgttgtaatttataatttatatttcctttcttaataaataaataaatagtcaagtttatgtttgagttttatgatttatattttaagttatttcaactgcaacaccagcaccacgacctacttacagcaaaaaacgtacaagaaggaaagaaggaataaaaagagtggtattctcttacaatatgttttatggcataaaaggtgtggccattcatatcaaatataaagtagtgttgtttaacgttacttttgtaggttgaatagtatattccaacagatgatgaggggttcccaatcctaaacccatttgccgttcccagaagcatgaaaccaccacgcacccgatcctctaggacaacaacaattgcattcattttatgtttcaggttcagggggaggtgtgggaggttttttaaagcaagtaaaacctctacaaatgtggtatggctgattatgatcagtcgacctgcaggcatgcaagcttggcgtaatcatggtcatagctgtttcctgtgtgaaattgttatccgctcacaattccacacaacatacgagccggaagcataaagtgtaaagcctggggtgcctaatgagtgagctaactcacattaattgcgttgcgctcactgcccgctttccagtcgggaaacctgtcgtgccagctgcattaatgaatcggccaacgcgcggggagaggcggtttgcgtattgggcgctcttccgcttcctcgctcactgactcgctgcgctcggtcgttcggctgcggcgagcggtatcagctcactcaaaggcggtaatacggttatccacagaatcaggggataacgcaggaaagaacatgtgagcaaaaggccagcaaaaggccaggaaccgtaaaaaggccgcgttgctggcgtttttccataggctccgcccccctgacgagcatcacaaaaatcgacgctcaagtcagaggtggcgaaacccgacaggactataaagataccaggcgtttccccctggaagctccctcgtgcgctctcctgttccgaccctgccgcttaccggatacctgtccgcctttctcccttcgggaagcgtggcgctttctcatagctcacgctgtaggtatctcagttcggtgtaggtcgttcgctccaagctgggctgtgtgcacgaaccccccgttcagcccgaccgctgcgccttatccggtaactatcgtcttgagtccaacccggtaagacacgacttatcgccactggcagcagccactggtaacaggattagcagagcgaggtatgtaggcggtgctacagagttcttgaagtggtggcctaactacggctacactagaagaacagtatttggtatctgcgctctgctgaagccagttaccttcggaaaaagagttggtagctcttgatccggcaaacaaaccaccgctggtagcggtggtttttttgtttgcaagcagcagattacgcgcagaaaaaaaggatctcaagaagatcctttgatcttttctacggggtctgacgctcagtggaacgaaaactcacgttaagggattttggtcatgagattatcaaaaaggatcttcacctagatccttttaaattaaaaatgaagttttaaatcaatctaaagtatatatgagtaaacttggtctgacagttaccaatgcttaatcagtgaggcacctatctcagcgatctgtctatttcgttcatccatagttgcctgactccccgtcgtgtagataactacgatacgggagggcttaccatctggccccagtgctgcaatgataccgcgagacccacgctcaccggctccagatttatcagcaataaaccagccagccggaagggccgagcgcagaagtggtcctgcaactttatccgcctccatccagtctattaattgttgccgggaagctagagtaagtagttcgccagttaatagtttgcgcaacgttgttgccattgctacaggcatcgtggtgtcacgctcgtcgtttggtatggcttcattcagctccggttcccaacgatcaaggcgagttacatgatcccccatgttgtgcaaaaaagcggttagctccttcggtcctccgatcgttgtcagaagtaagttggccgcagtgttatcactcatggttatggcagcactgcataattctcttactgtcatgccatccgtaagatgcttttctgtgactggtgagtactcaaccaagtcattctgagaatagtgtatgcggcgaccgagttgctcttgcccg

To generate **pMtnA FLAG-IntS6 puro**, the following sequence was inserted between the SpeI and XhoI sites:

cacaatcatactcttcctggtggacacctcgtcgtccatgtgccagaaggcgtatgtgaatggggtacagaaaacgtatctggacattgccaagggagccgtggagacgtttctcaagtatcgccagcgtacgcaggattgcctgggagatcgctacatgctgctcacattcgaggagccaccggcaaacgtgaaagctggatggaaggagaaccatgccaccttcatgaacgagctgaagaacctgcagagtcacggcctcacctcgatgggtgaatcgctgcgcaatgcgttcgatttgttaaatctgaatcgcatgcagtcgggcatcgatacgtacgggcagggcaggtgcccgttttatctggagccatcggtcatcattgtgattacggacggcggtcgctattcgtaccggaatggtgtccatcaggagatcatactgccgctaagtaaccaaataccgggcacaaagttcactaaggagccatttcgctgggatcagcgcttgttttcgcttgtcctccgcatgccgggcaacaagattgacgagcgagtggatggcaaggtgccgcatgacgattctcccatcgaacggatgtgcgaggttaccggcggacgttcatatcgagtgcgaagccactacgtactcaaccagtgcattgaaagtcttgtccaaaaggttcagcctggtgtggtgctgcagttcgagcctatgctgcccaaggaggctacttccgctaccgctggcgaagcggcgggtgcgagcacgatatccggcatgggaataccctcaacgtccggagcgggtcccgctcccgatattgtcttccatccggtaaagaagatgatctacgtgcagaagcacatcacgcagaaaacctttcccatcggctattggccactgccggagccctattggccggactccaaggcgattacattgccgcctcgcgacgctcatcccaagctgaaggtcctgacgccggcggtggacgagccacagctggtgcgcagcttccccgtcgacaagtacgagattgaaggctgtccgttaacgctgcagattctcaacaagcgcgagatgaacaagtgctggcaggtgattgtgaccaatggcatgcatggattcgagctgccctttggctacctaaaagcggcgcccaacttttcgcaggtacacctctatgtactggcctacaattatccagcgttgctgcccattctgcatgacctcattcacaagtacaatatgagtccgccgagcgatctgatgtacaaattcaacgcctatgtgcgttcaatacccccgtattactgcccgttcctgcgcaaggcgctggtcaacatcaatgtgccgtatcagctgctgcagtttctgctgccggagaatgtggacaactatctgtcgccaacgatcgccaatcagcttaagcacatcaagaatacggccaagcaggatcaggagaatctttgcatgaaggtgtacaagcagctgaagcagccgaagccaccatatcgccaagtggagacgggcaaactgttcacaggtgcgccgttacgcagagatttggtgcggcatccactgctccgggatatattttccaagttgcatggggaaatagatcccgtggagaattatacgattgtggtgccgcagccaacgcatcagtcgtcggccaagtcgcatcgcaatcccttcgatataccgcggcgggatttggtggaggaaatagccaagatgcgtgagaccttactgcgacccgtctcgctggtggccaaggactcgggccattgcctgcccatcgccgagatgggcaactaccaggagtacctcaagaacaaggacaatccgctgcgtgaaatcgagccgacgaatgtgcggcagcacatgttcggtaatccgtacaagaagaacaagcacatggtgatggtggacgaggcggatctgagcgatgtggcgccaatgaagtcgccaaatggcaatggaccgggtggagcgccacccggctcaccttccggatcgggctcgcccacaggaccgggtccctcatctccgggctcttcgcccggcggtggttcgggtcccgggatgcccggaatgcccggaatgggcggtggaatgtcaggactgatgctgggcgcaggtggcagtggcggatcctctaaaaaactggacggcacgtctcgtgggcgaaagaggaaagcaggcccgctgccacgtagctttgaattccgtcgatcatctacggactcgcgatcttcgagctccggctccgaattatccacaacgggtagtgcaccaggatcaccgataccaggcgccacgtcatccatatccggctgcgattccgccatgggcgctgatggcggtcccagttccagctgtttcgacgaggacagcaatagcaacagcagcttcgtgagctccacatcggaggcctcggccagtgattcagggatgtcaaactcgcatttggatcccttgcgaatcagtttcgttttcatcgaggaggagaccagcgatcaggatacgccgctcaatggttatgtgcacaatttcatcaatggcatcagcgacgatgcagaaacgggcgaaacggaagcagttccaggcggggcatccctgccgggtgcgtcttcagctaacgaaccgtcgtcaattggagcatcaccagcagtttcagccacgccagcaacatctgtgatccccgccagcaatggcagcgaagtttgctccgccgtcgcggccggcagcagcagttccatcagctccagcgccagttccccgggcgttctctacgtgcccctcaacggactgaccacccatgcggcctacagcagtaatctgggtggtccgagctcgccgttggacaaccttagctccctgggtggagcggcgggtggcggaggatcgggactactcggtgtcagcaagccgggatcgctgccactggccaatggttatggacacggtcatagttcagccatttccccaccttcgccattgtccttggtgcctctatcaccattgggaacaagtacgcccgctgccagcagctcgggactatcgtctagttccgcctacttcagccacaatgacatcaacgatgcctcggagatctcgcgcatactgcagacctgtcacaatggaaccagcagtggctcctcggtcggagcgggcgcaagcaacagcaatttgaatggcaatggcagcacggagagtggcggaggagtctcatccgacgatcatgcctcgctgggtggcaacagtttcgatgccctgaaacggacagcgggtactgcgggcagcactgcagcgggaaatccccactactactgggccagtggtctcaatcatcacaacaataacaacagcagcgccgctggagcagctagcggttcgacgttgagcaacaatcacaaccaccggggtcacaacaacagcagccccaacaaactggaggtgaagatcaactccagctgcggctcctcaccgacgcacaacaacgggggcatgggctcaccgggacagagcctgccgctcattttcaccgaggagcagcgcgaagcggcccggttgcacaacgtcgaactacgattgcagatcttccgtgatatacggcgacccggccgagactacagccagctgctggagcacctgaatctggtcaagggcgaccaggatatgcagtcggatttcgtggacatgtgtatcgtggaatcgctgcgcttccgccgccatcgcatggcgtccagcatacaggagtggtgggatcgcaagcagcagttaactgccgtgggaaccaatgaagcgacccccagcgcggagcaggccgtcgccaagagttaa

To generate **pMtnA FLAG-IntS12 puro**, the following sequence was inserted between the SpeI and EcoRI sites:

cgcggccaacatcgcggcggccgccgccgccgcgcaggaggtcgacccggtcctcaaaaaagcgattaaactgctccatagctccaacccgacgagcgccgcggagctccgcctcctgctggatgaagccctgaaagcccgcttcggccccgaaaagtccctgacgaataatatgaccccccgcatgctcgaagacgaggccaatttctcgggccgcgccgcgaccccgccccaacaaccgattaacgccgatgaaattattaacctgactaattcccccgataaagaaccgtcggacagcgtcgataccatcgccgacagcgacgatggcctgtccgcggtcggcatcgtgaatacgggcgatacgggagatttcggagatctgaactgttgtgtctgtggagaaatggtgtttacggccaccaatcggctgattgagtgctccaagtgcggtgccatgtaccatcaggagtgccacaagccgcccataaccaaggaggaggcggccgatgaccaggagcagaactggcagtgcgacacgtgctgcaacaagccaacgagcagcgggaggacaacatcctctgcagcggccgtaacgccaactgtcttcatagccgacgaacccatgccgttgaccagcaaagccaaatcgtcggtcgcgtcgtcgcgctcatcgaactcttccaactcctcgtcacccttctaccggcccgagcccagcagttccacgaatgccagcagcagtagcagcagcaagcatggccacaagtcctcctcttcgtcgtcgtccaaatcgcacaaggaagaacgatcctctaagtccacggcggcgtcttctcttagcgccatcggcggaatggagaagcacaacagcagtggcacctcatcgcgacgcagcggctccagcaccaaatccagctccaagagcagttcctccaagcatcacgaaagcggcagcagcagcaagcgcagatccaagcagtaa

The following protein expression plasmids used for antibody production were generated from **pRSFDuet-1** (Novagen):

**pRSFDuet-1 IntS6 AA 1035-1284	(Addgene #196904)**

**pRSFDuet-1 IntS8 AA 1-308	(Addgene #196905)**

**pRSFDuet-1 IntS11 AA 300-597**	**(Addgene #199329)**

To generate **pRSFDuet-1 IntS6 AA 1035-1284**, the following sequence was inserted between the BamHI and HindIII sites:

gacatcaacgatgcctcggagatctcgcgcatactgcagacctgtcacaatggaaccagcagtggctcctcggtcggagcgggcgcaagcaacagcaatttgaatggcaatggcagcacggagagtggcggaggagtctcatccgacgatcatgcctcgctgggtggcaacagtttcgatgccctgaaacggacagcgggtactgcgggcagcactgcagcgggaaatccccactactactgggccagtggtctcaatcatcacaacaataacaacagcagcgccgctggagcagctagcggttcgacgttgagcaacaatcacaaccaccggggtcacaacaacagcagccccaacaaactggaggtgaagatcaactccagctgcggctcctcaccgacgcacaacaacgggggcatgggctcaccgggacagagcctgccgctcattttcaccgaggagcagcgcgaagcggcccggttgcacaacgtcgaactacgattgcagatcttccgtgatatacggcgacccggccgagactacagccagctgctggagcacctgaatctggtcaagggcgaccaggatatgcagtcggatttcgtggacatgtgtatcgtggaatcgctgcgcttccgccgccatcgcatggcgtccagcatacaggagtggtgggatcgcaagcagcagttaactgccgtgggaaccaatgaagcgacccccagcgcggagcaggccgtcgccaagagttaa

To generate **pRSFDuet-1 IntS8 AA 1-308**, the following sequence was inserted between the BamHI and HindIII sites:

atggacgatcctcttaagcccaagccagttccactggctgccgaaacggtactgtggttcgagttcctgctcgatccgcacaagattacgcagcatctgcagcgcccgcatcccgagcccagcgccatggagctgatcgtgcagttcatcagcatgacgccgaacacggcgcaggagtcagtggggacgcctggcagtgatttgcagaatctaaaccagacgccatcgaattcgggacccattcccggcgtggttggtggtgcccctgcgccgacaacgcccactgcctccggcggagtgggcatgccccatagcccacaaaggcccgcggagaagggcctgcaattgaaccgcaagcagttggcactgaaaatcctcgagctgaaggtagccacttggctgaagtgggatctggatgcattggagaagaacctgcctgtgattatgcagctggccttgctgcgcgatctgtgcaccataagctacggctgctccttaagtattcccctgcccaatgattttgacgcgagaatttccgctgctggaaacgaaagagcagcaagattcgccttgaccatataccaccgcatgctgctgcgaatgcaactgataaaggagcaggcgttaaaagcgccacgtcctcaaaacacaatgtaccaaactgtggaccagctccagcagtttctggacacaccgactcaaccatccatcgagtatttgcaacaactctgcgcctcgacaaaacctttctacatatttcactacgacagctttgttccgctgcgatgtgatgatatcggcaatggccaaaactacgatgttatgcacctaataacaccccaggaattgagagcccaactgcactacgagctcgcccaatattacctgtacaccaaacagtatgttttggcccgggaagcggcctaa

To generate **pRSFDuet-1 IntS11 AA 300-597**, the following sequence was inserted between the SalI and HindIII sites:

ttcgacttcaagcacatcaagccctttgacaaggcctacatcgacaatcctggcgcgatggtagtgttcgccacgccaggcatgctgcacgcaggtctttccctgcagatcttcaagaagtgggcgccgaatgagaacaacatggtgattatgcccggctactgtgtgcagggaaccgtgggcaacaagattctcggtggcgccaagaaggtggagttcgagaaccgccaggtggtcgaggtcaaaatggccgtggagtacatgagcttctcggcccatgcagatgccaaaggcattatgcagctaattcagaactgtgagccaaaaaatgtcatgctcgtccatggtgaagcggggaagatgaagttcctgcgctcgaagatcaaagacgaattcaacctggaaacctacatgccggccaacggcgaaacctgtgtgatttctacgcctgtgaagataccagtagatgcatccgtttcactgcttaaagcagaggcccgctcctacaacgcccagccaccggatcccaagaggcggcgactaatccatggcgtcctcgtaatgaaggacaatcgaataatgctgcaaaacctaaccgatgccctaaaggaaatcggaattaatcgacatgttatgcgatttacgtccaaggtcaaaatggacgactccggaccggtcattcgcacaagcgaaagactgaagactctgctcgaagaaaagctagccggctggacggtgacgatgcaggagaacggctccatagccatcgagtccgttgaggtgaaggtggaggaggacgagaaggatcccaagcagaagaacattttgatatcgtggaccaaccaggacgaggacattggagcctacataccgaatgtgctgcagaatatgtgctag
