## Supplemental Figures 1-7 for "IntS6 and the Integrator phosphatase module tune the efficiency of select premature transcription termination events"

### Supplemental Figure S1

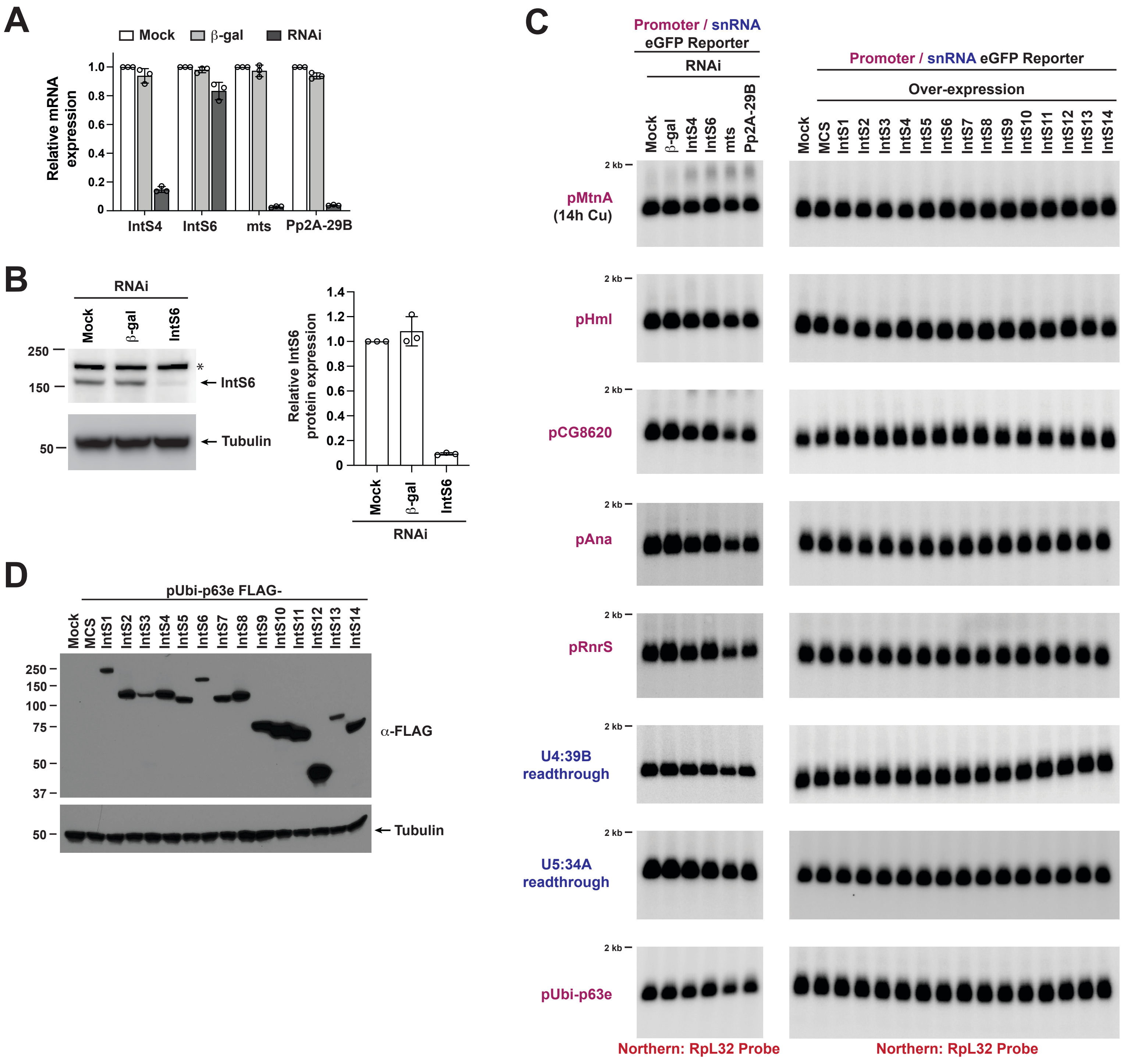

### Supplemental Figure S2

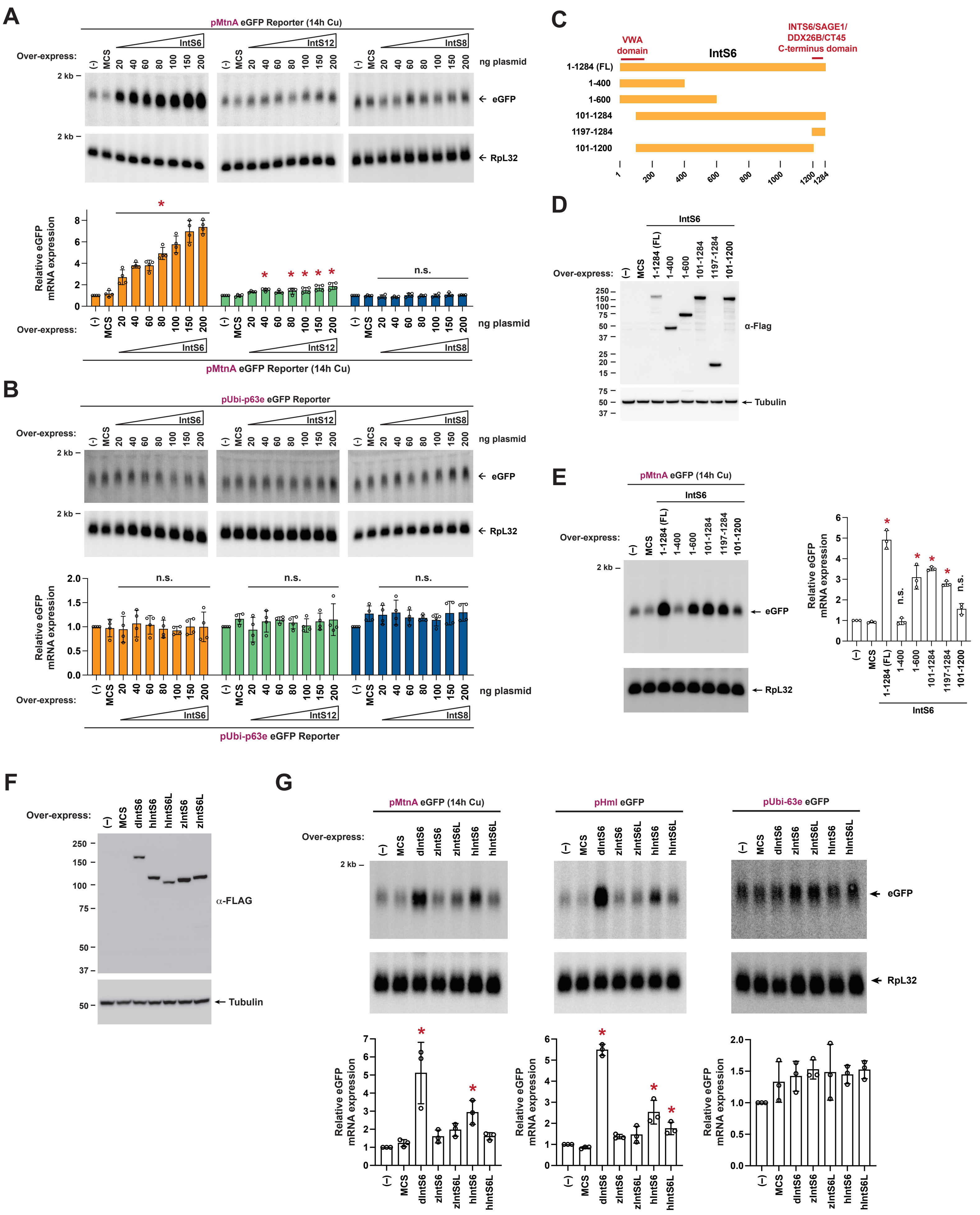

### Supplemental Figure S3

A

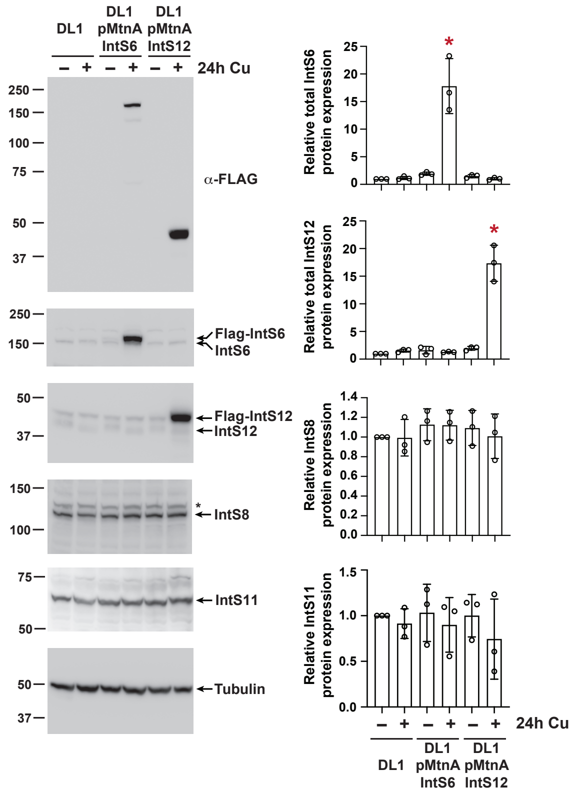


B

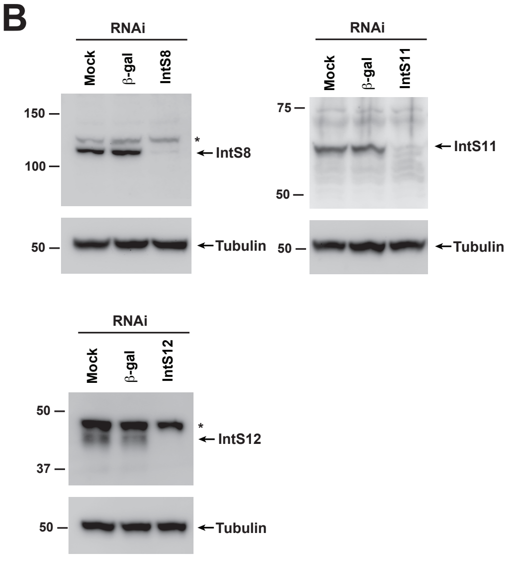


C

| Cell line | Treatment | Replicate | Total raw fragments | Total raw fragments after QC | Mappable fragments | % of total fragment after QC |
|---|---|---|---|---|---|---|
| DL1 | –Cu | A | 20,405,648 | 20,184,594 | 19,065,747 | 94.46% |
| | +Cu | | 18,838,707 | 18,566,085 | 17,486,883 | 94.19% |
| DL1 pMnA IntS6 | –Cu | | 17,949,058 | 17,728,039 | 16,718,665 | 94.31% |
| | +Cu | | 22,018,534 | 21,734,111 | 20,425,890 | 93.98% |
| DL1 pMnA IntS12 | –Cu | | 22,668,256 | 22,421,152 | 21,143,465 | 94.30% |
| | +Cu | | 21,081,347 | 20,799,709 | 19,600,773 | 94.24% |
| DL1 | –Cu | B | 21,431,925 | 21,184,885 | 20,020,634 | 94.50% |
| | +Cu | | 24,310,414 | 24,016,328 | 22,705,998 | 94.54% |
| DL1 pMnA IntS6 | –Cu | | 20,968,479 | 20,696,252 | 19,511,734 | 94.28% |
| | +Cu | | 18,762,373 | 18,523,988 | 17,448,283 | 94.19% |
| DL1 pMnA IntS12 | –Cu | | 20,566,105 | 20,302,204 | 19,181004 | 94.48% |
| | +Cu | | 20,015,012 | 19,752,161 | 18,636,567 | 94.35% |
| DL1 | –Cu | C | 19,063,955 | 18,808,889 | 17,742,723 | 94.33% |
| | +Cu | | 18,712,101 | 18,451,096 | 17,393,283 | 94.27% |
| DL1 pMnA IntS6 | –Cu | | 20,088,379 | 19,846,941 | 18,627,901 | 93.86% |
| | +Cu | | 19,717,306 | 19,469,650 | 18,341,313 | 94.20% |
| DL1 pMnA IntS12 | –Cu | | 19,028,561 | 18,772,741 | 17,674,543 | 94.15% |
| | +Cu | | 20,248,031 | 20,001,754 | 18,884,783 | 94.42% |

D

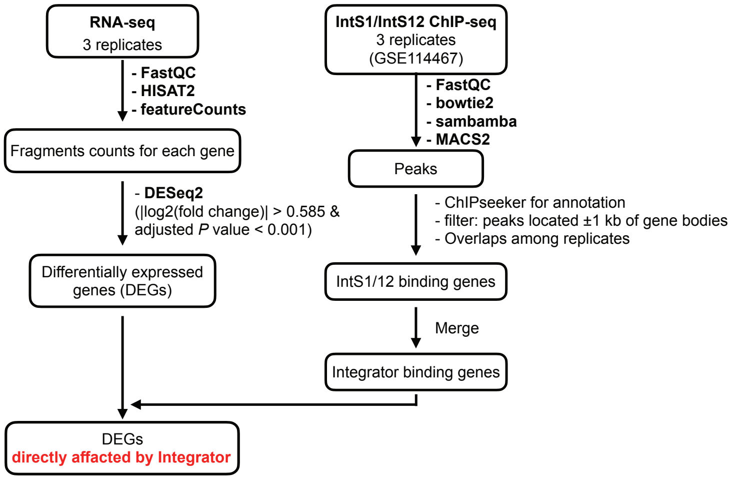


E

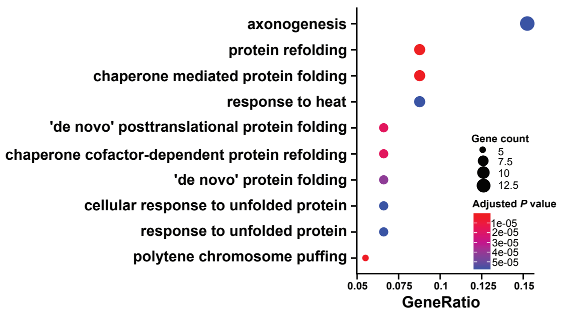


GO enrichment analysis for 107 up-regulated genes with IntS6 over-expression

F

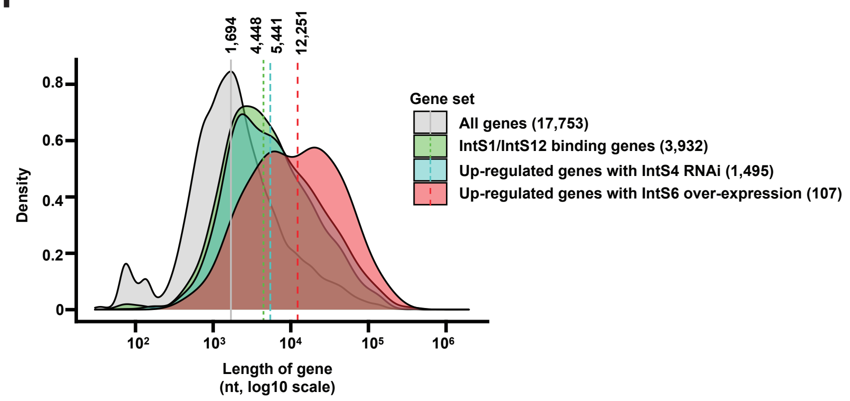

### Supplemental Figure S4

**A**

| Cell line | Treatment | Total raw fragments | Total raw fragments after QC | Mappable fragments | % of total fragment after QC |
|---|---|---|---|---|---|
| DL1 | Input ChIP-seq Rep 1 | 26,228,635 | 25,516,367 | 24,884,439 | 97.52% |
| | Input ChIP-seq Rep 2 | 27,223,555 | 26,367,428 | 25,755,453 | 97.68% |
| | Input ChIP-seq Rep 3 | 22,491,124 | 21,736,365 | 21,196,214 | 97.51% |
| | IntS12 ChIP-seq Rep 1 | 35,932,364 | 34,713,094 | 31,455,232 | 90.61% |
| | IntS12 ChIP-seq Rep 2 | 31,827,197 | 30,271,670 | 28,352,062 | 93.66% |
| | IntS12 ChIP-seq Rep 3 | 39,777,104 | 38,581,220 | 36,156,923 | 93.72% |
| | IntS1 ChIP-seq Rep 1 | 27,130,554 | 26,304,363 | 24,578,979 | 93.44% |
| | IntS1 ChIP-seq Rep 2 | 31,488,028 | 30,545,475 | 27,902,535 | 91.35% |
| | IntS1 ChIP-seq Rep 3 | 34,358,643 | 33,118,176 | 31,299,358 | 94.51% |

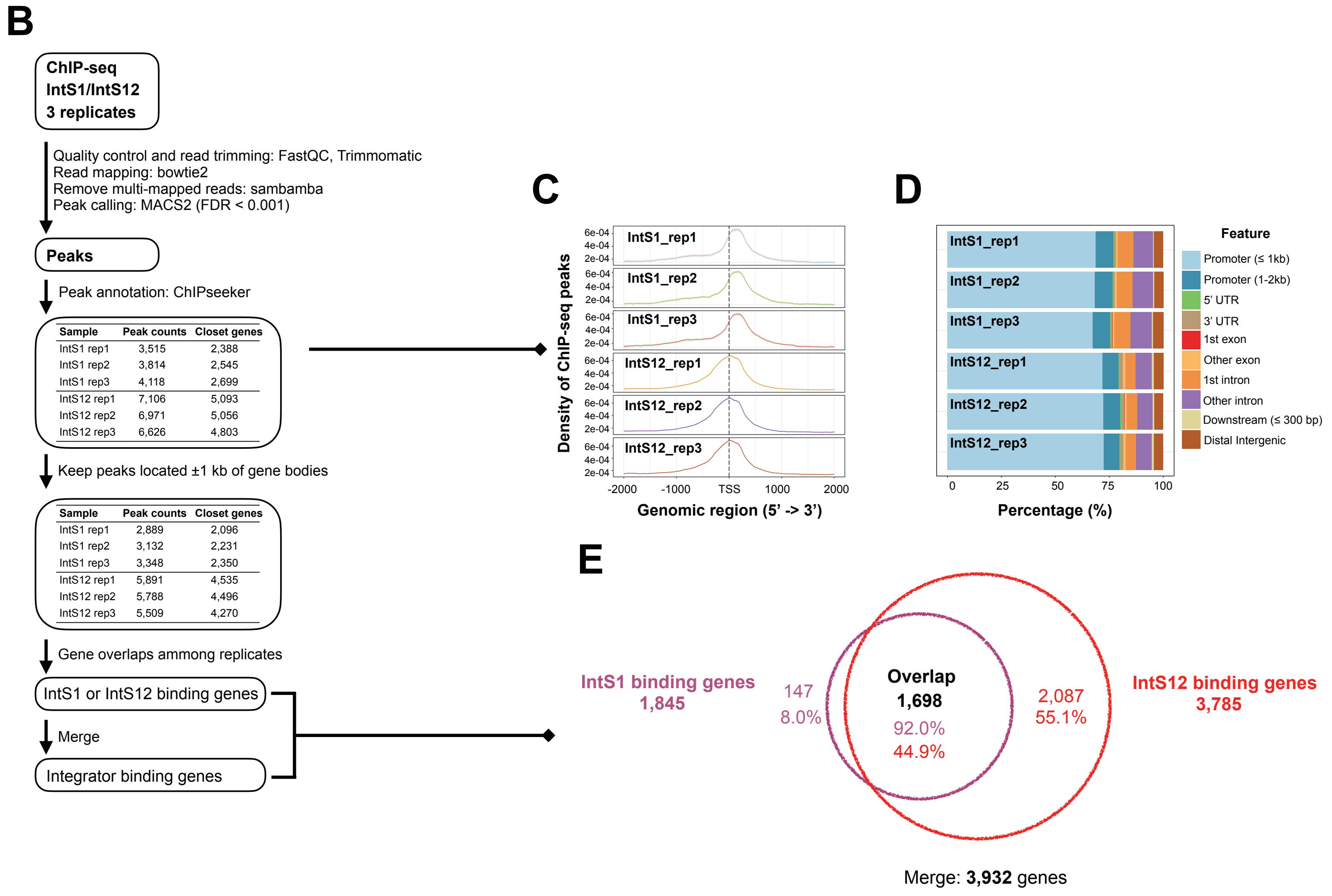

### Supplemental Figure S5

**A**

| Cell line | Treatment | Replicate | Total raw fragments | Total raw fragments after QC | Mappable fragments | % of total fragment after QC |
|---|---|---|---|---|---|---|
| DL1 | Mock | A | 22,171,785 | 21,678,505 | 19,931,033 | 91.94% |
| | β-gal | | 20,849,013 | 20,357,012 | 18,719,911 | 91.96% |
| | IntS4 dsRNA | | 20,791,117 | 20,304,747 | 18,580,892 | 91.51% |
| | IntS6 dsRNA | | 24,873,870 | 24,297,760 | 22,223,013 | 91.46% |
| | mts dsRNA | | 24,051,007 | 23,501,459 | 21,621,801 | 92.00% |
| | Pp2A-29B dsRNA | | 22,608,610 | 22,143,894 | 20,369,096 | 91.99% |
| | Mock | B | 24,145,833 | 23,570,455 | 21,599,574 | 91.64% |
| | β-gal | | 22,917,169 | 22,404,930 | 20,496,957 | 91.48% |
| | IntS4 dsRNA | | 26,735,089 | 26,104,104 | 23,762,636 | 91.03% |
| | IntS6 dsRNA | | 20,231,807 | 19,813,338 | 18,139,891 | 91.55% |
| | mts dsRNA | | 20,893,227 | 20,452,316 | 18,803,507 | 91.94% |
| | Pp2A-29B dsRNA | | 20,408,678 | 19,993,630 | 18,402,666 | 92.04% |
| | Mock | C | 22,026,825 | 21,584,441 | 19,825,005 | 91.85% |
| | β-gal | | 23,039,305 | 22,546,506 | 20,576,425 | 91.26% |
| | IntS4 dsRNA | | 20,078,746 | 19,631,208 | 17,825,607 | 90.80% |
| | IntS6 dsRNA | | 18,961,124 | 18,546,120 | 16,880,368 | 91.02% |
| | mts dsRNA | | 19,222,082 | 18,823,313 | 17,316,001 | 91.99% |
| | Pp2A-29B dsRNA | | 21,812,342 | 21,364,347 | 19,638,535 | 91.92% |

**B**

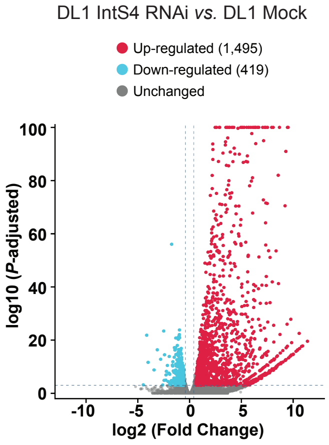


**C**

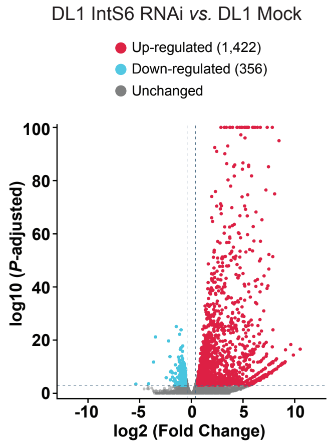


**D**

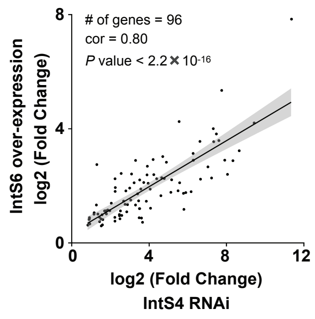


**E**

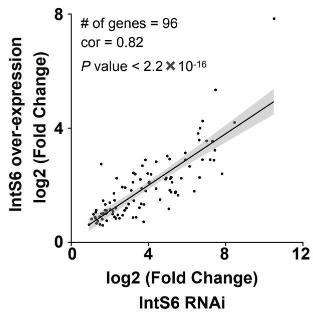


**F**

| Cell line | Treatment | Replicate | Total raw reads | Total raw reads after QC | Mappable reads | % of total reads after QC |
|---|---|---|---|---|---|---|
| DL1 | Control (β-gal) RNAi + eGFP | 1 | 23,529,728 | 23,393,483 | 21,925,101 | 93.72% |
| | | 2 | 33,850,956 | 33,658,251 | 31,435,322 | 93.40% |
| | | 3 | 12,033,584 | 11,965,598 | 11,173,495 | 93.38% |
| | IntS11 RNAi + eGFP | 1 | 15,877,946 | 15,788,918 | 14,553,790 | 92.18% |
| | | 2 | 16,463,448 | 16,371,234 | 15,212,661 | 92.92% |
| | | 3 | 17,537,551 | 17,434,328 | 16,165,080 | 92.72% |
| | IntS11 RNAi + IntS11 WT | 1 | 19,696,732 | 19,583,815 | 18,469,924 | 94.31% |
| | | 2 | 18,973,687 | 18,861,547 | 17,750,578 | 94.11% |
| | | 3 | 12,916,605 | 12,843,841 | 12,177,526 | 94.81% |
| | IntS11 RNAi + IntS11 E203Q | 1 | 22,505,894 | 22,379,294 | 20,941,029 | 93.57% |
| | | 2 | 21,725,224 | 21,601,751 | 20,482,375 | 94.82% |
| | | 3 | 13,422,683 | 13,353,135 | 12,552,111 | 94.00% |

**GSE144667**

**G**

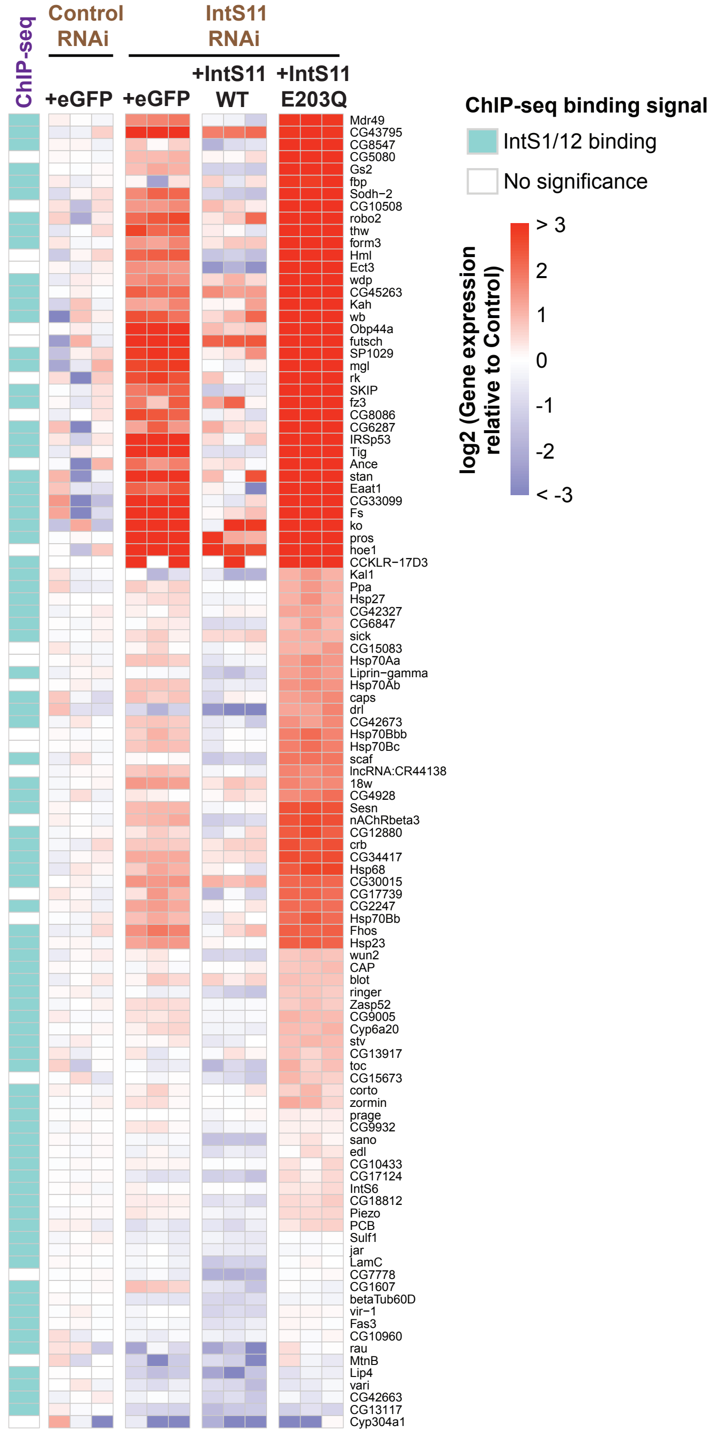

### Supplemental Figure S6

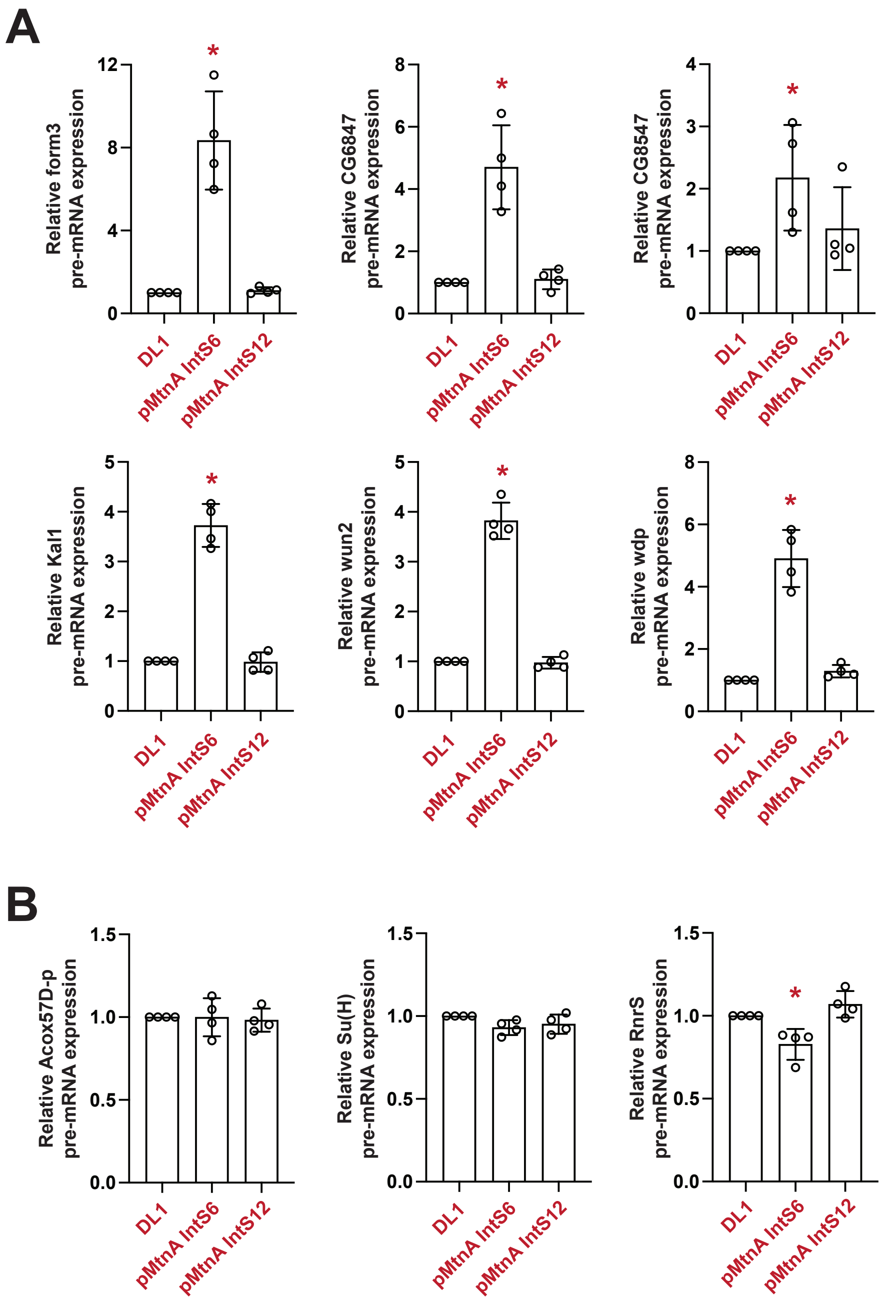

### Supplemental Figure S7

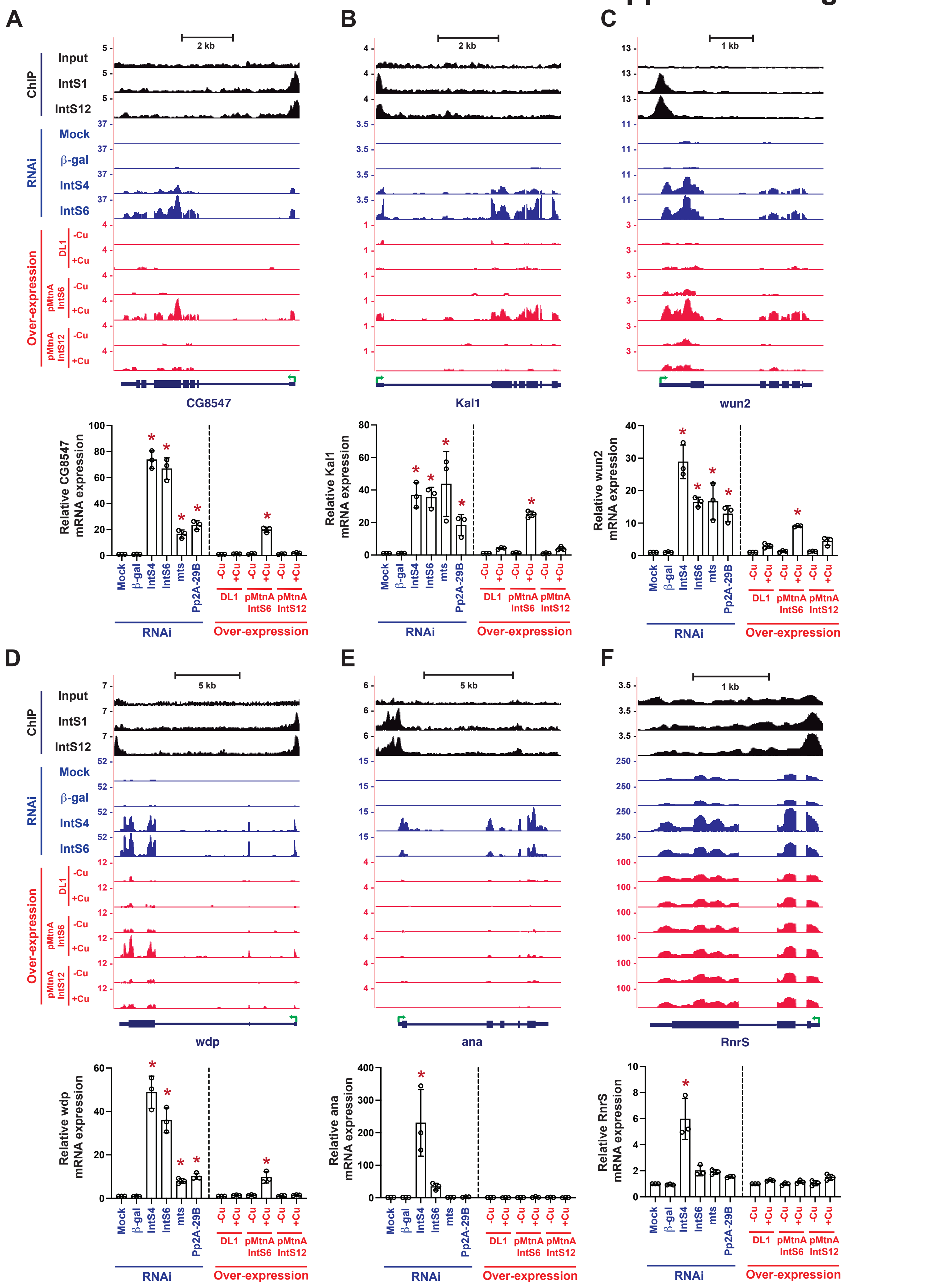
